# Convergent post-drought recovery of biomass and functional traits under constant and periodic warming in slow- and fast-growing plants

**DOI:** 10.64898/2026.02.01.703112

**Authors:** Nicolo Tartini, Ludovico Formenti, Yu Sun, Lauriane Bégué, Caroline Daniel, Itzel Lopez-Montoya, Gerard Martínez-De León, Nicholas O. E. Ofiti, Hang Zhao, Madhav P. Thakur

## Abstract

Extreme climate events such as droughts and heatwaves are intensifying under climate change, yet their combined effects on plant recovery remain unclear. In a two-year outdoor mesocosm experiment, we tested how grassland species with contrasting growth strategies recover from summer drought under four warming regimes: ambient, moderate warming (+2 °C), periodic heatwaves (+7 °C), and their combination. Experimental communities of native fast- and slow-growing species plus the invasive *Solidago canadensis* were assessed for above-ground biomass and leaf traits (SLA, LDMC, chlorophyll content, stomatal conductance) at one- and four-months post-drought. Biomass fully recovered within one month in both growth strategies, but leaf traits showed transient shifts, over-recovery in SLA and under-recovery in LDMC, likely reflecting production of new leaf tissues. These deviations generally returned to control levels by four months, regardless of warming treatments. *Solidago canadensis* exhibited high tolerance to heat and drought, with early biomass and trait recovery, indicating potential for dominance under climate extremes. Biomass recovery was similar across growth strategies, suggesting that growth-related differences play a minimal role in short-term recovery; however, early regrowth was characterised by contrasting trait shifts. Such lagged trait recovery, combined with rapid invasive recovery, suggests potential for longer-term shifts in grassland composition and function. We recommend that incorporating trait-based recovery dynamics is essential for predicting ecosystem stability under compound climate extremes.

## 1. Introduction

Over the past few decades, the magnitude and frequency of climate extremes, such as periodic heat waves and prolonged periods of droughts are increasing at an alarming rate (IPCC 2023; Gu *et al*. 2025). Periodic heatwaves and extreme droughts disrupt plant communities (Boeck *et al*. 2011; Smith *et al*. 2024), such as by negatively impacting biomass productivity, tissue quality, and altering species composition and diversity (Griffin-Nolan *et al*. 2019; Smith *et al*. 2024; Thakur *et al*. 2022). Such community-level shifts are ultimately shaped by how individual species respond to and recover from climate-induced stresses. Yet, we know little about how periodic heat waves (short, high-intensity temperature spikes) interact with constant warming (sustained temperature increases) to shape plant responses, particularly their post-drought recovery (Li *et al*. 2021; Breshears *et al*. 2021).

Most studies have shown that constant and moderate warming increase above-ground plant biomass while also simultaneously reducing their tissue quality (Jia *et al*. 2025; Lin *et al*. 2010; Liu *et al*. 2022; Wan *et al*. 2023; Wright *et al*. 2001; Zhou *et al*. 2022). In contrast, pulse events like heatwaves, that are shorter in duration but higher in intensity than constant warming, are typically associated with the loss in above-ground biomass productivity (Breshears *et al*. 2021; Wang *et al*. 2016), likely due to plant’s inability to overcome extreme heat stress (Breshears *et al*. 2021; López *et al*. 2022; Teskey *et al*. 2015). Drought alone is also consistently linked with declines in plant biomass (Yu *et al*. 2025; Zhou *et al*. 2020), whereas it has been shown that interactive effects of warming and drought on plant biomass are usually additive (Wilschut *et al*. 2022). Indeed, the combined effects of warming and drought likely vary across species, and can be effectively elucidated through trait-based approaches. For instance, species’ contrasting growth strategies—fast resource acquisition versus stress tolerance—may explain their differing responses to warming and drought (Reich 2014; Wright *et al*. 2004; Yan *et al*. 2025).

Drought recovery has been shown to differ between slow- and fast-growing species (Mentges *et al*. 2024; Pezner *et al*. 2020; Prugh *et al*. 2018; Stotz *et al*. 2022). Fast-growing plants are often less drought-resistant but typically recover quickly after re-wetting (Pezner *et al*. 2020; Zhou *et al*. 2022). Conversely, slow-growing plants tend to be more resistant to climatic stressors yet recover slowly, often due to lagged responses (Prugh *et al*. 2018; Reich 2014; Breshears *et al*. 2021; Luo *et al*. 2023a; Zwicke *et al*. 2015). Such differences can be assessed via key functional traits, including chlorophyll content, stomatal conductance, specific leaf area (SLA), and leaf dry matter content (LDMC) among others (Cornelissen *et al*. 2003; Poorter *et al*. 1995; Reich 2014; Wright *et al*. 2004). Drought generally reduces SLA and increases LDMC (Wellstein *et al*. 2017; Wilcox *et al*. 2021), while warming may increase SLA, potentially offsetting drought-induced declines within and across species. During post-drought recovery, fast-growing plants typically restore SLA and LDMC more rapidly than slow-growing plants (Chu & Farrell 2022; Boeck *et al*. 2018). Moreover, fast-growing plants also tend to lose chlorophyll quickly under stress but regain it rapidly upon rehydration, whereas many slow-growing plants maintain more stable chlorophyll levels (Chu & Farrell 2022; Luo *et al*. 2023b; Pezner *et al*. 2020; Wei *et al*. 2023; Zhang *et al*. 2021). Similar patterns occur for stomatal conductance: fast growers show sharp drought-induced declines but quick recovery, while slow growers modulate water use more effectively under drought (Li *et al*. 2021; Luo *et al*. 2023b; Valerio *et al*. 2022). However, the combined effects of drought and warming on these traits remain underexplored, especially regarding post-drought recovery in species with contrasting growth strategies. Emerging evidence suggests extreme heat can exacerbate drought impacts, further reducing SLA and causing prolonged physiological impairment (Tello-García *et al*. 2020; Wellstein *et al*. 2017), but it is unclear whether such compound stresses differentially constrain recovery within plant communities.

To better understand differences in post-drought recovery between slow- and fast-growing plant species, it is essential to consider recovery dynamics across the entire growing season, especially in seasonally structured ecosystems like temperate grasslands where climatic events may have temporally variable effects (Castillioni *et al*. 2022; Boeck *et al*. 2011; Sanders *et al*. 2025; Zhou *et al*. 2022). Yet, most climate-manipulation experiments focus on peak-season measurements (Stuble *et al*. 2021), potentially overlooking how recovery unfolds over time (Zhou *et al*. 2022). Slow-growing species may resist drought stress better but exhibit delayed recovery due to inherently conservative growth strategies, while fast-growing species, though more drought-sensitive, tend to recover quickly, often with pronounced trait shifts such as changes in SLA and LDMC (Figure 1). Moreover, any subsequent climatic events like heatwaves during late-season recovery may further modulate these trajectories, ultimately shaping species dominance and driving community reassembly under climate change (Figure 1).

**Figure 1.**
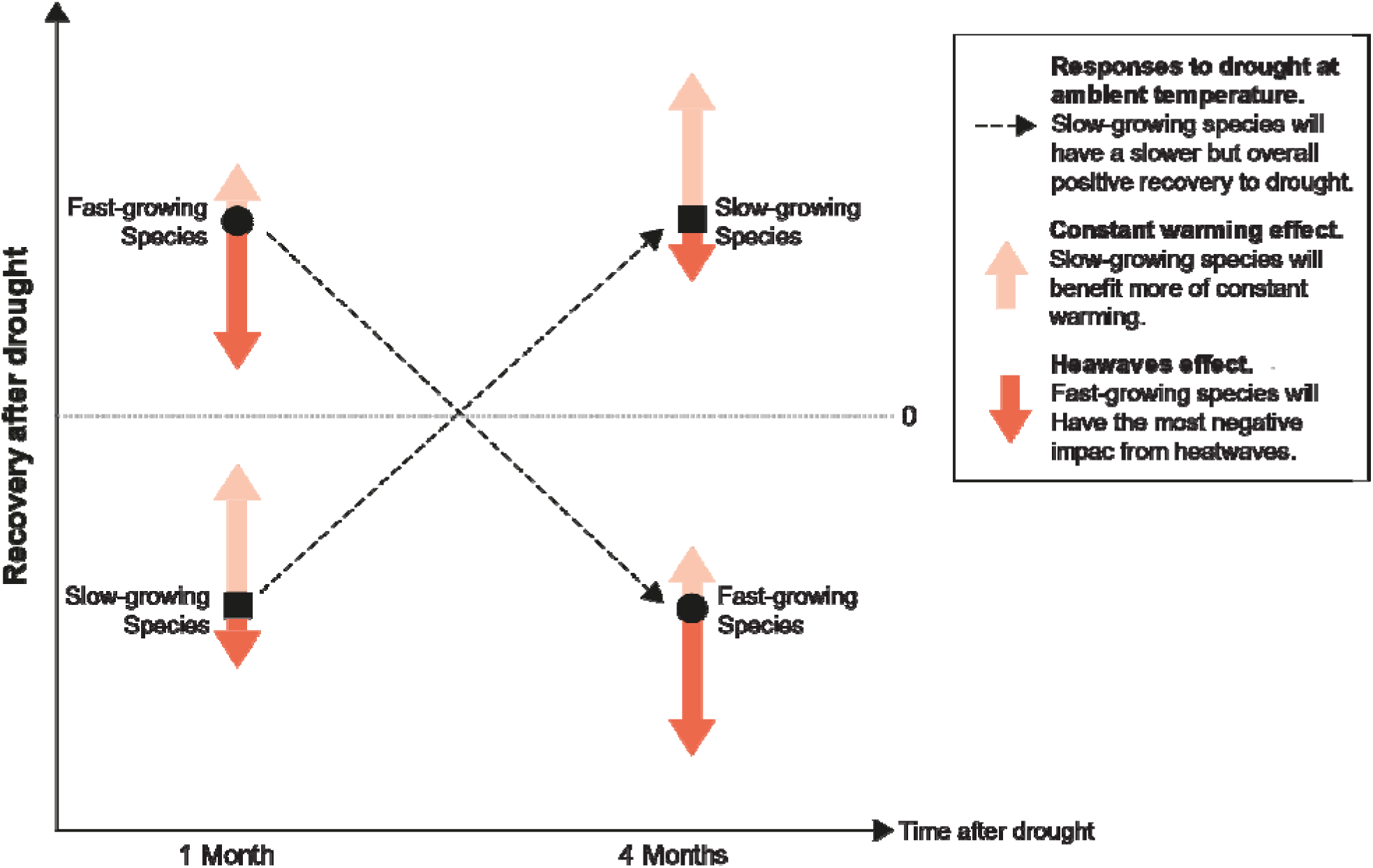
Key hypotheses related to the responses of slow- and fast-growing plants to drought under constant and periodic warming. Plant recovery (both biomass and traits) after drought is expected to depend on growth strategy. The one-month recovery of slow-growing species (squares) is predicted to be lower than that of fast-growing species, whereas their four-month recovery is expected to be higher. Fast-growing species (circles) are predicted to show the opposite pattern. Warming is hypothesized to modify post-drought recovery in both strategies: constant warming (orange arrows) is expected to enhance recovery at both stages, while periodic heat waves (dark red arrows) are expected to constrain it. The magnitude of warming effects is anticipated to differ between strategies—slow-growing species benefiting more from constant warming, and fast-growing species being more negatively affected by periodic heat waves.

We examined post-drought recovery in experimental grassland communities composed of species with contrasting growth strategies across two full growing seasons, using an outdoor experiment that combined extreme summer drought with either constant moderate warming, periodic heat waves, or both. Within this framework, we hypothesise that recovery dynamics would diverge between slow- and fast-growing species: fast growers would show rapid early recovery but decline over time, whereas slow growers would recover more gradually. We further predicted that warming regimes would modulate these trajectories, with constant warming facilitating recovery in both groups, while periodic heat waves would constrain it, particularly in fast growers (Figure 1). By testing these hypotheses, we assess whether gradual versus abrupt warming interacts with drought to reinforce or diminish functional differences in recovery between plant strategies.

## 2. Material and methods

### 2.1 Outdoor mesocosms

We established an outdoor experimental facility at the Hasli Ethological Station in Bern, Switzerland (46°58′01.362″N, 7°23′50.243″E; 490.6 m a.s.l.) to study post-drought recovery of plant communities under different warming regimes. The site has a mean annual air temperature of 9.3 °C and a mean annual precipitation of 1021 mm, based on 30-year reference averages MeteoSwiss (MeteoSwiss 2025). The facility is located in a managed meadow, and comprises the outdoor mesocosm units arranged across an area of ∼250 m² (Figure S1a).

The outdoor mesocosms were constructed from glass fiber–reinforced plastic (GRP), with inner dimensions of 1 × 1 × 0.45 m (L × W × H). Each had 20-mm insulated polyurethane (PUR) sandwich walls and a floor made of a porous polypropylene–polyester fiber matrix that allowed slow water drainage. Mesocosms were placed 1 m apart (Figure S1b; replication described under *Climate treatments*). Each mesocosm was filled with a customized artificial soil mixture (RICOTER AG, Switzerland) replicating the physicochemical properties of mineral soil from a nearby managed meadow. The mixture comprised 40% gravel (0–32 mm), 40% vegetal earth (0–15 mm), 10% sand (0–2 mm), and 10% garden compost, forming a ∼35 cm mineral layer. On top, we added a 5 cm layer of topsoil rich in organic matter, obtained by collecting the upper 10 cm of soil from the same meadow. The soil was sieved through a 1 cm motorized mesh (Scheppach RS 350; Scheppach GmbH, Ichenhausen, Germany) and thoroughly homogenized by hand before being added to the mesocosms. Soil properties (pH, organic matter content, and organic and inorganic C and N) were measured for the mineral- and the topsoil (Table S1).

### 2.2 Plant community establishment

In July 2022, each mesocosm was planted with an identical community of eight common grassland species characteristic of extensively managed central European meadows. Species were selected based on their position along the leaf economic spectrum (Wright *et al*. 2004), using specific leaf area (SLA) to represent both fast- and slow-growing strategies. To ensure phylogenetic balance, we included four plant families, each represented by one slow- and one fast-growing species: *Lotus corniculatus* (Fabaceae, slow) and *Trifolium pratense* (Fabaceae, fast); *Centaurea jacea* (Asteraceae, slow) and *Taraxacum officinale* (Asteraceae, fast); *Bromus erectus* (Poaceae, slow) and *Holcus lanatus* (Poaceae, fast); and *Salvia pratensis* (Lamiaceae, slow) and *Prunella vulgaris* (Lamiaceae, fast) (Figure 2c). Growth strategies were classified relative to other species within the same family. To better reflect managed grasslands of central Europe (e.g., Palearctic meadows), we also included the non-native invasive *Solidago canadensis*, which is one most common invasive plants in central Europe (Nentwig *et al*. 2018).

**Figure 2.**
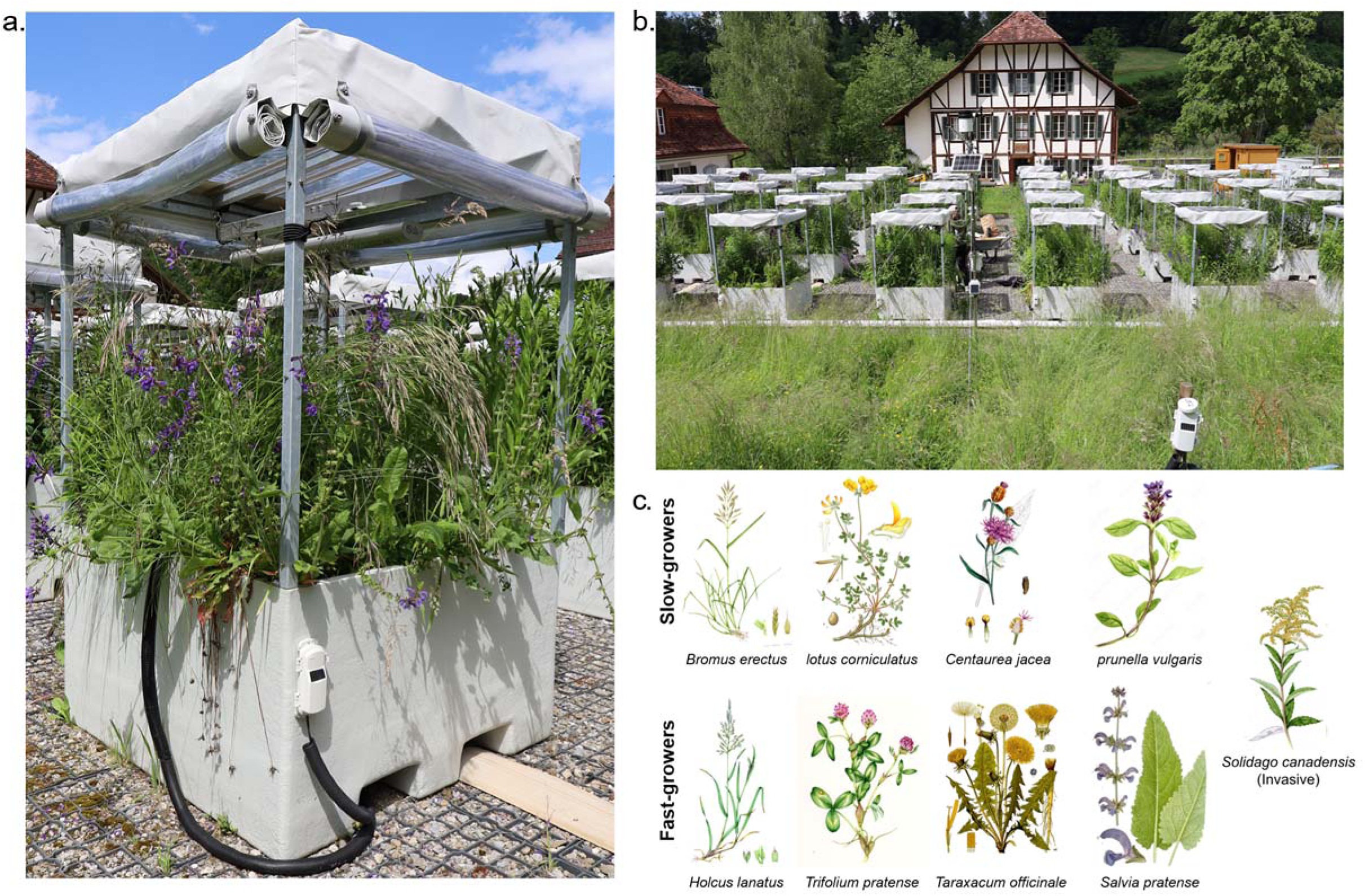
(a) Experiment setup mesocosm at the onset of the drought. (b) mesocosm with the meteorolo gicalstation installed at the Haslisite. (c) Illustration of the nine study species arranged by growth strategy; *Solidago canadensis* was excluded from the fast–slow classification due to its invasive status in Europe.

Seeds of all species were germinated indoors for three weeks in trays with germination soil (LANDI, Capito line) at 20/16 °C (16 h day/8 h night) before being transplanted into a standardized randomized grid applied uniformly across all mesocosms (Figure S2).

Each mesocosm received 12 individuals per species (108 plants in total). From planting until the onset of experimental treatments in March 2023, dead seedlings were regularly replaced. However, *Prunella vulgaris* failed to establish in 2022, and was therefore reintroduced during the experimental phase in spring–summer 2023.

In mid-November 2022, after plant communities had established but before the application of climatic treatments, baseline data were collected from each mesocosm (Figure S3). These data included total plant dry biomass, specific leaf area (SLA) and leaf dry matter content (LDMC). We measured specific leaf area (SLA; mm² mg⁻¹) on young, fully expanded leaves after 24 h of water saturation on moist paper at 4 °C. Leaf area was digitized using an Epson Perfection V600 Photo Scanner (Epson Corp., Nagano, Japan) (one leaf per species per mesocosm, except *L. corniculatus*, for which two leaves were sampled due to their small leaf size) and divided by leaf dry mass (dried at 40 °C for 48 h). Leaf dry matter content (LDMC; mg g⁻¹) was calculated as the ratio of oven-dry mass to water-saturated fresh mass (Cornelissen *et al*. 2003). Once fully dried (≥ 70 °C for several days), the plant material collected from each mesocosm was finely shredded and returned as litter to the same mesocosm in March 2023 to minimize nutrient loss from the system. The same procedure was repeated for plant material collected in 2023 and 2024 (details below).

### 2.3 Climate treatments

Climate treatments consisted of four warming regimes (‘Ambient temperature’, ‘Ambient + 2 °C’, ‘Ambient + periodic heat waves’, and ‘Ambient + 2 °C + periodic heat waves’) fully crossed with two precipitation regimes (control water and drought). Each treatment combination was replicated five times and randomly assigned to mesocosms (Figure S1b). The 40 mesocosms were arranged in five blocks to account for wind and light variability, with each block containing one replicate of all eight climate treatments.

Warming was applied using infrared heaters (VASNER SlimLine X20, 500–2000 W, Germany) mounted 80 cm above the soil surface of each mesocosm (Figure S4); in the ambient treatment, fake heaters were installed as controls at the same height. Air temperature (15 cm above the soil) in each mesocosm was continuously monitored with HOBO sensors (MX2201, Oneset Computer Corp., Bourne, USA), and a weather station (Oneset Computer Corp., Bourne, USA) was installed adjacent to our mesocosms to record ambient air temperature, relative humidity, solar radiation, and rainfall (2 m above the soil) (Figure S1).

The ‘Ambient +2 °C’ treatment increased air temperature by ∼2 °C above ambient (15 cm above the soil) throughout the growing season (mid-March to late October), starting in 2023 and repeated in 2024 (Figure S5). The ‘Ambient + periodic heat waves’ treatment raised air temperature by ∼7 °C at the same height for one week each month, producing seven heat waves per season (Figures S4 & S6). The ‘Ambient +2 °C + periodic heat waves’ treatment combined both approaches, maintaining a +2 °C temperature throughout the growing season while adding monthly heat waves, during each heat wave week (Figure S5). The ‘Ambient +2 °C’ treatment reflects the projected global mean temperature rise by the mid-21^st^ century under high-emission scenarios, whereas our periodic heat-wave treatments simulate regional extremes. Heat wave treatments are consistent with IPCC AR6 projections that the hottest days can warm 1.5–3 times more than the global average—resulting in anomalies approaching +6 to 10°C for one week each month (Figures S4-S6) (IPCC 2023).

The drought treatment consisted of a single extreme drought event applied each experimental year (2023 and 2024), beginning in late May until the end of June. By the end of the drought period, many plants in our experiment showed severe wilting and withering suggesting that our drought regimes were representative of both hydrological and ecological extreme drought (∼5–6 weeks of complete water deprivation; Figure S7). This period was chosen to simulate summer drought, which has increased in frequency across northern Europe, including in our study region (IPCC 2023). At the end of each simulated drought, soil water content (0–10 cm depth) declined to 0–5%, also corresponding to an “extreme drought” category according to the U.S. Drought Monitor (National Drought Mitigation Center, USDA, and NOAA; Figure S8). To maintain drought conditions and exclude confounding rainfall, all mesocosms (drought and non-drought) were covered with transparent plastic roofs during the drought period. We measured potential shading effects with a HOBO MX2202 data logger (Onset Computer Corp., Bourne, USA) equipped with a light sensor positioned near the plant canopy, which recorded a mean reduction of ∼25% in light intensity relative to the same period at a nearby climate station (Figure S9). Water control mesocosms received tap water (up to twice weekly) during the drought period according to 30-year average rainfall data (MeteoSwiss 2025), using a hose with a volume counter. Outside the experimental drought period, all mesocosms were regularly watered with tap water according to the 30-year average rainfall, particularly during prolonged periods of no or insufficient precipitation as indicated by our on-site weather station. Soil water content was regularly monitored before, during, and after drought using both (i) a TDR probe (FieldScout TDR 150 Soil Moisture Meter, Spectrum Technologies Inc., Aurora, USA) for specific time point measurements per growing-season at 10 cm depth (Figure S8), and (ii) continuous sensors (GroPoint, Onset Computer Corp., Bourne, USA) at 0–15 cm and 15–30 cm depths (Figures S10–S11).

### 2.4 Plant data collection (2023 and 2024)

Species-specific plant data were collected prior to drought (early growing season, mid–late May), one month after drought (mid-July), and four months after drought (mid-October). To avoid re-sampling the same plant material within the same year in each mesocosm, different quarters (0.5 x 0.5 m) were randomly selected each in 2023 and 2024. Most plant-specific measurements followed the 2022 protocol, with two exceptions. First, chlorophyll content was measured with a Force-A Dualex® meter (Force-A, Orsay, France). At the end of the 2024 growing season, however, we used a SPAD meter (Konica Minolta, Osaka, Japan) due to the unavailability of the Dualex®. Second, we measured stomatal conductance using an SC-1 leaf porometer (Decagon Devices Inc., USA), excluding values above 500 mmol m⁻² s⁻¹ because they exceed the instrument’s precision range. To further illustrate the extremity of the drought treatment, we also report stomatal conductance of each species immediately after drought, before rewetting (Figure S12). Plant recovery after drought (at one and four months) was quantified for all nine species across the four warming regimes, using measurements of shoot biomass, SLA, LDMC, chlorophyll content, and stomatal conductance.

### 2.5 Statistical analysis

Post-drought plant recovery across warming regimes was analysed separately for all response variables using R v.4.3.3 (R Core Team 2023). Mixed-effects models were fitted with the package glmmTMB v.r1.1.9 (Brooks, E., Mollie *et al*. 2017). All models included the interactive effects of drought, temperature, season, and either plant growing strategies or species (see details below). Our analyses focused on post-drought responses during the peak of the growing season (one-month recovery) and the end of the growing season (four-months recovery). Since plants in spring 2023 had not yet experienced any drought, pre-drought responses (early growing season) of 2023 were not used in our mixed-effects models. Responses in spring 2024 are used in mixed-effects models as they likely reflect the legacy effects of the 2023 drought (Figure S13). Accordingly, spring 2024 data were included in the mixed-effects models (responses shown in Figure S14, Tables S2-S11). Post-drought recovery was quantified as effect sizes from the mixed-effects models.

All models included plots (to account for the repeated measurements) and year (two levels) as independent random effects. We applied two main modelling approaches. First, we tested whether slow-versus fast-growth strategies explained recovery across eight native species, with species included as a random effect to capture variation beyond growth strategy (Tables S2-S6). We considered two main growing strategies: native slow- and fast-growing species, whereas invasive species (*S. canadensis*) was used as another level. Growing all species in a shared experimental community minimised environmental variability and strengthened trait-based inference. Second, we analysed species-specific responses by treating species as fixed effects in models that excluded growth strategy (Tables S7-S11). Models for *Prunella vulgaris* were analysed separately due to their poor establishment in 2022 and 2023.

To ensure optimal model fit for each response variable, we applied appropriate statistical distributions and transformations. Biomass was modelled with a negative binomial distribution, except for *P. vulgaris*, where a Gaussian distribution was used. SLA and LDMC were modelled with Gaussian distributions, whereas for *P. vulgaris,* a negative binomial distribution was more appropriate. Chlorophyll content was modelled with a negative binomial distribution. Stomatal conductance was log-transformed and modelled with a Gaussian distribution, except for *P. vulgaris*, where it was again fitted with a negative binomial distribution. We tested the fit of all mixed-effects models, checking the models’ residuals using the package DHARMa v.04.7 (Hartig 2016). The model assumptions were generally met, except the model for the above-ground biomass of *P. vulgaris*, which showed a poor fit (across different error distributions), and therefore, those results should be interpreted cautiously.

After fitting mixed-effects models for each response variable, we calculated drought effect sizes in all four warming treatments, using the packages emmeans v1.10.1(Lenth 2017) and effsize v0.8.1 (Torchiano 2013). Full model outputs and effect sizes are provided in the Supplementary Information (Tables S2-S11). Post-drought recovery at one and four months after drought was visualised as Cohen’s d with 95% confidence intervals (CIs) across all warming regimes. Effect sizes were plotted using the ggplot2 package (Wickham 2016). Recovery outcomes were classified as full recovery (non-significant, CI overlapping zero), under-recovery (significant negative effect), or over-recovery (significant positive effect).

## 3. Results

### 3.1 Growth-strategy-specific recovery

Full recovery of above-ground biomass in slow- and fast growing plants was evident already one month after drought across all warming treatments (Figure 3), despite clear signs of severe stress and wilting, including altered stomatal conductance recorded immediately post-drought (Figures S7, S12). This full biomass recovery persisted four months after drought in both growth strategies and across warming treatments (Figure 3).

**Figure 3.**
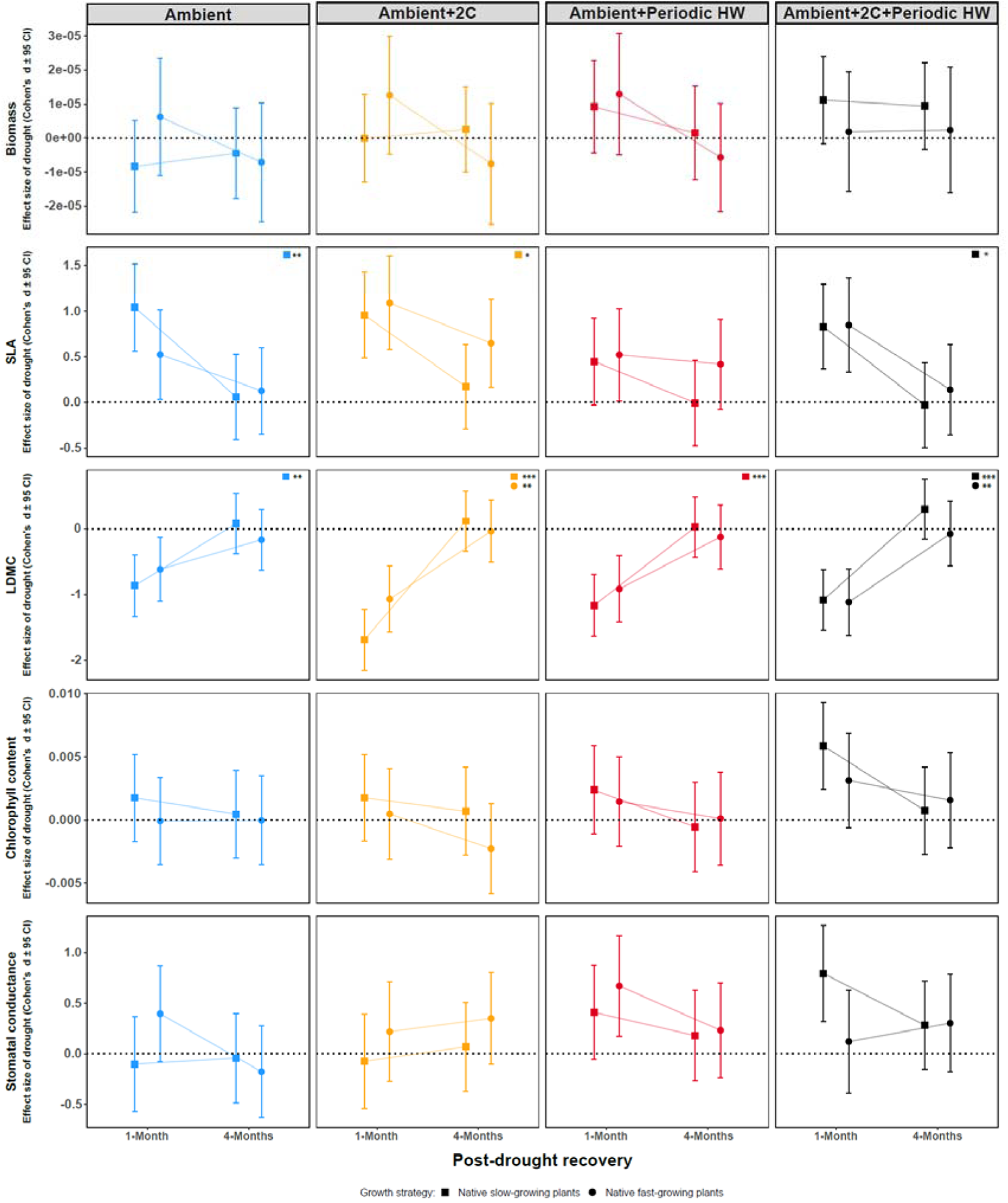
Post-drought recovery of slow- and fast-growing native plants one and four months after drought under four temperature regimes. Recovery is shown for shoot biomass, specific leaf area (SLA), leaf dry matter content (LDMC), chlorophyll content, and stomatal conductance, based on data from 2023 and 2024 (see Methods). Values represent Cohen’s d ± 95% confidence intervals (CIs); recovery is significant when CIs do not overlap zero. Significant differences between effect sizes of the two recovery periods within each growth strategy were assessed using estimated marginal means from mixed-effects models with Tukey-adjusted pairwise comparisons (p < 0.05; *p < 0.01; **p < 0.001). HW denotes heat waves.

By contrast, we found pronounced shifts in SLA and LDMC one month after drought across warming regimes. SLA showed significant over-recovery, increasing on average by 26.1% in slow-growing species and 18.6% in fast-growing species compared with water controls. Reciprocally, LDMC showed significant under-recovery, with decreases of 19.4% in slow-growing species and 17.5% in fast-growing species. Differences between growth strategies were not statistically significant for either of the traits (Table S2-S6). By four months after drought, both SLA and LDMC fully recovered across warming regimes and growth strategies, with the sole exception of SLA in fast-growing species under constant warming, which remained above control levels (Figure 3).

### 3.2 Species-specific recovery

All eight native species almost fully recovered in biomass and functional traits four months after drought; however, distinct differences in recovery emerged more clearly one month after drought, particularly in the warmed treatments (Figure 4). Interestingly, *T. pratense* was the only species to show under-recovery in above-ground biomass four months post-drought under constant warming, despite its earlier over-recovery in the same treatment just one month after drought (Figure 4). Indeed, many species (*B. erectus, T. pratense, T. officinale, and P. vulgaris*) showed the pattern of over-recovery one month after drought, and periodic heat waves contributed to this pattern in *B. erectus, T. pratense, T. officinale*.

**Figure 4.**
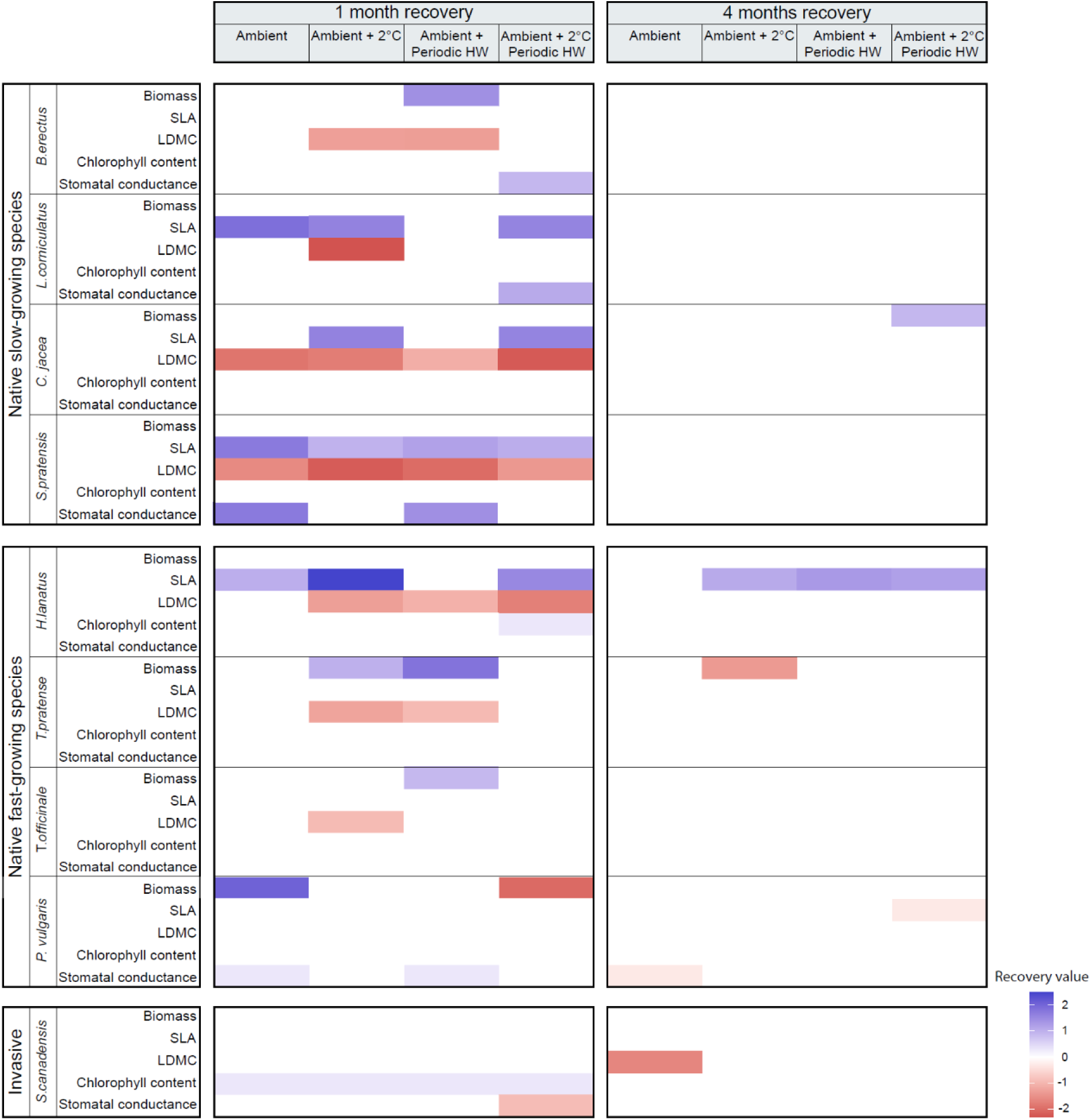
Overview of post-drought recovery (one month and four months after drought) across plant species under four temperature regimes. Recovery values are expressed as effect sizes (Cohen’s d). Negative values indicate under-recovery (negative drought effects; confidence intervals not overlapping zero), values around zero indicate full recovery (no difference between drought and non-drought treatments; confidence intervals overlapping zero), and positive values indicate over-recovery (positive drought effects; confidence intervals not overlapping zero). The effect of warming in spring 2024 (pre-drought) is shown in Figure S13.

Traits related to photosynthesis and gas exchange, namely chlorophyll content and stomatal conductance, also reached full-recovery in most species after one month of drought across warming regimes (Figure 4). By contrast, traits related to leaf tissue quality (SLA, LDMC) showed over- or under-recovery in many species one month after drought, and even persisted in a few cases four months (e.g., *H. lanatus*) after the extreme drought event. The LDMC in *B. erectus, C. jacea, S. pratensis, H. lanatus, T. prantense* was under-recovering one month after drought, and notably so in constantly warmed treatments (Figure 4). The combined constant and periodic heat wave treatments (‘Ambient + 2°C + periodic heat waves’) further amplified the under-recovery in *C. jacea* at this time point of post-drought recovery. Interestingly, *C. jacea* showed over-recovery in its biomass four months after drought under the same intensified warming regime (Figure 4). In addition, many species showed an inverse recovery pattern between SLA and LDMC, with SLA frequently over-recovering in cases where LDMC was under-recovering (Figure 4). Most cases of SLA over-recovery one month after drought converged to full recovery by four months, except for *H. lanatus*, where SLA values remained high under both constant warming and periodic heatwave treatments under both constant warming and periodic heatwave treatments.

The invasive species *S. canadensis* in our experimental plant community was largely tolerant to drought and showed a rapid recovery to both drought and warming, with a few notable exceptions. For instance, it showed a significant decline (under-recovery) in LDMC four months after drought, despite no change one month post-drought (Figure 4). Moreover, its stomatal conductance exhibited under-recovery one month after drought under the combined constant warming and periodic heatwave treatment (Figure 4).

## 4. Discussion

Using an outdoor experiment over two consecutive growing seasons, we found that several common temperate grassland species achieved full post-drought recovery under both constant warming and periodic heatwave conditions. More strikingly, by four months after extreme drought—coinciding with the end of the growing season—both slow- and fast-growing species reached full recovery, regardless of warming treatment and even sharing the same recovery trajectories in both one month and four months post drought data points. These findings contradict our hypothesis, which predicted a delayed recovery in slow-growing species and a faster recovery in fast-growing species (Figure 1).

Both fast- and slow-growing species recovered their biomass at similar rates under all warming regimes, suggesting that the severity of the drought was so intense that it forced plants to pursue the same recovery mechanisms, equalizing their responses. We suspect that extreme water limitation may have created a threshold condition, forcing plants into a common “stasis” response characterized by halted growth and canopy dieback (Breshears *et al*. 2021). Once rewetting occurred-timed with the peak of the growing season (e.g., July), when light and temperature conditions are optimal-plants rapidly regenerated above-ground tissues to ensure their end of the growing season survival (Zhou *et al*. 2022). This convergence in strategy, rather than the expected divergence based on growth rate (Jia *et al*. 2025; Stuble *et al*. 2021), highlights the adaptive capacity of grassland species to rebound from compound stress events, and aligns with recent evidence that phenotypic plasticity under extreme events may buffer against anticipated trade-offs (Wan *et al*. 2023).

Despite the rapid biomass recovery, leaf trait recovery suggests a more nuanced adaptive strategy of plants under drought and warming. Most species exhibited over-recovery in specific leaf area (SLA) and under-recovery in leaf dry matter content (LDMC) one month after drought, indicating the production of softer, less lignified foliage—traits typically associated with rapid regrowth rather than full functional recovery (Künzi *et al*. 2025; Wellstein *et al*. 2017). These patterns likely reflect the presence of young, newly formed tissues that had not yet matured structurally. By four months post-drought, however, most species exhibited full recovery in both SLA and LDMC, suggesting that trait-based recovery may lag behind biomass recovery. These findings underline the importance of temporal scale in interpreting the stability of plant species: apparent recovery at one functional level (biomass) may mask delays or vulnerabilities in other dimensions (tissue structure), with potential consequences for herbivory risk, carbon cycling, and overwintering success.

The invasive species *Solidago canadensis* showed a notable exception in our study. While largely unresponsive in biomass across treatments, *S. canadensis* exhibited under-recovery in LDMC four months post-drought, and its stomatal conductance was reduced one month after drought under the intensified warming treatment (‘Ambient + 2C + periodic heat waves’). This response is really striking if we compare it with the native species, which under the same warming condition showed a neutral or an increase in gas exchanges. The negative effect on stomatal conductance of *S. canadensis* could be a sign of a water parsimony strategy. Remarkably, *S. canadensis* partially retained above-ground foliage during drought when most native species had wilted (Figures S7 & S12), possibly enabling faster regrowth from residual tissues or rhizome reserves. This pattern of resistance and rapid recovery may provide *S. canadensis* with a competitive advantage in increasingly extreme climatic conditions, reinforcing concerns that climate change may amplify invasive species dominance in temperate grasslands (Prugh *et al*. 2018; Sanders *et al*. 2025). In line with this, our observation that *S. canadensis* has increased its biomass dominance over two years in all control mesocosms (Figures S3 & S14) further suggests that such traits may shift community trajectories over time, particularly when coupled with ecosystem disturbances (Griffin-Nolan *et al*. 2019; Luo *et al*. 2023b).

Taken together, our findings highlight the strong stability of temperate grassland species to extreme drought events even under warming, particularly when conditions for regrowth are favorable. The rapid recovery of biomass and, to a slightly delayed extent, leaf traits, suggests that grassland ecosystems may retain functional resilience under recurrent climate extremes-at least in the short term, as our results are based on two years of observation. However, this does not imply that grasslands remain unaffected: our mesocosms represent a young, still-establishing community dominated by an invasive species, and results may change as biotic interactions may lead to ecological debts under harsh and repeated abiotic conditions (Martínez-De León & Thakur 2024; Thakur *et al*. 2022). Furthermore, we have focused here on the above-ground plant responses. Yet, post-drought dynamics of below-ground responses, such as root biomass and traits, soil microbes and invertebrate communities, remain poorly understood and could feedback on longer-term ecosystem stability. Continued long-term monitoring is therefore essential to capture delayed effects, including potential shifts in species dominance, loss of diversity, and biogeochemical feedbacks that may only become evident over extended timescales.

## Acknowledgement

We thank Bruno Kampf and Erich Fuhrer from the Betrieb & Technik department of the University of Bern for their support in establishing the outdoor mesocosm experiment. We are also grateful to all current and former members of the Terrestrial Ecology Group and the Institute of Ecology & Evolution for their help in setting up the experiment in 2022. Figure 1 was drawn by Nadine Kamber (https://www.definitivdesign.ch/). We further thank Philippe Graber, Anouk Gremion, Loraine Hablützel, Arianne Marty, Dennis Loosli, Johan-Jakob Schindler, Felix Rentschler, Anine Wyser, and Roem Yesil for their assistance with data collection. We acknowledge the use of Open AI ChatGPT (versions 4 and 5) for the language and coding refinements. This project was funded by the Swiss State Secretariat for Education, Research and Innovation (SERI) under contract number M822.00029.

## Author contribution

MPT conceived the study. LF and MPT designed the experiment. NT, LF, and YS collected the data, with contributions from all co-authors. NT analysed the data with substantial input from LF, CD, GMDL, NOEO and MPT. NT, LF, and MPT wrote the first draft of the manuscript. All authors contributed to revising the manuscript.

## Notes

### Competing Interest Statement

The authors have declared no competing interest.

