## Supplementary figures 1-14, Supplementary tables 1-11 for "Convergent post-drought recovery of biomass and functional traits under constant and periodic warming in slow- and fast-growing plants"

^#^ shared authors

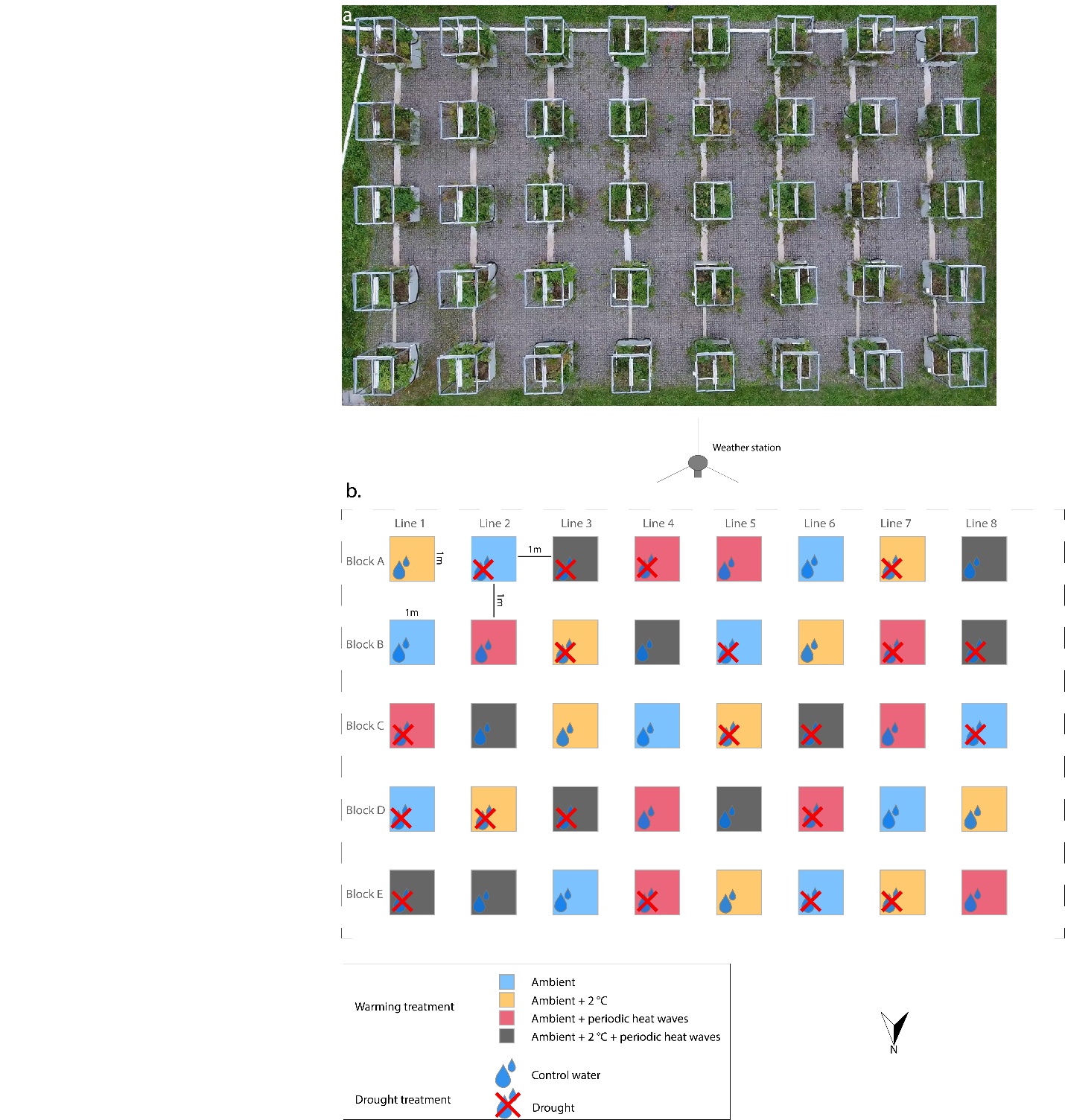

**Figure S1. a)** Aerial view of the forty mesocosms (~250 m²). **b)** Schematic representation of the randomized treatment assignment. Each block contains all eight combinations of warming and drought treatments, which were randomly allocated to the experimental units within each of the five blocks.

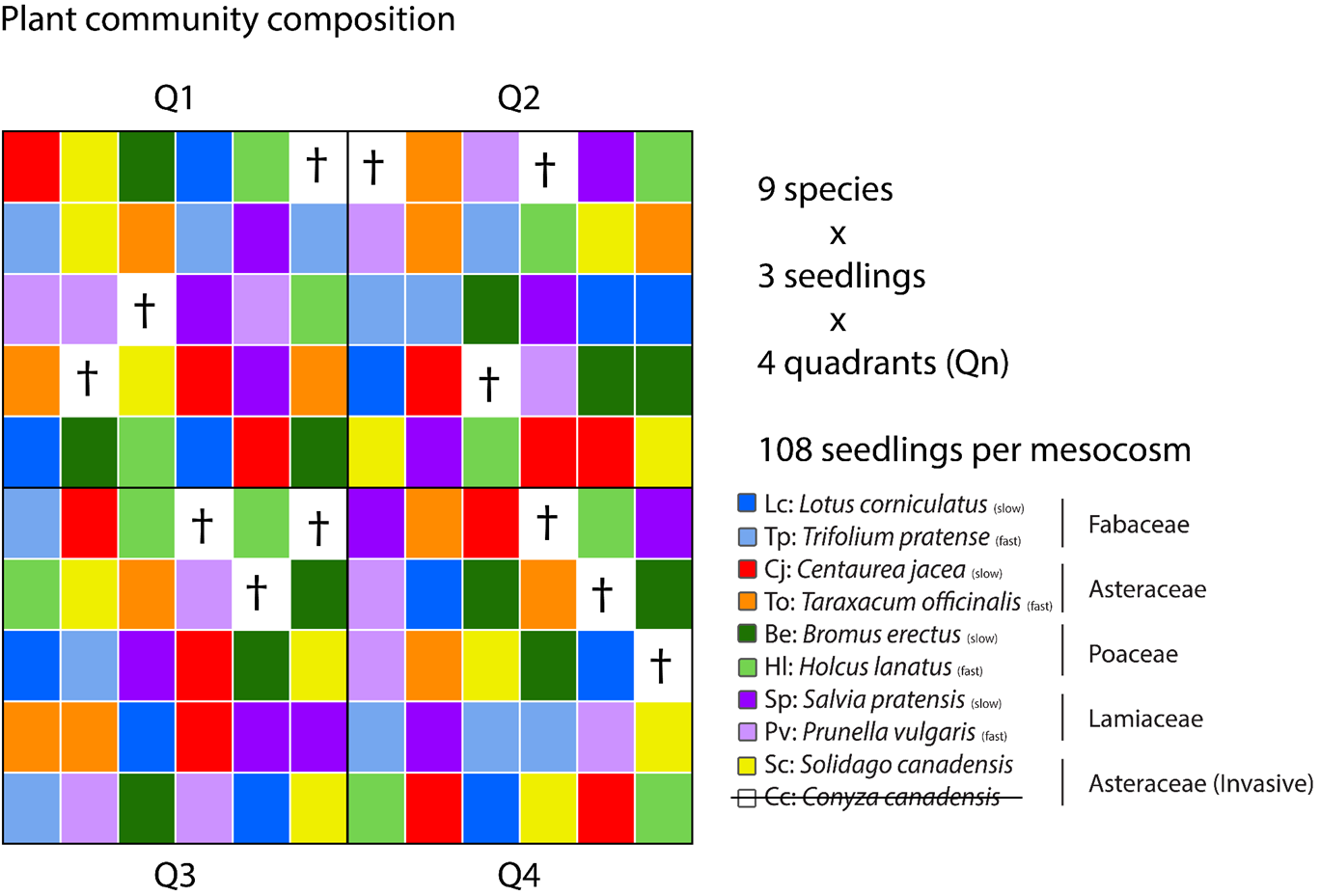

**Figure S2.** Schematic showing the initial planting arrangement of 12 individuals per species (represented by different colors) within each mesocosm. The same layout was used across all mesocosms. Crosses on white squares indicate failed establishment of *Conyza canadensis* in 2022.

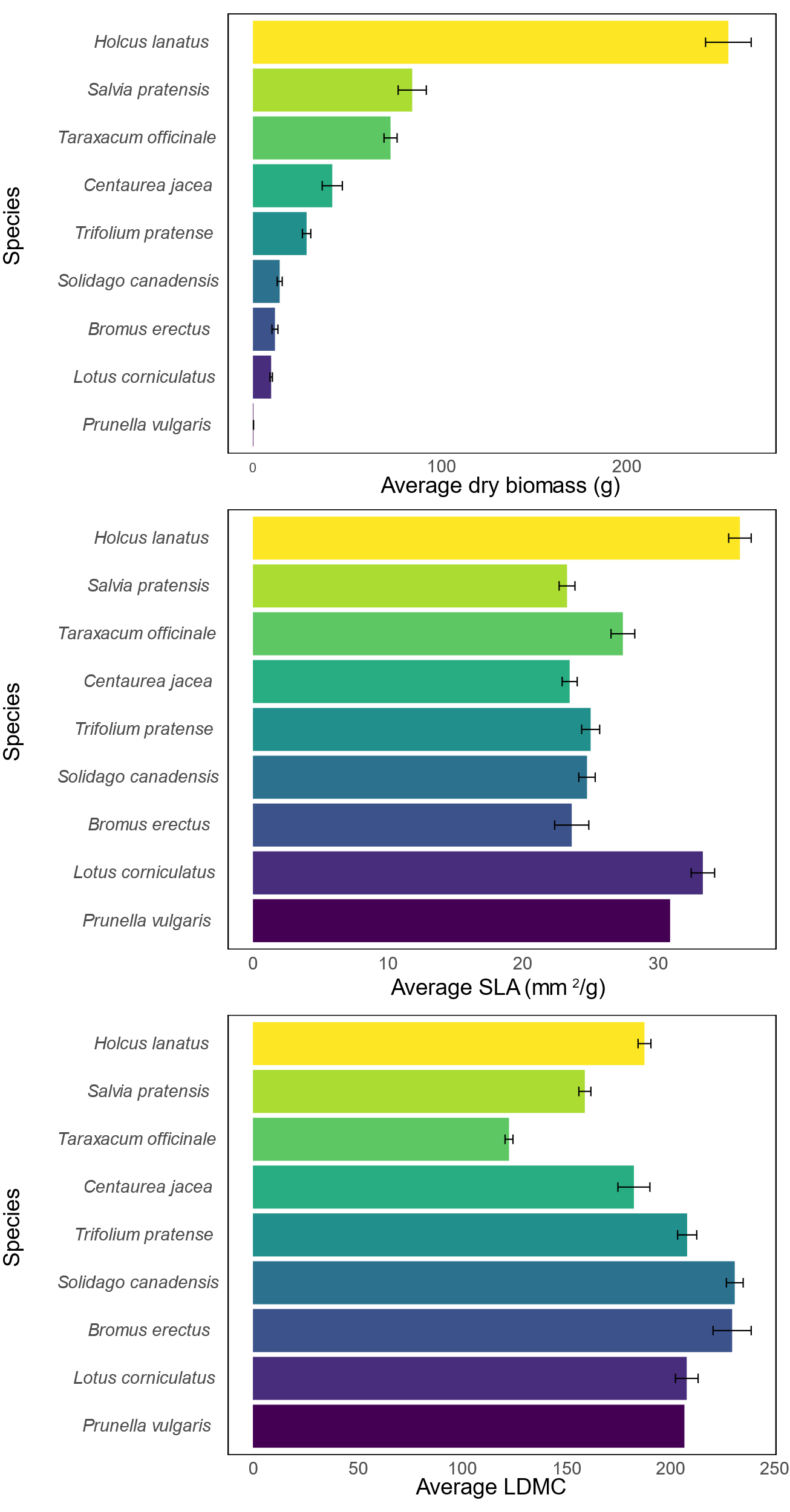

**Figure S3.** Baseline plant specific biomass (a) and traits (b, c, d) measured at the end of the growing season in 2022, a year before the climate treatments were applied. Please note that *Prunella vulgaris* did not establish in the year 2022.

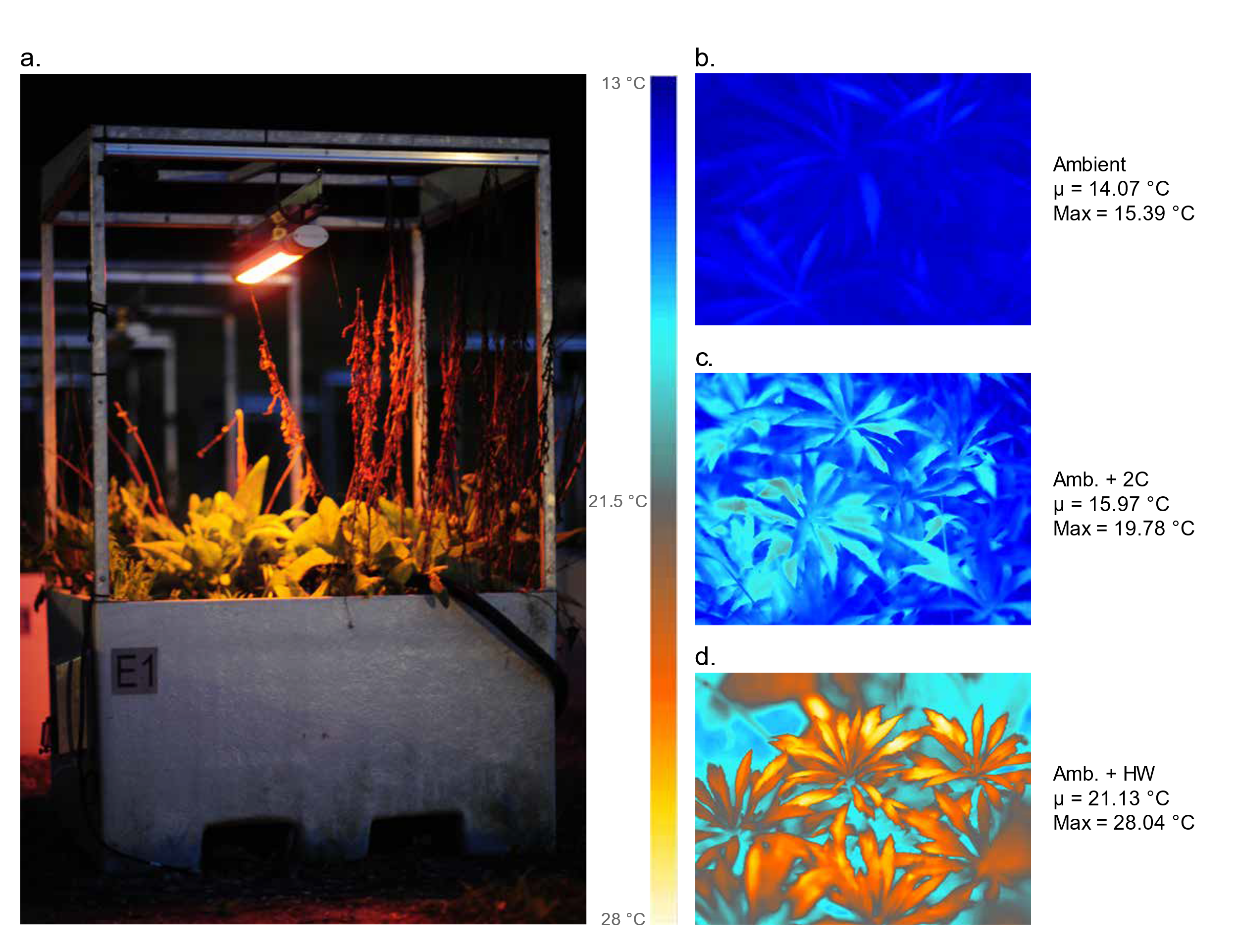

**Figure S4.** Example of the infrared heater setup. Heaters were manually calibrated to generate either constant warming or periodic heat waves. For illustration, the heater in the image was digitally enhanced to show heating; under experimental conditions, it emits negligible visible light at night. Thermal images (b-d) show warmed leaves of *Solidago canadensis* approximately 25 cm above the soil surface, taken during a heatwave week. In the thermal images, “µ” denotes mean temperature and “Max” denotes maximum temperature. During heat waves, canopy temperatures in the Amb + HW treatment reached up to 12 °C above those in the ambient warming treatment, with mean canopy temperatures up to ~7 °C higher, consistent with temperature readings from loggers placed 15 cm above the soil surface (see Fig. S6), where canopy height was lower (e.g., at the start of the growing season).

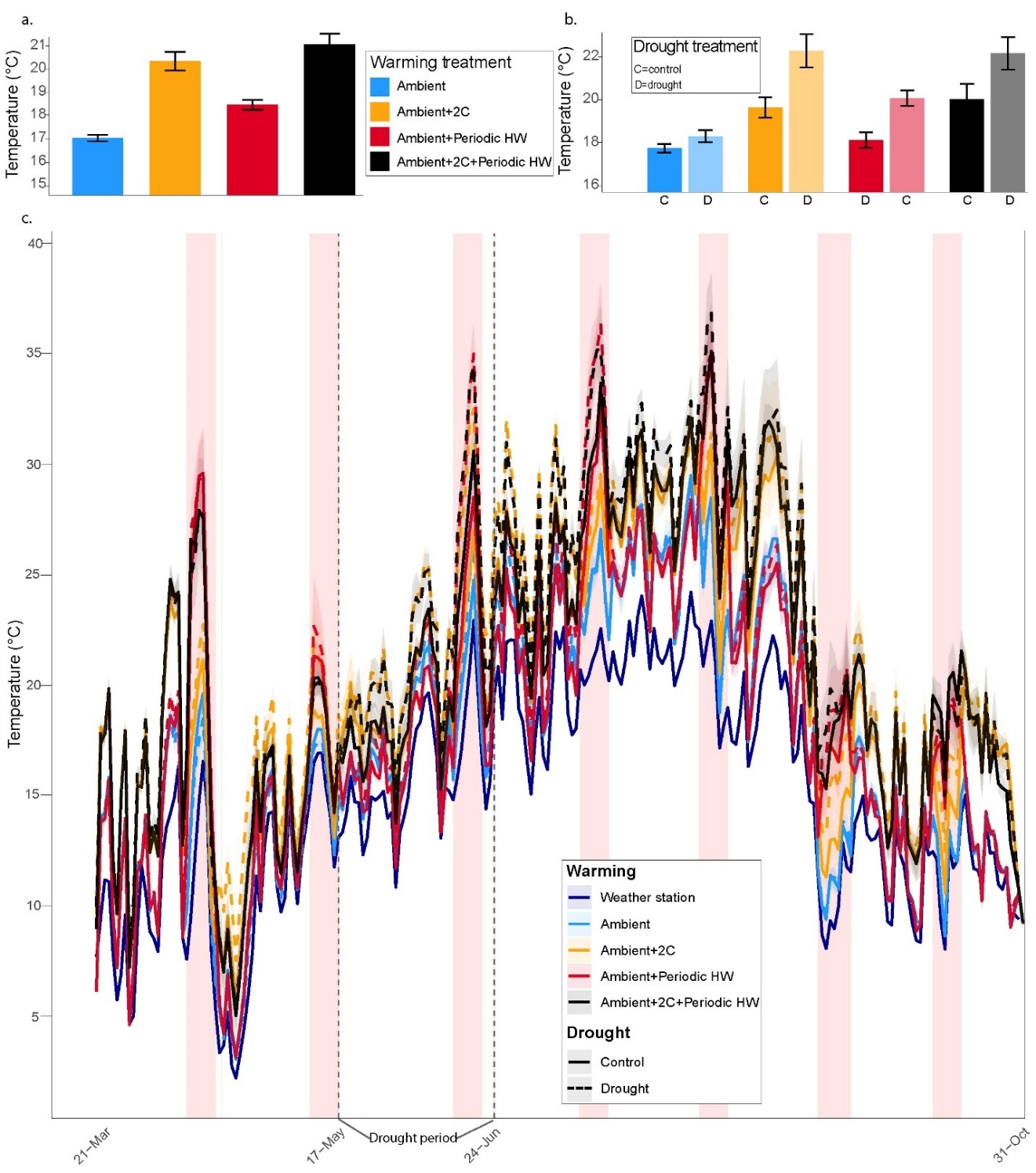

**Figure S5.** Warming treatment temperature measurements taken 15 cm above the soil surface (sensors were placed at the center of each mesocosm directly beneath the infrared heater). (a) Mean temperature (mean ± SE) across treatments for the entire growing season, measured 15 cm above the soil surface. (b) Mean temperature (mean ± SE) across each warming × drought treatment during the 2024 drought period (17 May–24 June). (c) Seasonal temperature dynamics, with dark blue indicating air temperature recorded at the nearby weather station and light blue showing temperature 15 cm above the soil surface, which remained consistently higher due to the sensor’s position above a heat-absorbing surface.

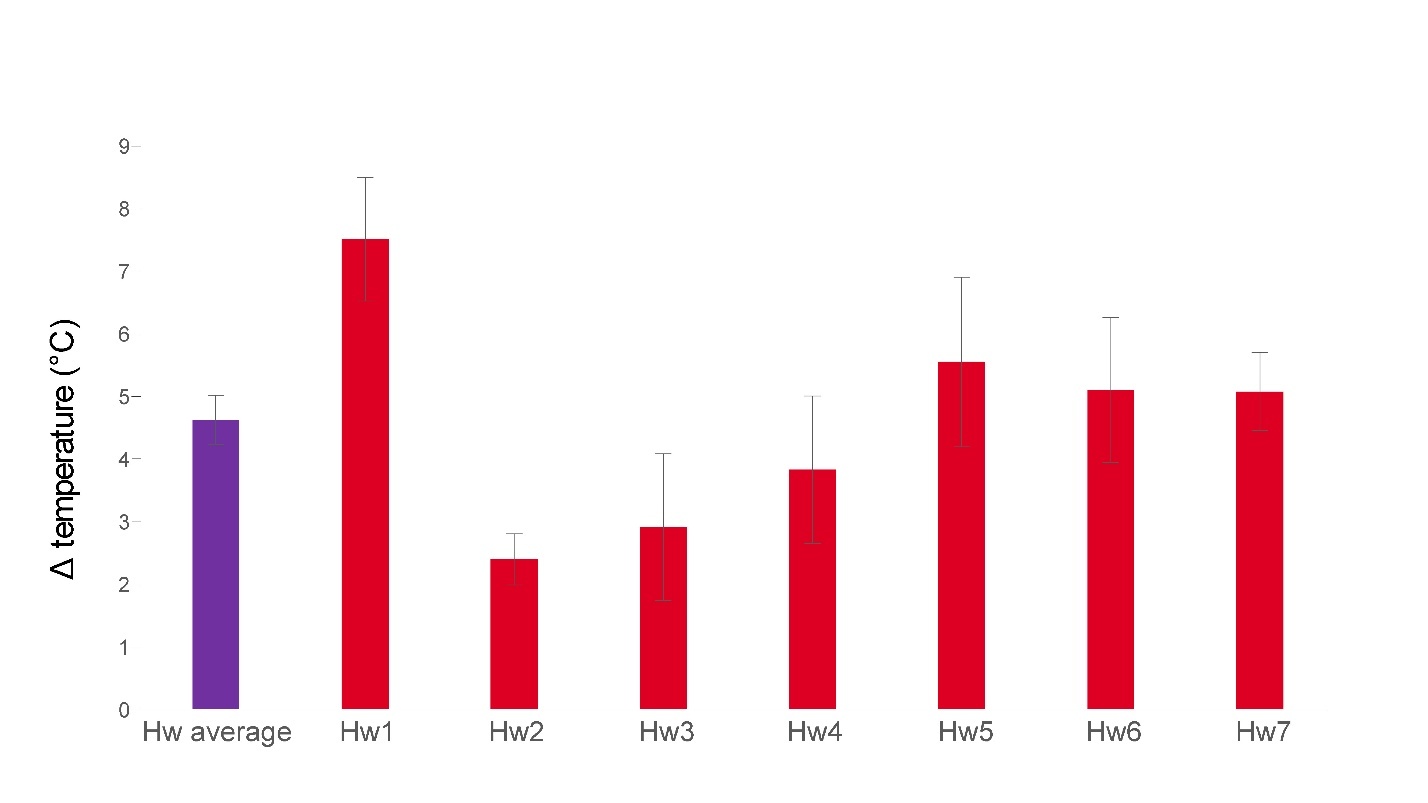
**Figure S6**. Δ temperature between the extreme heat-wave treatment and ambient conditions. The average temperature difference (7-day mean ± SE) measured 15 cm above the soil surface is shown in violet. The Hw (heat-wave) average represents the mean Δ across all seven heat waves relative to ambient temperature (2024 data). In red, Hw1–7 indicate the Δ of each individual heat wave compared to ambient temperature (2024 data).

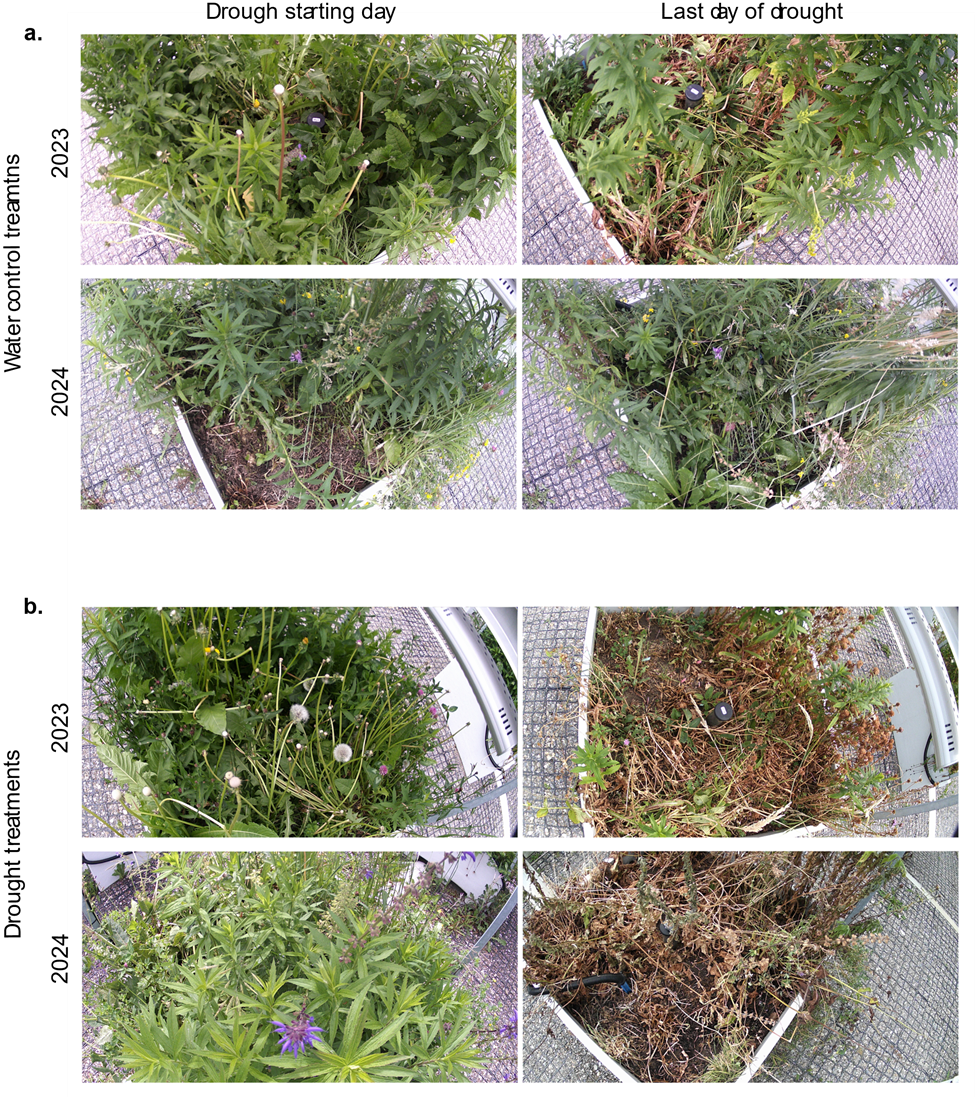

**Figure S7**. Images of four mesocosms at the start and end of the drought periods in 2023 and 2024, illustrating visible differences in canopy cover and plant condition. (a) Control treatment, where plant communities were watered throughout. (b) Drought treatment, where watering was withheld.

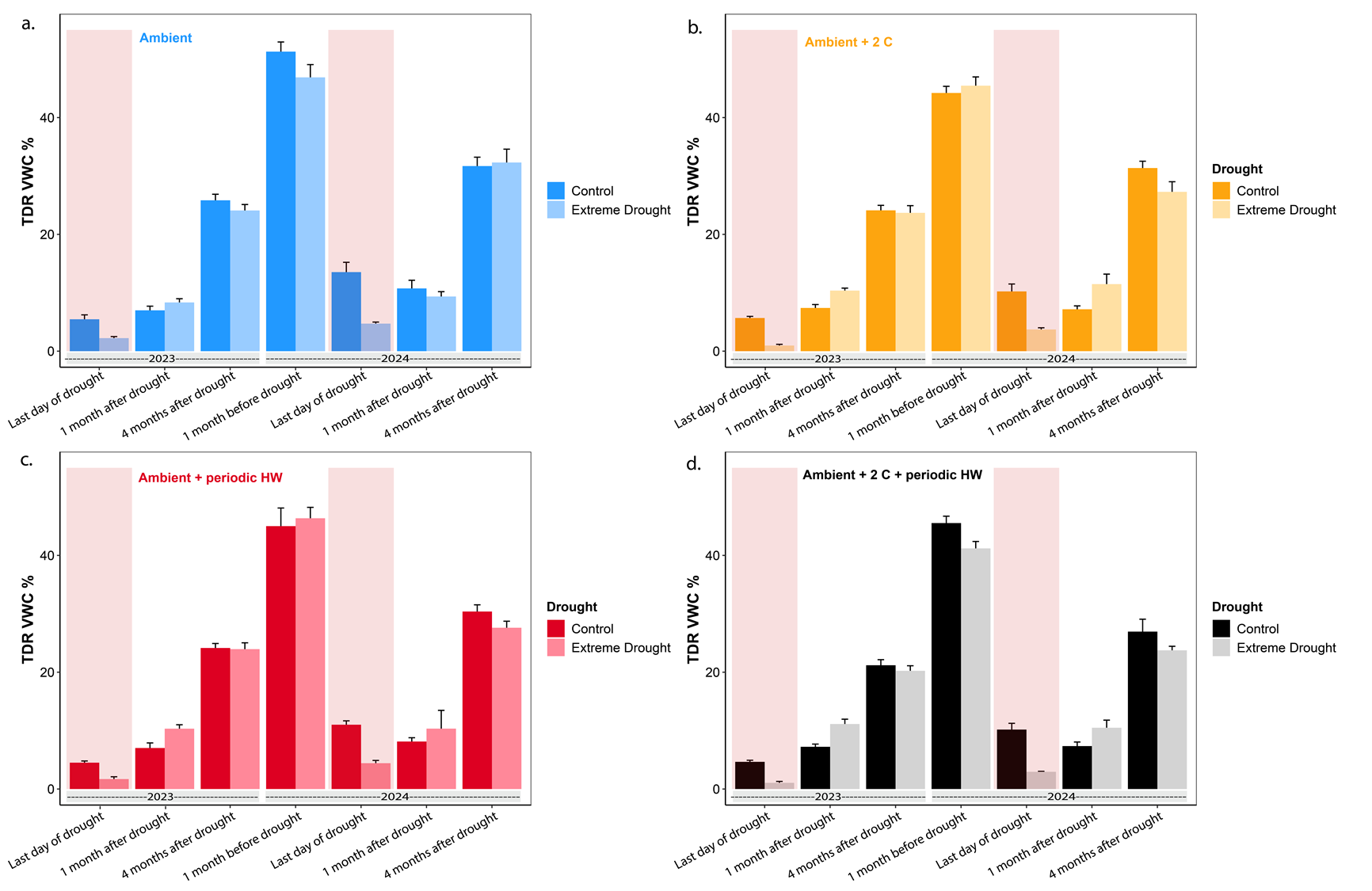

**Figure S8.** Volumetric soil water content (mean ± SE, %) under four warming treatments, measured with TDR at specific time points during the growing season: (a) Ambient (blue), (b) Ambient + 2 °C (orange), (c) Ambient + periodic heat waves (red), and (d) Ambient + 2 °C + periodic heat waves (black). Dark bars indicate control (well-watered) conditions, while light bars represent drought treatments. Shaded red areas denote drought periods.

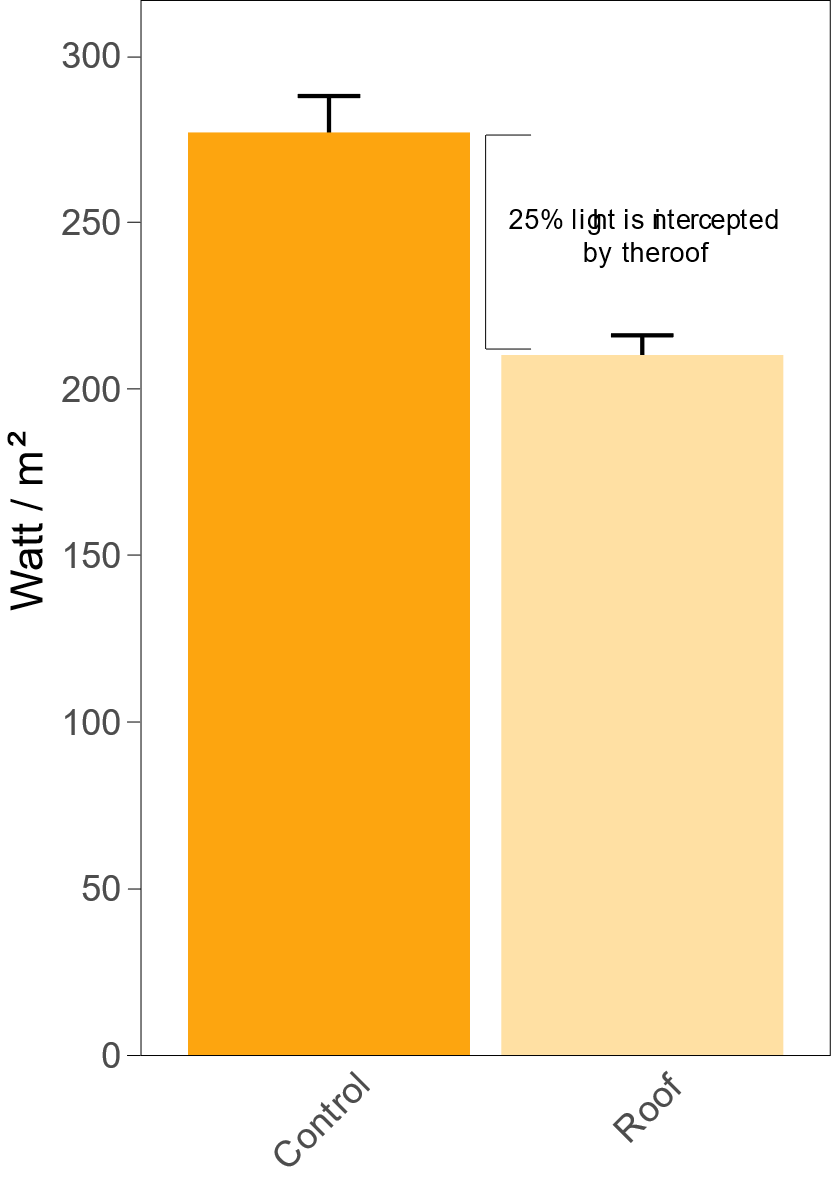

**Figure S9.** Proportion of light (mean ± SE) intercepted by the roof structure placed over mesocosms during the experimental drought period. Light intensity (lux) was measured with sensors positioned either beneath the roof or under open (no-roof) conditions. Measurements were recorded using a HOBO MX2202 data logger (Onset, Bourne, USA) and converted to photosynthetically active radiation (W m⁻²) using a regression model relating lux to W m⁻², established from concurrent weather station readings (R² = 0.97). Note that roofs were placed over all mesocosms during the drought period, resulting in identical light reductions across treatments (see Methods for details).

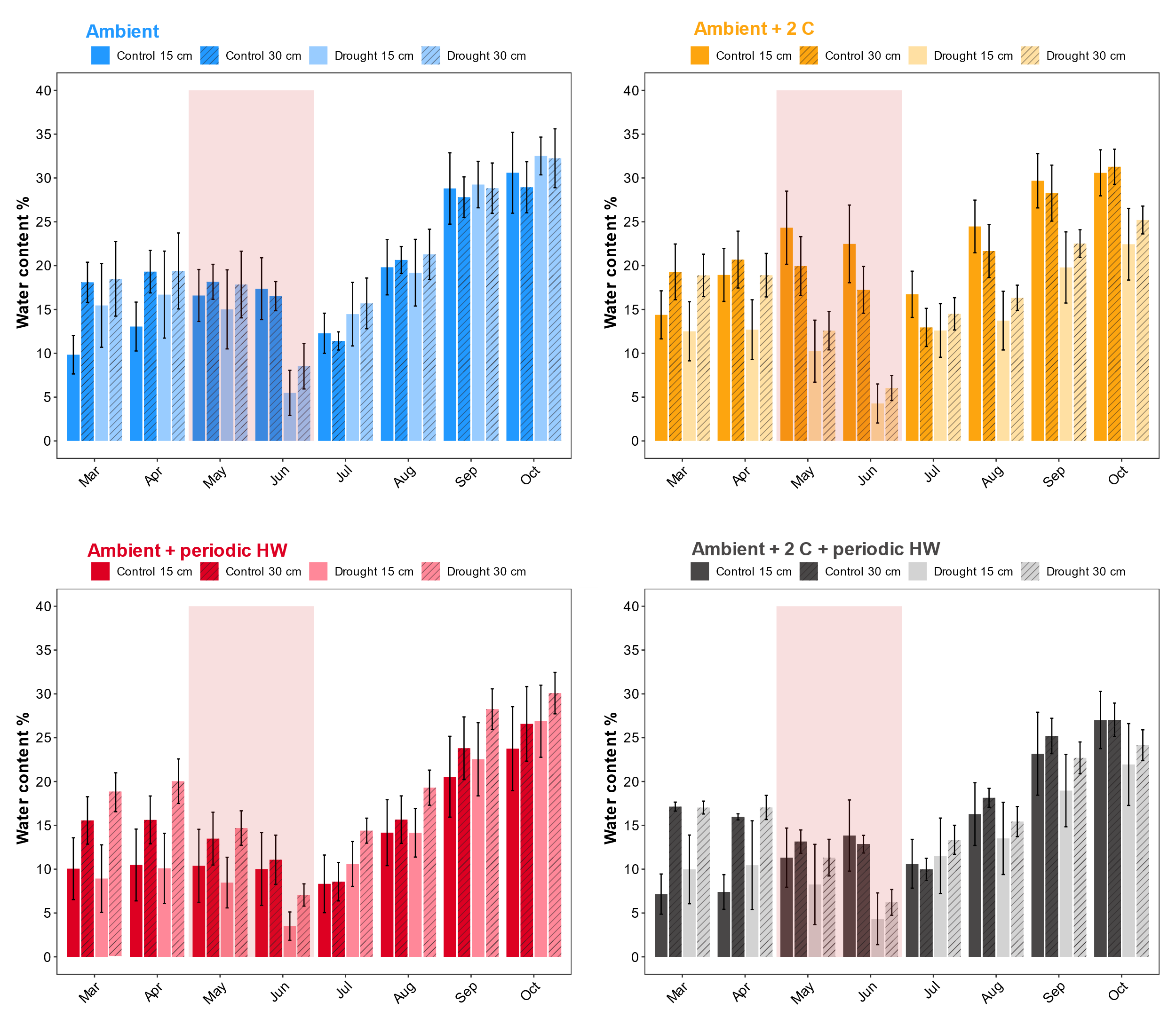

**Figure S10.** Monthly soil water content at two depths during the 2024 experimental year. Values represent mean volumetric water content across four warming regimes (different colours) and two drought treatments (Control: darker shades; Extreme drought: lighter shades). Bars indicate two soil depth intervals: solid bars for 0–15 cm and diagonally striped bars for 15–30 cm. The light red shading over May–June marks the imposed drought period (17 May–24 June 2024). Drought treatment bars are also shown before this period to illustrate legacy effects from the 2023 drought. Soil moisture sensors experienced a technical failure in 2023; therefore, data for that year are unavailable.

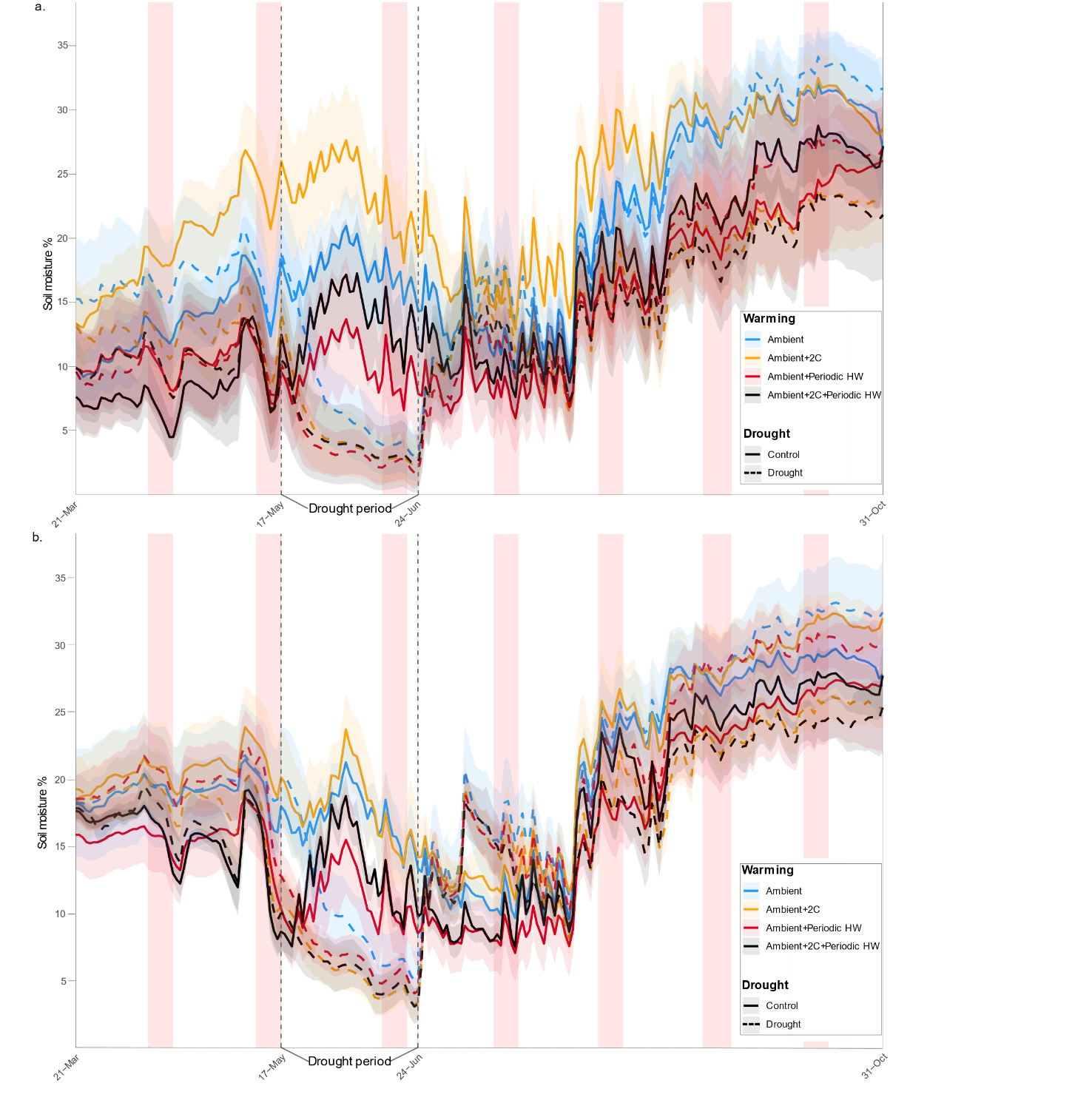

**Figure S11.** Temporal dynamics of soil moisture at two depths during the 2024 growing season. Daily soil moisture (mean ± SE) is shown for the 0–15 cm (panel a) and 15–30 cm (panel b) soil layers. Red squares indicate the extreme heat event (+10 °C above ambient), while vertical dotted lines denote the extreme drought period. Colours represent the four warming regimes, and line styles indicate drought treatments (solid = control; dotted = drought). Soil moisture sensors experienced a technical failure in 2023; therefore, data for that year are unavailable.

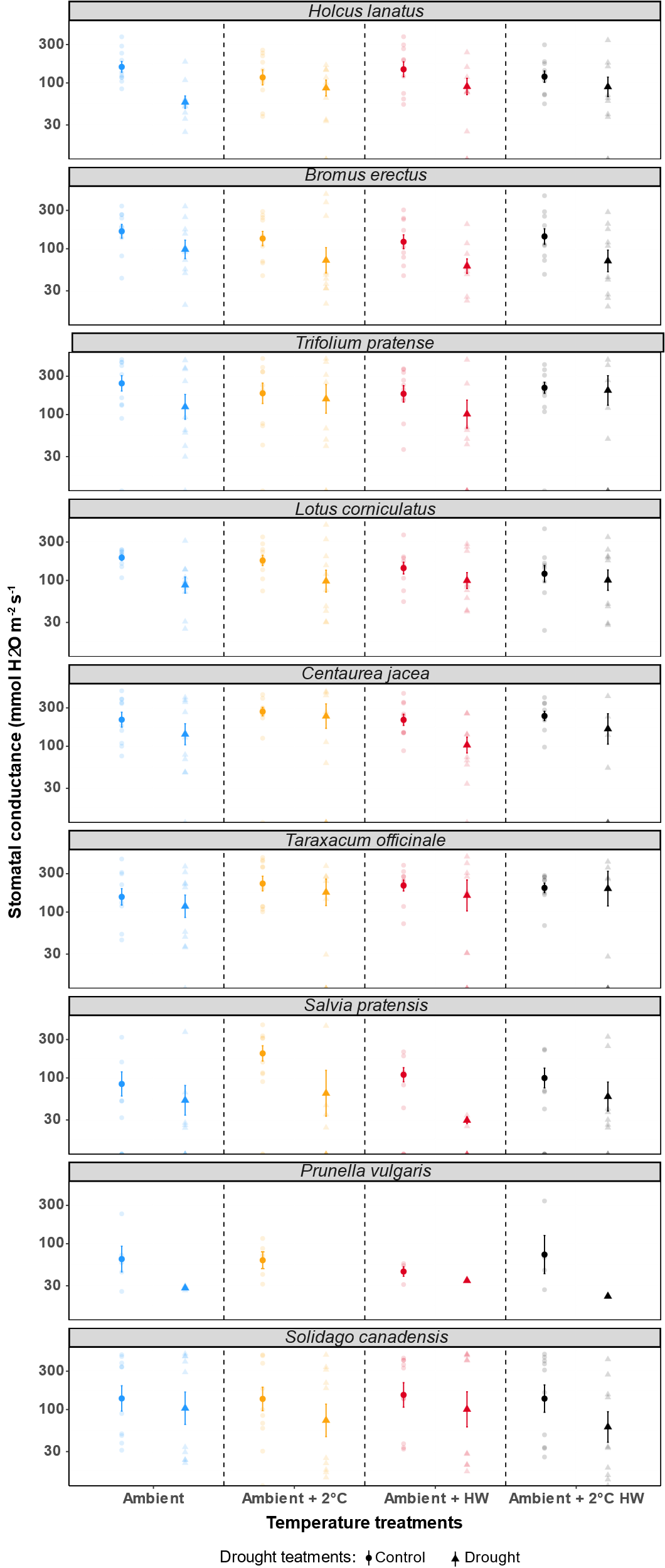

**Figure S12.** Stomatal conductance (mean ± SE) measured on the final day of the drought treatment (data pooled across 2023 and 2024, except for *Prunella vulgaris*). Circles represent control plants and triangles represent drought-stressed plants. Faded points show raw data.

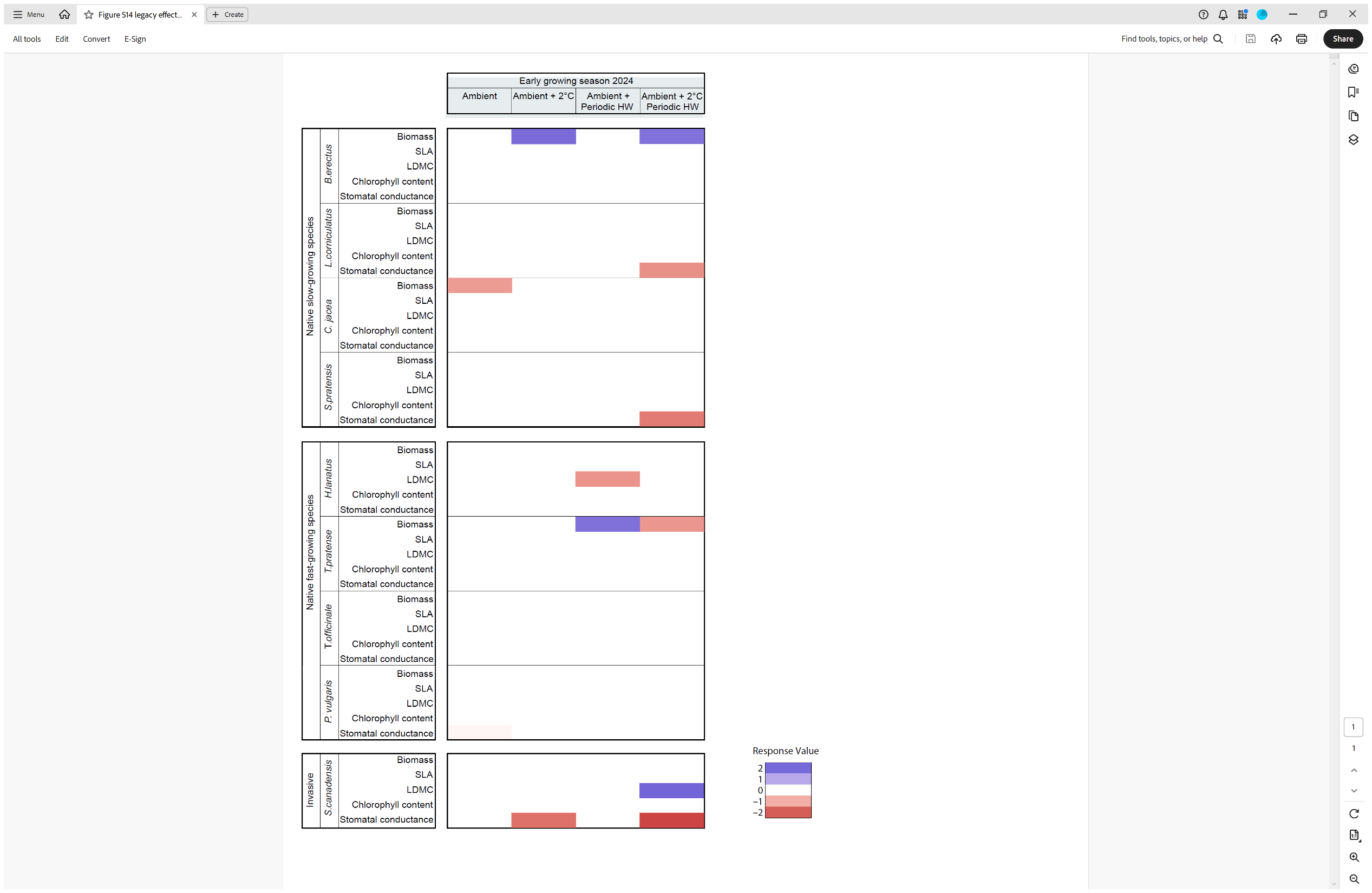

**Figure S13.** Effect size of different warming regimes across all plants before the drought event of 2024 (2023 drought legacy effect of on plant performance in 2024 spring). Data are derived from the same model run used for the 1-month and 4-months recovery table.

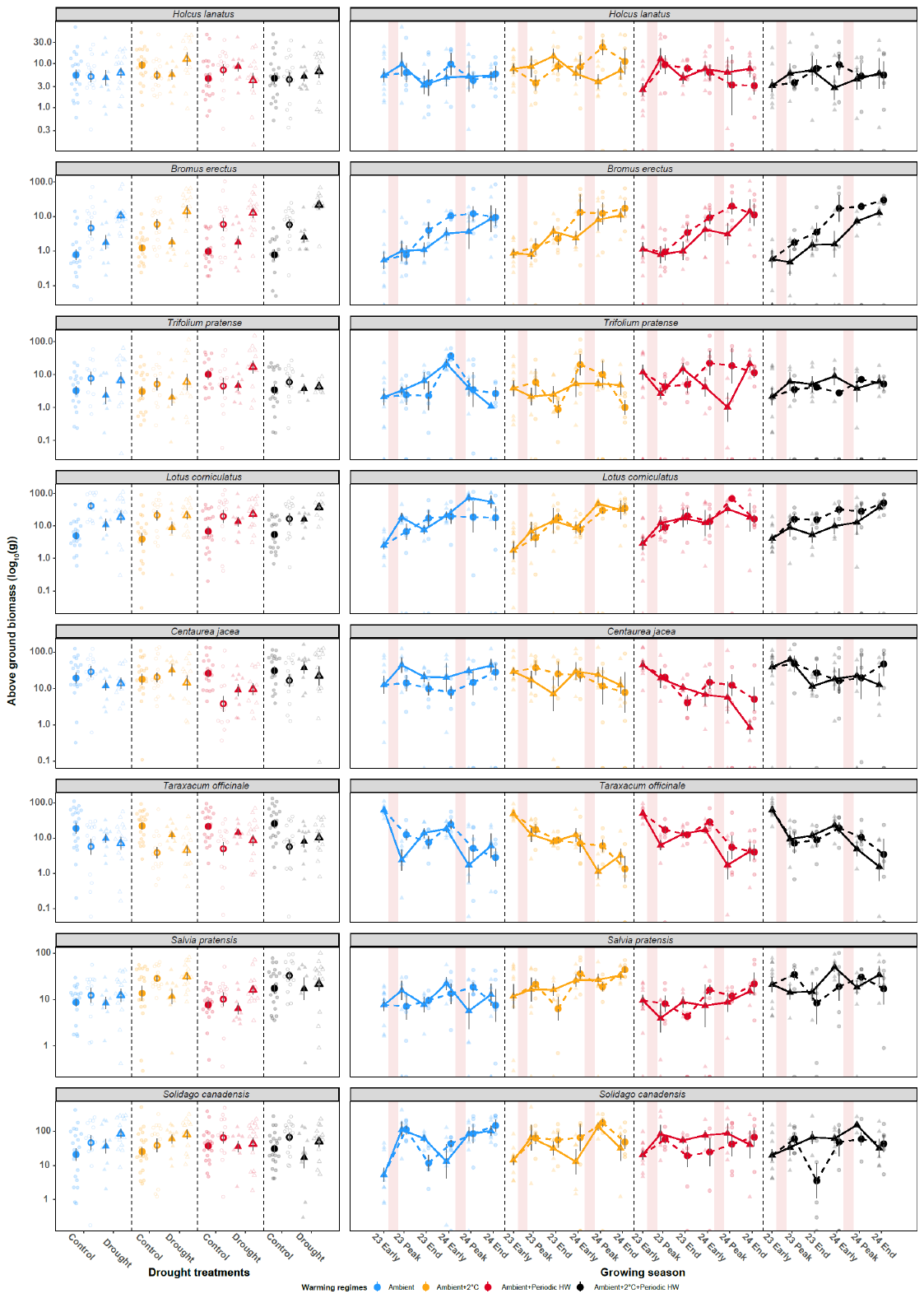

**Figure S14.** Average aboveground biomass per species and year, and biomass dynamics across the two years of data collection. Left panels show yearly biomass production in 2023 and 2024 under different temperature treatments for control and drought conditions; early spring 2023 data are included as control since no drought was applied before the spring harvest. Right panels depict biomass dynamics across the two years. Values are log-transformed for clarity. 23 Early = spring sampling in 2023; 23 Peak = 1 month after drought at the peak of the 2023 growing season; 23 End = 4 months after drought at the end of the 2023 growing season. 24 Early = spring sampling in 2024; 24 Peak = 1 month after drought at the peak of the 2024 growing season; 24 End = 4 months after drought at the end of the 2024 growing season. Light red horizontal bars in the left panels indicate the drought period.

**Table S1.** Initial soil parameters (mean ± SD) for the three soil categories used in the experiment. The artificial substrate refers to the lower layer used to fill the mesocosms. The top layer corresponds to the organic-rich topsoil (0–10 cm, sieved to 2 mm) collected from a nearby natural meadow to inoculate mesocosms with a native soil community. The baseline soil represents the upper 10 cm of soil sampled after plant establishment but before the onset of experimental treatments. SD denotes standard deviation.

| **Soil factors** | **Type of soil** | | |
| --- | --- | --- | --- |
|  | **Artificial substrate– July 2022** | **Top layer - July 2022** | **Baseline soil – November 2022** |
|  | Mean ± SD | Mean ± SD | Mean ± SD |
| **SOM [mg/ g soil]** | 35.52 ± 2.68 | 49.06±8.92 | 80.18±16.85 |
| **Total N [mg/ g soil]** | 2.60±0.14 | 3.41±0.28 | 4.65±0.87 |
| **Soil Organic Carbon [mg/ g soil]** | 20.60±1.56 | 29.95±4.45 | 46.51±9.77 |
| **C: N ratio** | 7.90±0.14 | 8.84±0.67 | 9.99±0.47 |
| **pH** | 7.47±0.05 | 7.21±0.06 | 7.15±0.23 |

**Table S2.** Output of the mixed-effects models testing plant biomass responses to treatments and season across different growth strategies (growth-strategy-specific recovery model). The model included drought, temperature, harvest time point, and plant growth strategy as explanatory variables, with *Solidago canadensis* considered as a growing strategy level due to its invasive status which does not fit usual fast-slow growing strategies. Significant estimates are highlighted in bold. Abbreviations: D = drought treatment; [+2 °C] = ambient +2 °C warming; HW = ambient + heatwave warming (+10 °C); [+2 °C+HW] = ambient +2 °C + heatwave warming; Native fast-growing = native fast-growing species; *S. canadensis* = invasive growth strategy; 1-month recovery = one month after drought (peak growing season); 4-months recovery = 4-months after drought (end of the growing season).

| Predictor | Estimate | SE | Z | p-value |
| --- | --- | --- | --- | --- |
| (Intercept) | 10.088 | 0.287 | 35.171 | <0.001 |
| D | -0.123 | 0.259 | -0.477 | 0.634 |
| [+2°C] | -0.087 | 0.254 | -0.345 | 0.730 |
| HW | -0.401 | 0.271 | -1.478 | 0.139 |
| [+2°C+HW] | -0.032 | 0.258 | -0.123 | 0.902 |
| FG | -0.462 | 0.393 | -1.176 | 0.240 |
| *S. canadensis* | -0.062 | 0.602 | -0.103 | 0.918 |
| 1-Month recovery (1M) | 0.122 | 0.225 | 0.543 | 0.587 |
| 4-Months recovery (4M) | 0.063 | 0.227 | 0.279 | 0.781 |
| D x [+2°C] | 0.296 | 0.364 | 0.813 | 0.416 |
| D x HW | 0.438 | 0.386 | 1.137 | 0.256 |
| D x [+2°C+HW] | 0.291 | 0.369 | 0.789 | 0.430 |
| D x FG | 0.421 | 0.386 | 1.090 | 0.276 |
| D x *S. canadensis* | 0.931 | 0.544 | 1.712 | 0.087 |
| [+2°C] x FG | -0.126 | 0.398 | -0.317 | 0.751 |
| HW x FG | 0.296 | 0.408 | 0.725 | 0.469 |
| [+2°C+HW] x FG | -0.078 | 0.406 | -0.191 | 0.849 |
| [+2°C] x *S. canadensis* | 0.082 | 0.571 | 0.143 | 0.886 |
| **HW x *S. canadensis*** | **1.674** | **0.509** | **3.289** | **0.001** |
| **[+2°C+HW] x *S. canadensis*** | **1.113** | **0.511** | **2.178** | **0.029** |
| D x 1M | -0.118 | 0.325 | -0.362 | 0.717 |
| D x 4M | -0.007 | 0.325 | -0.020 | 0.984 |
| [+2°C] x 1M | 0.016 | 0.318 | 0.052 | 0.959 |
| HW x 1M | 0.030 | 0.337 | 0.089 | 0.929 |
| [+2°C+HW] x 1M | -0.105 | 0.323 | -0.323 | 0.746 |
| [+2°C] x 4M | 0.091 | 0.318 | 0.287 | 0.774 |
| HW x 4M | 0.117 | 0.338 | 0.347 | 0.728 |
| [+2°C+HW] x 4M | -0.029 | 0.321 | -0.090 | 0.928 |
| FG x 1M | -0.567 | 0.357 | -1.589 | 0.112 |
| ***S. canadensis* x 1M** | **1.454** | **0.455** | **3.198** | **0.001** |
| FG x 4M | -0.259 | 0.355 | -0.728 | 0.466 |
| ***S. canadensis* x 4M** | **1.367** | **0.456** | **2.999** | **0.003** |
| D x [+2°C] x FG | -0.400 | 0.558 | -0.716 | 0.474 |
| D x HW x FG | -0.385 | 0.567 | -0.679 | 0.497 |
| D x [+2°C+HW] x FG | -0.486 | 0.576 | -0.844 | 0.399 |
| D x [+2°C] x *S. canadensis* | 0.036 | 0.771 | 0.047 | 0.963 |
| **D x HW x *S. canadensis*** | **-2.123** | **0.750** | **-2.831** | **0.005** |
| D x [+2°C+HW] x *S. canadensis* | -1.291 | 0.743 | -1.739 | 0.082 |
| D x [+2°C] x 1M | -0.059 | 0.456 | -0.129 | 0.897 |
| D x HW x 1M | 0.068 | 0.477 | 0.142 | 0.887 |
| D x [+2°C+HW] x 1M | 0.272 | 0.459 | 0.594 | 0.553 |
| D x [+2°C] x 4M | -0.093 | 0.454 | -0.205 | 0.838 |
| D x HW x 4M | -0.266 | 0.479 | -0.556 | 0.578 |
| D x [+2°C+HW] x 4M | 0.110 | 0.458 | 0.241 | 0.810 |
| D x FG x 1M | -0.001 | 0.502 | -0.002 | 0.998 |
| D x *S. canadensis* x 1M | -0.698 | 0.619 | -1.126 | 0.260 |
| D x FG x 4M | -0.497 | 0.504 | -0.986 | 0.324 |
| **D x *S. canadensis* x 4M** | **-1.245** | **0.631** | **-1.972** | **0.049** |
| [+2°C] x FG x 1M | 0.294 | 0.514 | 0.571 | 0.568 |
| HW x FG x 1M | 0.083 | 0.527 | 0.157 | 0.875 |
| [+2°C+HW] x FG x 1M | 0.421 | 0.517 | 0.814 | 0.415 |
| [+2°C] x *S. canadensis* x 1M | 0.004 | 0.642 | 0.007 | 0.994 |
| **HW x *S. canadensis* x 1M** | **-1.418** | **0.592** | **-2.396** | **0.017** |
| [+2°C+HW] x *S. canadensis* x 1M | -1.165 | 0.593 | -1.964 | 0.050 |
| [+2°C] x FG x 4M | 0.089 | 0.505 | 0.176 | 0.861 |
| HW x FG x 4M | 0.173 | 0.515 | 0.337 | 0.736 |
| [+2°C+HW] x FG x 4M | 0.058 | 0.513 | 0.113 | 0.910 |
| [+2°C] x *S. canadensis* x 4M | -0.833 | 0.660 | -1.262 | 0.207 |
| **HW x *S. canadensis* x 4M** | **-1.861** | **0.603** | **-3.086** | **0.002** |
| **[+2°C+HW] x *S. canadensis* x 4M** | **-1.555** | **0.602** | **-2.583** | **0.010** |
| D x [+2°C] x FG x 1M | 0.346 | 0.719 | 0.482 | 0.630 |
| D x HW x FG x 1M | 0.073 | 0.730 | 0.099 | 0.921 |
| D x [+2°C+HW] x FG x 1M | -0.202 | 0.733 | -0.276 | 0.783 |
| D x [+2°C] x *S. canadensis* x 1M | -0.189 | 0.875 | -0.216 | 0.829 |
| D x HW x *S. canadensis* x 1M | 1.163 | 0.867 | 1.341 | 0.180 |
| D x [+2°C+HW] x *S. canadensis* x 1M | 0.505 | 0.857 | 0.589 | 0.556 |
| D x [+2°C] x FG x 4M | 0.183 | 0.721 | 0.254 | 0.799 |
| D x HW x FG x 4M | 0.253 | 0.723 | 0.350 | 0.726 |
| D x [+2°C+HW] x FG x 4M | 0.359 | 0.739 | 0.486 | 0.627 |
| D x [+2°C] x *S. canadensis* x 4M | 0.568 | 0.902 | 0.630 | 0.529 |
| **D x HW x *S. canadensis* x 4M** | **2.295** | **0.888** | **2.584** | **0.010** |
| D x [+2°C+HW] x *S. canadensis* x 4M | 0.630 | 0.892 | 0.707 | 0.480 |

**Table S3:** Output of the mixed-effects models testing plant SLA to treatments and season across different growth strategies (growth-strategy-specific recovery model). The model included drought, temperature, harvest time point, and plant growth strategy as explanatory variables, with *Solidago canadensis* considered as a growing strategy level due to its invasive status which does not fit usual fast-slow growing strategies. Significant estimates are highlighted in bold. Abbreviations: D = drought treatment; [+2 °C] = ambient +2 °C warming; HW = ambient + heatwave warming (+10 °C); [+2 °C+HW] = ambient +2 °C + heatwave warming; Native fast-growing = native fast-growing species; *S. canadensis* = invasive growth strategy; 1-month recovery = one month after drought (peak growing season); 4-months recovery = 4-months after drought (end of the growing season).

| Predictor | Estimate | SE | Z | p-value |
| --- | --- | --- | --- | --- |
| (Intercept) | 25.792 | 2.576 | 10.012 | <0.001 |
| D | -0.775 | 2.119 | -0.366 | 0.715 |
| [+2°C] | -1.411 | 2.119 | -0.666 | 0.506 |
| HW | -0.869 | 2.119 | -0.410 | 0.682 |
| [+2°C+HW] | -0.449 | 2.145 | -0.209 | 0.834 |
| **FG** | **8.070** | **3.567** | **2.262** | **0.024** |
| *S. canadensis* | -0.728 | 5.641 | -0.129 | 0.897 |
| **1M** | **-5.670** | **1.829** | **-3.100** | **0.002** |
| 4M | -1.402 | 1.803 | -0.777 | 0.437 |
| D x [+2°C] | 2.271 | 3.015 | 0.753 | 0.451 |
| D x HW | 2.577 | 2.997 | 0.860 | 0.390 |
| D x [+2°C+HW] | -0.245 | 3.015 | -0.081 | 0.935 |
| D x FG | -1.058 | 2.933 | -0.361 | 0.718 |
| D x *S. canadensis* | 1.937 | 4.607 | 0.420 | 0.674 |
| [+2°C] x FG | 2.414 | 2.933 | 0.823 | 0.411 |
| HW x FG | 0.092 | 2.914 | 0.032 | 0.975 |
| [+2°C+HW] x FG | -2.987 | 2.952 | -1.012 | 0.312 |
| [+2°C] x *S. canadensis* | 4.162 | 4.607 | 0.903 | 0.366 |
| HW x *S. canadensis* | 6.439 | 4.607 | 1.398 | 0.162 |
| [+2°C+HW] x *S. canadensis* | 0.338 | 4.619 | 0.073 | 0.942 |
| **D x 1M** | **7.549** | **2.552** | **2.958** | **0.003** |
| D x 4M | 1.166 | 2.529 | 0.461 | 0.645 |
| [+2°C] x 1M | 2.352 | 2.547 | 0.923 | 0.356 |
| HW x 1M | 1.163 | 2.558 | 0.454 | 0.649 |
| [+2°C+HW] x 1M | 0.466 | 2.569 | 0.182 | 0.856 |
| [+2°C] x 4M | 0.671 | 2.529 | 0.265 | 0.791 |
| HW x 4M | 0.720 | 2.534 | 0.284 | 0.776 |
| [+2°C+HW] x 4M | -1.288 | 2.551 | -0.505 | 0.614 |
| FG x 1M | 0.347 | 2.578 | 0.135 | 0.893 |
| *S. canadensis* x 1M | 0.089 | 4.005 | 0.022 | 0.982 |
| FG x 4M | -1.304 | 2.540 | -0.513 | 0.608 |
| *S. canadensis* x 4M | -4.122 | 3.993 | -1.032 | 0.302 |
| D x [+2°C] x FG | -3.688 | 4.190 | -0.880 | 0.379 |
| D x HW x FG | -3.230 | 4.199 | -0.769 | 0.442 |
| D x [+2°C+HW] x FG | 2.964 | 4.207 | 0.704 | 0.481 |
| D x [+2°C] x *S. canadensis* | -5.633 | 6.524 | -0.863 | 0.388 |
| D x HW x *S. canadensis* | -8.374 | 6.515 | -1.285 | 0.199 |
| D x [+2°C+HW] x *S. canadensis* | -1.088 | 6.524 | -0.167 | 0.867 |
| D x [+2°C] x 1M | -2.814 | 3.617 | -0.778 | 0.437 |
| D x HW x 1M | -6.426 | 3.605 | -1.783 | 0.075 |
| D x [+2°C+HW] x 1M | -1.134 | 3.605 | -0.314 | 0.753 |
| D x [+2°C] x 4M | -1.543 | 3.588 | -0.430 | 0.667 |
| D x HW x 4M | -3.008 | 3.580 | -0.840 | 0.401 |
| D x [+2°C+HW] x 4M | -0.348 | 3.588 | -0.097 | 0.923 |
| D x FG x 1M | -2.312 | 3.647 | -0.634 | 0.526 |
| D x *S. canadensis* x 1M | -9.512 | 5.697 | -1.670 | 0.095 |
| D x FG x 4M | 1.493 | 3.604 | 0.414 | 0.679 |
| D x *S. canadensis* x 4M | 2.397 | 5.645 | 0.425 | 0.671 |
| [+2°C] x FG x 1M | -5.859 | 3.648 | -1.606 | 0.108 |
| HW x FG x 1M | -3.143 | 3.645 | -0.862 | 0.389 |
| [+2°C+HW] x FG x 1M | -0.650 | 3.668 | -0.177 | 0.859 |
| [+2°C] x *S. canadensis* x 1M | -4.192 | 5.653 | -0.742 | 0.458 |
| HW x *S. canadensis* x 1M | -9.493 | 5.752 | -1.650 | 0.099 |
| [+2°C+HW] x *S. canadensis* x 1M | 2.750 | 5.663 | 0.486 | 0.627 |
| [+2°C] x FG x 4M | -3.371 | 3.596 | -0.937 | 0.349 |
| HW x FG x 4M | -1.309 | 3.592 | -0.365 | 0.715 |
| [+2°C+HW] x FG x 4M | 3.308 | 3.615 | 0.915 | 0.360 |
| [+2°C] x *S. canadensis* x 4M | -1.049 | 5.645 | -0.186 | 0.853 |
| HW x *S. canadensis* x 4M | -7.023 | 5.647 | -1.244 | 0.214 |
| [+2°C+HW] x *S. canadensis* x 4M | 1.857 | 5.655 | 0.328 | 0.743 |
| D x [+2°C] x FG x 1M | 7.922 | 5.204 | 1.523 | 0.128 |
| D x HW x FG x 1M | 7.062 | 5.212 | 1.355 | 0.175 |
| D x [+2°C+HW] x FG x 1M | 0.529 | 5.212 | 0.101 | 0.919 |
| D x [+2°C] x *S. canadensis* x 1M | 8.902 | 8.031 | 1.108 | 0.268 |
| **D x HW x *S. canadensis* x 1M** | **16.518** | **8.092** | **2.041** | **0.041** |
| D x [+2°C+HW] x *S. canadensis* x 1M | -0.935 | 8.055 | -0.116 | 0.908 |
| D x [+2°C] x FG x 4M | 6.358 | 5.135 | 1.238 | 0.216 |
| D x HW x FG x 4M | 5.555 | 5.157 | 1.077 | 0.281 |
| D x [+2°C+HW] x FG x 4M | -2.293 | 5.164 | -0.444 | 0.657 |
| D x [+2°C] x *S. canadensis* x 4M | -1.762 | 7.988 | -0.221 | 0.825 |
| D x HW x *S. canadensis* x 4M | 7.855 | 7.985 | 0.984 | 0.325 |
| D x [+2°C+HW] x *S. canadensis* x 4M | -0.077 | 7.988 | -0.010 | 0.992 |

**Table S4:** Output of the mixed-effects models testing plant LDMC responses to treatments and season across different growth strategies (growth-strategy-specific recovery model). The model included drought, temperature, harvest time point, and plant growth strategy as explanatory variables, with *Solidago canadensis* considered as a growing strategy level due to its invasive status which does not fit usual fast-slow growing strategies. Significant estimates are highlighted in bold. Abbreviations: D = drought treatment; [+2 °C] = ambient +2 °C warming; HW = ambient + heatwave warming (+10 °C); [+2 °C+HW] = ambient +2 °C + heatwave warming; Native fast-growing = native fast-growing species; *S. canadensis* = invasive growth strategy; 1-month recovery = one month after drought (peak growing season); 4-months recovery = 4-months after drought (end of the growing season).

| Predictor | Estimate | SE | Z | p-value |
| --- | --- | --- | --- | --- |
| (Intercept) | 209.884 | 18.393 | 11.411 | <0.001 |
| D | 5.182 | 12.310 | 0.421 | 0.674 |
| [+2°C] | 18.649 | 12.310 | 1.515 | 0.130 |
| HW | 14.938 | 12.310 | 1.214 | 0.225 |
| [+2°C+HW] | 7.678 | 12.466 | 0.616 | 0.538 |
| FG | -23.473 | 25.744 | -0.912 | 0.362 |
| *S. canadensis* | -46.115 | 40.705 | -1.133 | 0.257 |
| 1M | **25.701** | **10.732** | **2.395** | **0.017** |
| 4M | -16.116 | 10.581 | -1.523 | 0.128 |
| D x [+2°C] | -17.845 | 17.408 | -1.025 | 0.305 |
| D x HW | -20.090 | 17.408 | -1.154 | 0.248 |
| D x [+2°C+HW] | -0.768 | 17.519 | -0.044 | 0.965 |
| D x FG | -4.197 | 17.201 | -0.244 | 0.807 |
| D x *S. canadensis* | -2.217 | 27.018 | -0.082 | 0.935 |
| [+2°C] x FG | -12.221 | 17.201 | -0.710 | 0.477 |
| HW x FG | -9.721 | 17.088 | -0.569 | 0.569 |
| [+2°C+HW] x FG | 1.016 | 17.314 | 0.059 | 0.953 |
| [+2°C] x *S. canadensis* | -17.606 | 27.018 | -0.652 | 0.515 |
| HW x *S. canadensis* | -19.630 | 27.018 | -0.727 | 0.468 |
| [+2°C+HW] x *S. canadensis* | -8.810 | 27.090 | -0.325 | 0.745 |
| D x 1M | **-38.181** | **14.967** | **-2.551** | **0.011** |
| D x 4M | -2.138 | 14.830 | -0.144 | 0.885 |
| [+2°C] x 1M | -15.783 | 14.936 | -1.057 | 0.291 |
| HW x 1M | -13.316 | 15.001 | -0.888 | 0.375 |
| [+2°C+HW] x 1M | -11.026 | 15.065 | -0.732 | 0.464 |
| [+2°C] x 4M | -12.014 | 14.830 | -0.810 | 0.418 |
| HW x 4M | -14.703 | 14.862 | -0.989 | 0.323 |
| [+2°C+HW] x 4M | -2.995 | 14.961 | -0.200 | 0.841 |
| FG x 1M | -21.133 | 15.118 | -1.398 | 0.162 |
| ***S. canadensis* x 1M** | **100.202** | **23.486** | **4.267** | **<0.001** |
| FG x 4M | 13.583 | 14.896 | 0.912 | 0.362 |
| ***S. canadensis* x 4M** | **150.124** | **23.419** | **6.410** | **<0.001** |
| D x [+2°C] x FG | 27.643 | 24.493 | 1.129 | 0.259 |
| D x HW x FG | 15.635 | 24.625 | 0.635 | 0.525 |
| D x [+2°C+HW] x FG | -1.080 | 24.673 | -0.044 | 0.965 |
| D x [+2°C] x *S. canadensis* | 21.967 | 38.210 | 0.575 | 0.565 |
| D x HW x *S. canadensis* | 23.218 | 38.210 | 0.608 | 0.543 |
| D x [+2°C+HW] x *S. canadensis* | 56.234 | 38.260 | 1.470 | 0.142 |
| D x [+2°C] x 1M | -13.658 | 21.119 | -0.647 | 0.518 |
| D x HW x 1M | 8.473 | 21.139 | 0.401 | 0.689 |
| D x [+2°C+HW] x 1M | -7.578 | 21.140 | -0.358 | 0.720 |
| D x [+2°C] x 4M | 19.218 | 20.951 | 0.917 | 0.359 |
| D x HW x 4M | 17.976 | 20.996 | 0.856 | 0.392 |
| D x [+2°C+HW] x 4M | 8.989 | 21.043 | 0.427 | 0.669 |
| D x FG x 1M | 13.571 | 21.385 | 0.635 | 0.526 |
| D x *S. canadensis* x 1M | 23.850 | 33.412 | 0.714 | 0.475 |
| D x FG x 4M | -5.251 | 21.111 | -0.249 | 0.804 |
| D x *S. canadensis* x 4M | -57.199 | 33.105 | -1.728 | 0.084 |
| [+2°C] x FG x 1M | 22.197 | 21.363 | 1.039 | 0.299 |
| HW x FG x 1M | 23.208 | 21.377 | 1.086 | 0.278 |
| [+2°C+HW] x FG x 1M | 16.603 | 21.513 | 0.772 | 0.440 |
| [+2°C] x *S. canadensis* x 1M | 2.398 | 33.152 | 0.072 | 0.942 |
| HW x *S. canadensis* x 1M | 24.478 | 33.731 | 0.726 | 0.468 |
| [+2°C+HW] x *S. canadensis* x 1M | 34.120 | 33.211 | 1.027 | 0.304 |
| [+2°C] x FG x 4M | 5.470 | 21.089 | 0.259 | 0.795 |
| HW x FG x 4M | 6.324 | 21.065 | 0.300 | 0.764 |
| [+2°C+HW] x FG x 4M | -2.744 | 21.227 | -0.129 | 0.897 |
| [+2°C] x *S. canadensis* x 4M | -13.763 | 33.105 | -0.416 | 0.678 |
| HW x *S. canadensis* x 4M | 16.314 | 33.119 | 0.493 | 0.622 |
| [+2°C+HW] x *S. canadensis* x 4M | -6.046 | 33.163 | -0.182 | 0.855 |
| D x [+2°C] x FG x 1M | -13.396 | 30.432 | -0.440 | 0.660 |
| D x HW x FG x 1M | -15.329 | 30.568 | -0.501 | 0.616 |
| D x [+2°C+HW] x FG x 1M | -9.604 | 30.566 | -0.314 | 0.753 |
| D x [+2°C] x *S. canadensis* x 1M | 13.941 | 47.057 | 0.296 | 0.767 |
| D x HW x *S. canadensis* x 1M | -31.898 | 47.456 | -0.672 | 0.501 |
| D x [+2°C+HW] x *S. canadensis* x 1M | -63.539 | 47.240 | -1.345 | 0.179 |
| D x [+2°C] x FG x 4M | -23.998 | 30.036 | -0.799 | 0.424 |
| D x HW x FG x 4M | -11.931 | 30.225 | -0.395 | 0.693 |
| D x [+2°C+HW] x FG x 4M | -3.677 | 30.281 | -0.121 | 0.903 |
| D x [+2°C] x *S. canadensis* x 4M | 47.596 | 46.807 | 1.017 | 0.309 |
| D x HW x *S. canadensis* x 4M | 14.512 | 46.827 | 0.310 | 0.757 |
| D x [+2°C+HW] x *S. canadensis* x 4M | -21.019 | 46.848 | -0.449 | 0.654 |

**Table S5:** Output of the mixed-effects models testing plant chlorophyll content responses to treatments and season across different growth strategies (growth-strategy-specific recovery model). The model included drought, temperature, harvest time point, and plant growth strategy as explanatory variables, with *Solidago canadensis* considered as a growing strategy level due to its invasive status which does not fit usual fast-slow growing strategies. Significant estimates are highlighted in bold. Abbreviations: D = drought treatment; [+2 °C] = ambient +2 °C warming; HW = ambient + heatwave warming (+10 °C); [+2 °C+HW] = ambient +2 °C + heatwave warming; Native fast-growing = native fast-growing species; *S. canadensis* = invasive growth strategy; 1-month recovery = one month after drought (peak growing season); 4-months recovery = 4-months after drought (end of the growing season).

| Predictor | Estimate | SE | Z | p-value |
| --- | --- | --- | --- | --- |
| (Intercept) | 10.366 | 0.078 | 133.133 | <0.001 |
| D | -0.075 | 0.063 | -1.193 | 0.233 |
| [+2°C] | -0.019 | 0.063 | -0.305 | 0.760 |
| HW | -0.009 | 0.063 | -0.136 | 0.891 |
| [+2°C+HW] | -0.001 | 0.063 | -0.018 | 0.986 |
| FG | -0.034 | 0.104 | -0.323 | 0.747 |
| *S. canadensis* | -0.172 | 0.164 | -1.047 | 0.295 |
| **1M** | **-0.195** | **0.055** | **-3.561** | **<0.001** |
| 4M | 0.049 | 0.055 | 0.898 | 0.369 |
| D x [+2°C] | 0.051 | 0.089 | 0.579 | 0.563 |
| D x HW | 0.020 | 0.089 | 0.222 | 0.824 |
| D x [+2°C+HW] | 0.074 | 0.089 | 0.832 | 0.405 |
| D x FG | 0.079 | 0.089 | 0.891 | 0.373 |
| D x *S. canadensis* | 0.166 | 0.140 | 1.179 | 0.238 |
| [+2°C] x FG | -0.051 | 0.089 | -0.575 | 0.565 |
| HW x FG | 0.005 | 0.089 | 0.057 | 0.955 |
| [+2°C+HW] x FG | -0.061 | 0.089 | -0.686 | 0.493 |
| [+2°C] x *S. canadensis* | 0.111 | 0.140 | 0.793 | 0.428 |
| HW x *S. canadensis* | 0.084 | 0.140 | 0.595 | 0.552 |
| [+2°C+HW] x *S. canadensis* | -0.001 | 0.140 | -0.007 | 0.994 |
| D x 1M | 0.120 | 0.077 | 1.551 | 0.121 |
| D x 4M | 0.087 | 0.077 | 1.128 | 0.259 |
| [+2°C] x 1M | 0.041 | 0.077 | 0.536 | 0.592 |
| HW x 1M | 0.013 | 0.077 | 0.166 | 0.868 |
| [+2°C+HW] x 1M | -0.059 | 0.077 | -0.768 | 0.442 |
| [+2°C] x 4M | 0.073 | 0.077 | 0.952 | 0.341 |
| HW x 4M | 0.059 | 0.077 | 0.763 | 0.446 |
| [+2°C+HW] x 4M | 0.032 | 0.077 | 0.409 | 0.683 |
| FG x 1M | 0.094 | 0.077 | 1.220 | 0.223 |
| *S. canadensis* x 1M | 0.124 | 0.122 | 1.019 | 0.308 |
| FG x 4M | 0.080 | 0.078 | 1.034 | 0.301 |
| *S. canadensis* x 4M | 0.109 | 0.122 | 0.898 | 0.369 |
| D x [+2°C] x FG | -0.013 | 0.126 | -0.103 | 0.918 |
| D x HW x FG | -0.060 | 0.127 | -0.473 | 0.636 |
| D x [+2°C+HW] x FG | 0.036 | 0.128 | 0.278 | 0.781 |
| D x [+2°C] x *S. canadensis* | -0.074 | 0.199 | -0.374 | 0.709 |
| D x HW x *S. canadensis* | -0.022 | 0.199 | -0.113 | 0.910 |
| D x [+2°C+HW] x *S. canadensis* | 0.000 | 0.199 | 0.001 | 0.999 |
| D x [+2°C] x 1M | -0.051 | 0.109 | -0.470 | 0.638 |
| D x HW x 1M | -0.004 | 0.109 | -0.037 | 0.971 |
| D x [+2°C+HW] x 1M | 0.030 | 0.109 | 0.280 | 0.780 |
| D x [+2°C] x 4M | -0.046 | 0.109 | -0.418 | 0.676 |
| D x HW x 4M | -0.046 | 0.109 | -0.418 | 0.676 |
| D x [+2°C+HW] x 4M | -0.067 | 0.109 | -0.615 | 0.539 |
| D x FG x 1M | -0.125 | 0.109 | -1.151 | 0.250 |
| D x *S. canadensis* x 1M | -0.015 | 0.172 | -0.088 | 0.930 |
| D x FG x 4M | -0.092 | 0.109 | -0.839 | 0.401 |
| D x *S. canadensis* x 4M | -0.114 | 0.172 | -0.662 | 0.508 |
| [+2°C] x FG x 1M | 0.053 | 0.109 | 0.489 | 0.625 |
| HW x FG x 1M | 0.023 | 0.110 | 0.214 | 0.831 |
| [+2°C+HW] x FG x 1M | 0.146 | 0.109 | 1.332 | 0.183 |
| [+2°C] x *S. canadensis* x 1M | -0.118 | 0.172 | -0.684 | 0.494 |
| HW x *S. canadensis* x 1M | -0.024 | 0.172 | -0.140 | 0.888 |
| [+2°C+HW] x *S. canadensis* x 1M | 0.131 | 0.172 | 0.761 | 0.447 |
| [+2°C] x FG x 4M | 0.073 | 0.109 | 0.668 | 0.504 |
| HW x FG x 4M | -0.030 | 0.110 | -0.268 | 0.789 |
| [+2°C+HW] x FG x 4M | 0.014 | 0.110 | 0.129 | 0.897 |
| [+2°C] x *S. canadensis* x 4M | -0.138 | 0.172 | -0.803 | 0.422 |
| HW x *S. canadensis* x 4M | -0.149 | 0.172 | -0.863 | 0.388 |
| [+2°C+HW] x *S. canadensis* x 4M | -0.038 | 0.172 | -0.221 | 0.825 |
| D x [+2°C] x FG x 1M | 0.027 | 0.155 | 0.175 | 0.861 |
| D x HW x FG x 1M | 0.083 | 0.156 | 0.533 | 0.594 |
| D x [+2°C+HW] x FG x 1M | -0.059 | 0.157 | -0.375 | 0.707 |
| D x [+2°C] x *S. canadensis* x 1M | 0.111 | 0.243 | 0.456 | 0.648 |
| D x HW x *S. canadensis* x 1M | 0.047 | 0.243 | 0.193 | 0.847 |
| D x [+2°C+HW] x *S. canadensis* x 1M | -0.111 | 0.243 | -0.458 | 0.647 |
| D x [+2°C] x FG x 4M | -0.049 | 0.156 | -0.318 | 0.751 |
| D x HW x FG x 4M | 0.090 | 0.157 | 0.574 | 0.566 |
| D x [+2°C+HW] x FG x 4M | -0.002 | 0.158 | -0.012 | 0.990 |
| D x [+2°C] x *S. canadensis* x 4M | 0.045 | 0.243 | 0.185 | 0.853 |
| D x HW x *S. canadensis* x 4M | 0.071 | 0.244 | 0.290 | 0.772 |
| D x [+2°C+HW] x *S. canadensis* x 4M | 0.009 | 0.243 | 0.036 | 0.971 |

**Table S6:** Output of the mixed-effects model testing plants’ stomatal conductance responses to treatments and season across different growth strategies (growth-strategy-specific recovery model). The model included drought, temperature, harvest time point, and plant growth strategy as explanatory variables, with *Solidago canadensis* considered as a growing strategy level due to its invasive status which does not fit usual fast-slow growing strategies. Significant estimates are highlighted in bold. Abbreviations: D = drought treatment; [+2 °C] = ambient +2 °C warming; HW = ambient + heatwave warming (+10 °C); [+2 °C+HW] = ambient +2 °C + heatwave warming; Native fast-growing = native fast-growing species; *S. canadensis* = invasive growth strategy; 1-month recovery = one month after drought (peak growing season); 4-months recovery = 4-months after drought (end of the growing season).

| Term | Estimate | SE | Z | p-value |
| --- | --- | --- | --- | --- |
| (Intercept) | 5.446 | 0.206 | 26.390 | <0.001 |
| D | -0.265 | 0.181 | -1.464 | 0.143 |
| [+2°C] | -0.218 | 0.181 | -1.205 | 0.228 |
| HW | -0.310 | 0.181 | -1.715 | 0.086 |
| [+2°C+HW] | -0.073 | 0.181 | -0.402 | 0.688 |
| FG | -0.248 | 0.285 | -0.869 | 0.385 |
| *S. canadensis* | -0.696 | 0.450 | -1.545 | 0.122 |
| **1M** | **-0.390** | **0.161** | **-2.423** | **0.015** |
| 4M | -0.061 | 0.158 | -0.388 | 0.698 |
| D x [+2°C] | 0.120 | 0.256 | 0.468 | 0.640 |
| D x HW | 0.223 | 0.256 | 0.873 | 0.383 |
| D x [+2°C+HW] | -0.196 | 0.256 | -0.768 | 0.443 |
| D x FG | 0.127 | 0.256 | 0.498 | 0.619 |
| D x *S. canadensis* | -0.218 | 0.404 | -0.540 | 0.589 |
| [+2°C] x FG | 0.234 | 0.256 | 0.916 | 0.360 |
| HW x FG | 0.314 | 0.259 | 1.209 | 0.227 |
| [+2°C+HW] x FG | -0.053 | 0.257 | -0.208 | 0.836 |
| [+2°C] x *S. canadensis* | 0.326 | 0.404 | 0.806 | 0.420 |
| HW x *S. canadensis* | 0.133 | 0.404 | 0.330 | 0.741 |
| [+2°C+HW] x *S. canadensis* | 0.191 | 0.404 | 0.473 | 0.636 |
| D x 1M | 0.207 | 0.227 | 0.911 | 0.362 |
| D x 4M | 0.240 | 0.222 | 1.081 | 0.280 |
| [+2°C] x 1M | 0.339 | 0.226 | 1.498 | 0.134 |
| HW x 1M | 0.015 | 0.226 | 0.068 | 0.946 |
| [+2°C+HW] x 1M | -0.121 | 0.226 | -0.536 | 0.592 |
| [+2°C] x 4M | 0.131 | 0.222 | 0.591 | 0.555 |
| HW x 4M | 0.302 | 0.223 | 1.353 | 0.176 |
| [+2°C+HW] x 4M | -0.014 | 0.222 | -0.061 | 0.951 |
| FG x 1M | 0.110 | 0.227 | 0.484 | 0.628 |
| *S. canadensis* x 1M | 0.515 | 0.352 | 1.462 | 0.144 |
| FG x 4M | 0.257 | 0.225 | 1.145 | 0.252 |
| *S. canadensis* x 4M | 0.312 | 0.351 | 0.890 | 0.373 |
| D x [+2°C] x FG | -0.257 | 0.364 | -0.705 | 0.481 |
| D x HW x FG | -0.235 | 0.368 | -0.638 | 0.524 |
| D x [+2°C+HW] x FG | 0.267 | 0.369 | 0.725 | 0.468 |
| D x [+2°C] x *S. canadensis* | -0.379 | 0.572 | -0.663 | 0.507 |
| D x HW x *S. canadensis* | -0.244 | 0.572 | -0.427 | 0.669 |
| D x [+2°C+HW] x *S. canadensis* | -0.283 | 0.572 | -0.494 | 0.621 |
| D x [+2°C] x 1M | -0.104 | 0.320 | -0.325 | 0.745 |
| D x HW x 1M | 0.070 | 0.321 | 0.219 | 0.827 |
| **D x [+2°C+HW] x 1M** | **0.710** | **0.321** | **2.210** | **0.027** |
| D x [+2°C] x 4M | -0.055 | 0.314 | -0.176 | 0.860 |
| D x HW x 4M | -0.095 | 0.315 | -0.302 | 0.763 |
| D x [+2°C+HW] x 4M | 0.383 | 0.314 | 1.221 | 0.222 |
| D x FG x 1M | 0.158 | 0.321 | 0.492 | 0.623 |
| D x *S. canadensis* x 1M | 0.433 | 0.498 | 0.869 | 0.385 |
| D x FG x 4M | -0.204 | 0.316 | -0.646 | 0.518 |
| D x *S. canadensis* x 4M | 0.094 | 0.496 | 0.190 | 0.850 |
| [+2°C] x FG x 1M | -0.420 | 0.321 | -1.308 | 0.191 |
| HW x FG x 1M | -0.166 | 0.324 | -0.512 | 0.609 |
| [+2°C+HW] x FG x 1M | 0.238 | 0.323 | 0.738 | 0.460 |
| [+2°C] x *S. canadensis* x 1M | -0.476 | 0.501 | -0.950 | 0.342 |
| HW x *S. canadensis* x 1M | 0.231 | 0.497 | 0.464 | 0.643 |
| [+2°C+HW] x *S. canadensis* x 1M | 0.270 | 0.497 | 0.544 | 0.587 |
| [+2°C] x FG x 4M | -0.213 | 0.315 | -0.675 | 0.500 |
| HW x FG x 4M | -0.342 | 0.319 | -1.069 | 0.285 |
| [+2°C+HW] x FG x 4M | 0.038 | 0.318 | 0.120 | 0.905 |
| [+2°C] x *S. canadensis* x 4M | -0.302 | 0.496 | -0.610 | 0.542 |
| HW x *S. canadensis* x 4M | 0.105 | 0.496 | 0.212 | 0.832 |
| [+2°C+HW] x *S. canadensis* x 4M | -0.080 | 0.496 | -0.161 | 0.872 |
| D x [+2°C] x FG x 1M | 0.140 | 0.458 | 0.306 | 0.759 |
| D x HW x FG x 1M | 0.098 | 0.462 | 0.213 | 0.831 |
| **D x [+2°C+HW] x FG x 1M** | **-0.938** | **0.463** | **-2.024** | **0.043** |
| D x [+2°C] x *S. canadensis* x 1M | 0.288 | 0.706 | 0.408 | 0.683 |
| D x HW x *S. canadensis* x 1M | -0.022 | 0.704 | -0.031 | 0.976 |
| D x [+2°C+HW] x *S. canadensis* x 1M | -0.863 | 0.707 | -1.221 | 0.222 |
| D x [+2°C] x FG x 4M | 0.495 | 0.448 | 1.104 | 0.270 |
| D x HW x FG x 4M | 0.341 | 0.454 | 0.753 | 0.452 |
| D x [+2°C+HW] x FG x 4M | -0.180 | 0.454 | -0.395 | 0.693 |
| D x [+2°C] x *S. canadensis* x 4M | 0.606 | 0.701 | 0.864 | 0.388 |
| D x HW x *S. canadensis* x 4M | 0.113 | 0.701 | 0.161 | 0.872 |
| D x [+2°C+HW] x *S. canadensis* x 4M | -0.014 | 0.701 | -0.020 | 0.984 |

**Table S7a.** Output of the mixed-effects models testing specific plant biomass responses to treatments and season (species-specific recovery model). The model included drought, temperature, harvest time point, and plant species as explanatory variables. Due to model-fitting issues, *Prunella vulgaris* was analysed separately in an independent model (see table S7b). Significant estimates are highlighted in bold. Abbreviations: D: Water treatment drought, [+2°C]: Ambient temperature plus constant 2°C warming, HW: Ambient temperature plus heatwave warming events (+10°C), [+2°C+HW]: Ambient temperature plus constant 2°C and heatwave warming. 1M: collection point at 1-month recovery after the end of drought (peak of the growing season), 4M: Data collection point at 4-months recovery after the end of drought (end of the growing season). *Bromus erectus*: Be, *Trifolium pratense*: Tp, *Lotus corniculatus*: Lc, *Taraxacum officinale*: To, *Centaurea jacea*: Cj, *Salvia pratense*: Sp, *Solidago canadensis*: Sc. The intercept is the fast-growing species *Holcus lanatus*.

| Predictor | Estimate | SE | Z | p-value |
| --- | --- | --- | --- | --- |
| (Intercept) | 8.692 | 0.459 | 18.953 | <0.001 |
| D | 0.667 | 0.608 | 1.098 | 0.272 |
| [+2°C] | 0.007 | 0.608 | 0.012 | 0.991 |
| HW | 0.277 | 0.608 | 0.456 | 0.648 |
| [+2°C+HW] | -0.333 | 0.645 | -0.517 | 0.605 |
| Be | -0.424 | 0.608 | -0.697 | 0.486 |
| Tp | **1.270** | **0.608** | **2.090** | **0.037** |
| Lc | 1.193 | 0.608 | 1.962 | 0.050 |
| Cj | **1.753** | **0.608** | **2.883** | **0.004** |
| To | 1.018 | 0.608 | 1.675 | 0.094 |
| Sp | **1.293** | **0.608** | **2.128** | **0.033** |
| Sc | **1.822** | **0.608** | **2.997** | **0.003** |
| 1M | 1.033 | 0.529 | 1.954 | 0.051 |
| 4M | -0.141 | 0.528 | -0.267 | 0.790 |
| D x [+2°C] | 0.018 | 0.860 | 0.021 | 0.983 |
| D x HW | -0.752 | 0.860 | -0.875 | 0.382 |
| D x [+2°C+HW] | 0.324 | 0.886 | 0.366 | 0.714 |
| D x Be | 0.128 | 0.860 | 0.149 | 0.882 |
| D x Tp | -0.260 | 0.886 | -0.294 | 0.769 |
| D x Lc | -0.527 | 0.860 | -0.614 | 0.540 |
| D x Cj | **-2.153** | **0.886** | **-2.429** | **0.015** |
| D x To | -0.303 | 0.860 | -0.353 | 0.724 |
| D x Sp | -0.996 | 0.860 | -1.159 | 0.246 |
| D x Sc | -0.640 | 0.886 | -0.722 | 0.470 |
| [+2°C] x Be | -0.182 | 0.860 | -0.212 | 0.832 |
| HW x Be | 0.381 | 0.860 | 0.443 | 0.658 |
| [+2°C+HW] x Be | 0.261 | 0.912 | 0.286 | 0.775 |
| [+2°C] x Tp | -0.534 | 0.860 | -0.621 | 0.535 |
| HW x Tp | -1.630 | 0.886 | -1.839 | 0.066 |
| [+2°C+HW] x Tp | -0.335 | 0.912 | -0.367 | 0.714 |
| [+2°C] x Lc | -0.682 | 0.860 | -0.793 | 0.428 |
| HW x Lc | -0.022 | 0.860 | -0.025 | 0.980 |
| [+2°C+HW] x Lc | -0.035 | 0.912 | -0.039 | 0.969 |
| [+2°C] x Cj | 0.174 | 0.860 | 0.202 | 0.840 |
| HW x Cj | -1.415 | 0.860 | -1.646 | 0.100 |
| [+2°C+HW] x Cj | 0.156 | 0.886 | 0.176 | 0.861 |
| [+2°C] x To | -0.020 | 0.860 | -0.023 | 0.981 |
| HW x To | -0.314 | 0.860 | -0.365 | 0.715 |
| [+2°C+HW] x To | 0.618 | 0.886 | 0.697 | 0.486 |
| [+2°C] x Sp | 0.121 | 0.860 | 0.141 | 0.888 |
| HW x Sp | -0.614 | 0.886 | -0.693 | 0.489 |
| [+2°C+HW] x Sp | 1.068 | 0.886 | 1.206 | 0.228 |
| [+2°C] x Sc | -0.357 | 0.860 | -0.415 | 0.678 |
| HW x Sc | 0.456 | 0.860 | 0.530 | 0.596 |
| [+2°C+HW] x Sc | 0.804 | 0.886 | 0.908 | 0.364 |
| D x 1M | -1.265 | 0.745 | -1.699 | 0.089 |
| D x 4M | -0.240 | 0.745 | -0.322 | 0.747 |
| [+2°C] x 1M | -0.519 | 0.745 | -0.697 | 0.486 |
| HW x 1M | -0.194 | 0.745 | -0.260 | 0.795 |
| [+2°C+HW] x 1M | -0.312 | 0.775 | -0.402 | 0.687 |
| [+2°C] x 4M | 1.221 | 0.745 | 1.639 | 0.101 |
| HW x 4M | 0.103 | 0.745 | 0.138 | 0.890 |
| [+2°C+HW] x 4M | 1.393 | 0.775 | 1.797 | 0.072 |
| Be x 1M | -0.711 | 0.745 | -0.954 | 0.340 |
| Tp x 1M | **-2.335** | **0.745** | **-3.134** | **0.002** |
| Lc x 1M | -0.183 | 0.760 | -0.241 | 0.810 |
| Cj x 1M | -0.580 | 0.751 | -0.771 | 0.440 |
| To x 1M | **-1.979** | **0.745** | **-2.657** | **0.008** |
| Sp x 1M | -1.299 | 0.771 | -1.686 | 0.092 |
| Sc x 1M | 0.419 | 0.751 | 0.558 | 0.577 |
| Be x 4M | 1.184 | 0.745 | 1.589 | 0.112 |
| Tp x 4M | -0.573 | 0.785 | -0.730 | 0.465 |
| Lc x 4M | 0.670 | 0.760 | 0.881 | 0.378 |
| Cj x 4M | 0.233 | 0.760 | 0.306 | 0.759 |
| To x 4M | 0.069 | 0.745 | 0.093 | 0.926 |
| Sp x 4M | -0.428 | 0.760 | -0.564 | 0.573 |
| Sc x 4M | 1.075 | 0.745 | 1.444 | 0.149 |
| D x [+2°C] x Be | 1.301 | 1.235 | 1.054 | 0.292 |
| D x HW x Be | 0.228 | 1.216 | 0.188 | 0.851 |
| D x [+2°C+HW] x Be | 0.906 | 1.272 | 0.713 | 0.476 |
| D x [+2°C] x Tp | 0.627 | 1.235 | 0.508 | 0.612 |
| D x HW x Tp | 2.393 | 1.253 | 1.909 | 0.056 |
| D x [+2°C+HW] x Tp | -2.304 | 1.376 | -1.674 | 0.094 |
| D x [+2°C] x Lc | -0.388 | 1.235 | -0.314 | 0.754 |
| D x HW x Lc | 0.224 | 1.216 | 0.184 | 0.854 |
| D x [+2°C+HW] x Lc | 0.288 | 1.301 | 0.221 | 0.825 |
| D x [+2°C] x Cj | 0.970 | 1.235 | 0.786 | 0.432 |
| D x HW x Cj | **2.595** | **1.253** | **2.071** | **0.038** |
| D x [+2°C+HW] x Cj | 1.365 | 1.253 | 1.090 | 0.276 |
| D x [+2°C] x To | -0.797 | 1.216 | -0.655 | 0.512 |
| D x HW x To | 0.911 | 1.216 | 0.749 | 0.454 |
| D x [+2°C+HW] x To | -0.808 | 1.235 | -0.655 | 0.513 |
| D x [+2°C] x Sp | 0.686 | 1.216 | 0.564 | 0.572 |
| D x HW x Sp | 0.967 | 1.284 | 0.753 | 0.451 |
| D x [+2°C+HW] x Sp | -0.360 | 1.235 | -0.291 | 0.771 |
| D x [+2°C] x Sc | 1.186 | 1.284 | 0.924 | 0.356 |
| D x HW x Sc | 0.153 | 1.253 | 0.122 | 0.903 |
| D x [+2°C+HW] x Sc | -0.191 | 1.301 | -0.147 | 0.883 |
| D x [+2°C] x 1M | 0.882 | 1.058 | 0.834 | 0.404 |
| D x HW x 1M | 1.132 | 1.064 | 1.064 | 0.287 |
| D x [+2°C+HW] x 1M | -0.167 | 1.079 | -0.154 | 0.877 |
| D x [+2°C] x 4M | -0.673 | 1.058 | -0.637 | 0.524 |
| D x HW x 4M | 0.245 | 1.053 | 0.233 | 0.816 |
| D x [+2°C+HW] x 4M | -1.114 | 1.085 | -1.026 | 0.305 |
| D x Be x 1M | 1.003 | 1.053 | 0.952 | 0.341 |
| D x Tp x 1M | 1.177 | 1.085 | 1.085 | 0.278 |
| D x Lc x 1M | 0.659 | 1.069 | 0.616 | 0.538 |
| D x Cj x 1M | 1.909 | 1.090 | 1.751 | 0.080 |
| D x To x 1M | 1.715 | 1.053 | 1.629 | 0.103 |
| D x Sp x 1M | 1.577 | 1.076 | 1.465 | 0.143 |
| D x Sc x 1M | 1.124 | 1.079 | 1.041 | 0.298 |
| D x Be x 4M | -0.348 | 1.053 | -0.330 | 0.741 |
| D x Tp x 4M | -0.882 | 1.114 | -0.792 | 0.428 |
| D x Lc x 4M | 0.015 | 1.075 | 0.014 | 0.989 |
| D x Cj x 4M | 1.144 | 1.096 | 1.044 | 0.297 |
| D x To x 4M | -0.761 | 1.053 | -0.722 | 0.470 |
| D x Sp x 4M | 0.621 | 1.064 | 0.584 | 0.559 |
| D x Sc x 4M | 0.138 | 1.075 | 0.128 | 0.898 |
| [+2°C] x Be x 1M | 0.956 | 1.053 | 0.908 | 0.364 |
| HW x Be x 1M | -1.057 | 1.058 | -0.999 | 0.318 |
| [+2°C+HW] x Be x 1M | 0.156 | 1.101 | 0.141 | 0.888 |
| [+2°C] x Tp x 1M | 0.866 | 1.072 | 0.808 | 0.419 |
| HW x Tp x 1M | 0.776 | 1.103 | 0.704 | 0.482 |
| [+2°C+HW] x Tp x 1M | 1.440 | 1.107 | 1.302 | 0.193 |
| [+2°C] x Lc x 1M | 0.557 | 1.075 | 0.519 | 0.604 |
| HW x Lc x 1M | -0.585 | 1.064 | -0.550 | 0.583 |
| [+2°C+HW] x Lc x 1M | -0.083 | 1.106 | -0.075 | 0.941 |
| [+2°C] x Cj x 1M | -0.094 | 1.058 | -0.089 | 0.929 |
| HW x Cj x 1M | 0.784 | 1.069 | 0.733 | 0.463 |
| [+2°C+HW] x Cj x 1M | 0.596 | 1.098 | 0.543 | 0.587 |
| [+2°C] x To x 1M | 1.253 | 1.058 | 1.184 | 0.237 |
| HW x To x 1M | 0.264 | 1.053 | 0.251 | 0.802 |
| [+2°C+HW] x To x 1M | 0.672 | 1.080 | 0.623 | 0.534 |
| [+2°C] x Sp x 1M | 0.941 | 1.082 | 0.869 | 0.385 |
| HW x Sp x 1M | -0.124 | 1.098 | -0.113 | 0.910 |
| [+2°C+HW] x Sp x 1M | -0.249 | 1.098 | -0.227 | 0.821 |
| [+2°C] x Sc x 1M | 0.912 | 1.058 | 0.862 | 0.389 |
| HW x Sc x 1M | -0.485 | 1.058 | -0.459 | 0.647 |
| [+2°C+HW] x Sc x 1M | -0.617 | 1.080 | -0.571 | 0.568 |
| [+2°C] x Be x 4M | -1.247 | 1.053 | -1.185 | 0.236 |
| HW x Be x 4M | -0.322 | 1.058 | -0.304 | 0.761 |
| [+2°C+HW] x Be x 4M | -1.784 | 1.096 | -1.628 | 0.104 |
| [+2°C] x Tp x 4M | -1.199 | 1.087 | -1.102 | 0.270 |
| HW x Tp x 4M | 2.033 | 1.114 | 1.826 | 0.068 |
| [+2°C+HW] x Tp x 4M | -0.820 | 1.129 | -0.726 | 0.468 |
| [+2°C] x Lc x 4M | -0.312 | 1.069 | -0.292 | 0.770 |
| HW x Lc x 4M | -0.227 | 1.064 | -0.213 | 0.831 |
| [+2°C+HW] x Lc x 4M | -1.516 | 1.111 | -1.364 | 0.172 |
| [+2°C] x Cj x 4M | -1.899 | 1.064 | -1.785 | 0.074 |
| HW x Cj x 4M | -0.475 | 1.083 | -0.439 | 0.661 |
| [+2°C+HW] x Cj x 4M | -1.869 | 1.085 | -1.722 | 0.085 |
| [+2°C] x To x 4M | -1.950 | 1.053 | -1.852 | 0.064 |
| HW x To x 4M | -0.316 | 1.053 | -0.300 | 0.764 |
| **[+2°C+HW] x To x 4M** | **-2.120** | **1.075** | **-1.973** | **0.049** |
| [+2°C] x Sp x 4M | -0.585 | 1.069 | -0.547 | 0.584 |
| HW x Sp x 4M | 0.272 | 1.085 | 0.251 | 0.802 |
| [+2°C+HW] x Sp x 4M | -1.186 | 1.085 | -1.093 | 0.275 |
| [+2°C] x Sc x 4M | -1.419 | 1.053 | -1.348 | 0.178 |
| HW x Sc x 4M | -1.031 | 1.053 | -0.979 | 0.328 |
| [+2°C+HW] x Sc x 4M | -1.942 | 1.075 | -1.807 | 0.071 |
| D x [+2°C] x Be x 1M | -2.224 | 1.512 | -1.471 | 0.141 |
| D x HW x Be x 1M | 0.399 | 1.500 | 0.266 | 0.790 |
| D x [+2°C+HW] x Be x 1M | -0.949 | 1.555 | -0.610 | 0.542 |
| D x [+2°C] x Tp x 1M | -0.768 | 1.550 | -0.496 | 0.620 |
| D x HW x Tp x 1M | -1.128 | 1.567 | -0.720 | 0.472 |
| D x [+2°C+HW] x Tp x 1M | 1.371 | 1.682 | 0.815 | 0.415 |
| D x [+2°C] x Lc x 1M | -0.463 | 1.530 | -0.302 | 0.762 |
| D x HW x Lc x 1M | 0.033 | 1.511 | 0.022 | 0.983 |
| D x [+2°C+HW] x Lc x 1M | 0.223 | 1.581 | 0.141 | 0.888 |
| D x [+2°C] x Cj x 1M | -0.771 | 1.528 | -0.505 | 0.614 |
| D x HW x Cj x 1M | -2.445 | 1.553 | -1.574 | 0.115 |
| D x [+2°C+HW] x Cj x 1M | -0.539 | 1.547 | -0.348 | 0.728 |
| D x [+2°C] x To x 1M | -0.699 | 1.496 | -0.467 | 0.640 |
| D x HW x To x 1M | -1.138 | 1.500 | -0.758 | 0.448 |
| D x [+2°C+HW] x To x 1M | -0.056 | 1.511 | -0.037 | 0.971 |
| D x [+2°C] x Sp x 1M | -1.785 | 1.518 | -1.176 | 0.239 |
| D x HW x Sp x 1M | -1.018 | 1.572 | -0.647 | 0.517 |
| D x [+2°C+HW] x Sp x 1M | 0.811 | 1.531 | 0.529 | 0.596 |
| D x [+2°C] x Sc x 1M | -2.250 | 1.555 | -1.447 | 0.148 |
| D x HW x Sc x 1M | -1.138 | 1.531 | -0.743 | 0.457 |
| D x [+2°C+HW] x Sc x 1M | -0.033 | 1.567 | -0.021 | 0.983 |
| D x [+2°C] x Be x 4M | -0.522 | 1.508 | -0.346 | 0.729 |
| D x HW x Be x 4M | -0.338 | 1.493 | -0.227 | 0.821 |
| D x [+2°C+HW] x Be x 4M | 0.525 | 1.550 | 0.339 | 0.735 |
| D x [+2°C] x Tp x 4M | -0.753 | 1.543 | -0.488 | 0.626 |
| D x HW x Tp x 4M | -1.953 | 1.562 | -1.250 | 0.211 |
| D x [+2°C+HW] x Tp x 4M | 3.154 | 1.722 | 1.831 | 0.067 |
| D x [+2°C] x Lc x 4M | 0.841 | 1.526 | 0.551 | 0.582 |
| D x HW x Lc x 4M | 0.117 | 1.505 | 0.078 | 0.938 |
| D x [+2°C+HW] x Lc x 4M | 1.135 | 1.592 | 0.713 | 0.476 |
| D x [+2°C] x Cj x 4M | 0.740 | 1.543 | 0.479 | 0.632 |
| D x HW x Cj x 4M | -1.367 | 1.552 | -0.881 | 0.378 |
| D x [+2°C+HW] x Cj x 4M | 0.984 | 1.555 | 0.633 | 0.527 |
| D x [+2°C] x To x 4M | 2.192 | 1.500 | 1.461 | 0.144 |
| D x HW x To x 4M | 0.172 | 1.489 | 0.115 | 0.908 |
| D x [+2°C+HW] x To x 4M | 2.438 | 1.520 | 1.604 | 0.109 |
| D x [+2°C] x Sp x 4M | -0.155 | 1.504 | -0.103 | 0.918 |
| D x HW x Sp x 4M | -0.094 | 1.560 | -0.060 | 0.952 |
| D x [+2°C+HW] x Sp x 4M | 0.864 | 1.523 | 0.567 | 0.571 |
| D x [+2°C] x Sc x 4M | -0.077 | 1.549 | -0.050 | 0.960 |
| D x HW x Sc x 4M | 0.284 | 1.523 | 0.186 | 0.852 |
| D x [+2°C+HW] x Sc x 4M | 0.280 | 1.570 | 0.179 | 0.858 |

**Table S7b:** Output of the general model of aboveground biomass responses to treatments and season for *Prunella vulgaris*. Explanatory variables included drought, temperature, harvest time point. Significant estimates are highlighted in bold. **Abbreviations:** D: Water treatment drought, [+2°C]: Ambient temperature plus constant 2°C warming, HW: Ambient temperature plus heatwave warming events (+10°C), [+2°C+HW]: Ambient temperature plus constant 2°C and heatwave warming. 1M: collection point at 1-month recovery after the end of drought (peak of the growing season), 4M: Data collection point at 4-months recovery after the end of drought (end of the growing season).

| Predictor | Estimate | SE | Z | p-value |
| --- | --- | --- | --- | --- |
| (Intercept) | 0.122 | 0.292 | 0.418 | 0.676 |
| D | -0.056 | 0.412 | -0.136 | 0.892 |
| [+2°C] | 0.016 | 0.412 | 0.039 | 0.969 |
| HW | 0.388 | 0.412 | 0.941 | 0.347 |
| [+2°C+HW] | -0.111 | 0.412 | -0.270 | 0.787 |
| 1M | 0.245 | 0.397 | 0.616 | 0.538 |
| 4M | 0.158 | 0.397 | 0.398 | 0.691 |
| D x [+2°C] | 0.028 | 0.583 | 0.048 | 0.962 |
| D x HW | -0.454 | 0.583 | -0.778 | 0.436 |
| D x [+2°C+HW] | 0.045 | 0.583 | 0.078 | 0.938 |
| D x 1M | 0.455 | 0.562 | 0.811 | 0.418 |
| D x 4M | 0.200 | 0.562 | 0.356 | 0.722 |
| [+2°C] x 1M | 0.071 | 0.562 | 0.127 | 0.899 |
| HW x 1M | -0.487 | 0.562 | -0.866 | 0.386 |
| [+2°C+HW] x 1M | 0.861 | 0.562 | 1.534 | 0.125 |
| [+2°C] x 4M | 0.642 | 0.562 | 1.143 | 0.253 |
| HW x 4M | -0.183 | 0.562 | -0.325 | 0.745 |
| [+2°C+HW] x 4M | 0.107 | 0.562 | 0.190 | 0.849 |
| D x [+2°C] x 1M | -0.881 | 0.794 | -1.109 | 0.267 |
| D x HW x 1M | -0.063 | 0.794 | -0.080 | 0.936 |
| **D x [+2°C+HW] x 1M** | **-1.561** | **0.794** | **-1.966** | **0.049** |
| D x [+2°C] x 4M | -0.928 | 0.794 | -1.169 | 0.243 |
| D x HW x 4M | 0.624 | 0.794 | 0.786 | 0.432 |
| D x [+2°C+HW] x 4M | -0.394 | 0.794 | -0.496 | 0.620 |

**Table S8a:** Output of the mixed-effects models testing specific SLA to treatments and season (species-specific recovery model). The model included drought, temperature, harvest time point, and plant species as explanatory variables. Due to model-fitting issues, *Prunella vulgaris* was analysed separately in an independent model (see table S8b). Significant estimates are highlighted in bold. Abbreviations: D: Water treatment drought, [+2°C]: Ambient temperature plus constant 2°C warming, HW: Ambient temperature plus heatwave warming events (+10°C), [+2°C+HW]: Ambient temperature plus constant 2°C and heatwave warming. 1M: collection point at 1-month recovery after the end of drought (peak of the growing season), 4M: Data collection point at 4-months recovery after the end of drought (end of the growing season). *Bromus erectus*: Be, *Trifolium pratense*: Tp, *Lotus corniculatus*: Lc, *Taraxacum officinale*: To, *Centaurea jacea*: Cj, *Salvia pratense*: Sp, *Solidago canadensis*: Sc. The intercept is the fast-growing species *Holcus lanatus*.

| Predictor | Estimate | SE | Z | p-value |
| --- | --- | --- | --- | --- |
| (Intercept) | 36.596 | 2.569 | 14.247 | <0.001 |
| D | 1.031 | 3.608 | 0.286 | 0.775 |
| [+2°C] | 1.938 | 3.825 | 0.507 | 0.612 |
| HW | 4.186 | 3.608 | 1.160 | 0.246 |
| [+2°C+HW] | 3.090 | 3.608 | 0.856 | 0.392 |
| Be | -12.061 | 3.563 | -3.385 | <0.001 |
| Tp | -13.672 | 3.563 | -3.838 | <0.001 |
| Lc | -12.340 | 3.563 | -3.464 | <0.001 |
| **Cj** | **-7.826** | **3.563** | **-2.197** | **0.028** |
| **To** | **-11.447** | **3.563** | **-3.213** | **0.001** |
| **Sp** | **-11.173** | **3.563** | **-3.136** | **0.002** |
| **Sc** | **-11.576** | **3.563** | **-3.249** | **0.001** |
| 1M | -4.630 | 3.091 | -1.498 | 0.134 |
| 4M | 1.495 | 3.091 | 0.484 | 0.629 |
| D x [+2°C] | -1.303 | 5.258 | -0.248 | 0.804 |
| D x HW | -8.517 | 5.103 | -1.669 | 0.095 |
| D x [+2°C+HW] | -1.488 | 5.103 | -0.292 | 0.771 |
| D x Be | -4.189 | 5.038 | -0.831 | 0.406 |
| D x Tp | 0.804 | 5.038 | 0.160 | 0.873 |
| D x Lc | 5.524 | 5.038 | 1.096 | 0.273 |
| D x Cj | -6.769 | 5.038 | -1.344 | 0.179 |
| D x To | -5.454 | 5.038 | -1.082 | 0.279 |
| D x Sp | -1.786 | 5.038 | -0.355 | 0.723 |
| D x Sc | 0.130 | 5.038 | 0.026 | 0.979 |
| [+2°C] x Be | -7.465 | 5.195 | -1.437 | 0.151 |
| HW x Be | -7.571 | 5.038 | -1.503 | 0.133 |
| [+2°C+HW] x Be | -7.565 | 5.038 | -1.501 | 0.133 |
| [+2°C] x Tp | -0.263 | 5.195 | -0.051 | 0.960 |
| HW x Tp | -0.582 | 5.038 | -0.115 | 0.908 |
| [+2°C+HW] x Tp | 0.166 | 5.196 | 0.032 | 0.974 |
| [+2°C] x Lc | 3.822 | 5.195 | 0.736 | 0.462 |
| HW x Lc | 1.566 | 5.038 | 0.311 | 0.756 |
| [+2°C+HW] x Lc | 3.090 | 5.195 | 0.595 | 0.552 |
| [+2°C] x Cj | -4.820 | 5.195 | -0.928 | 0.354 |
| HW x Cj | -7.932 | 5.038 | -1.574 | 0.115 |
| [+2°C+HW] x Cj | -6.766 | 5.038 | -1.343 | 0.179 |
| [+2°C] x To | -4.375 | 5.195 | -0.842 | 0.400 |
| HW x To | -8.134 | 5.038 | -1.614 | 0.106 |
| **[+2°C+HW] x To** | **-14.653** | **5.038** | **-2.908** | **0.004** |
| [+2°C] x Sp | -4.927 | 5.195 | -0.948 | 0.343 |
| HW x Sp | -6.281 | 5.038 | -1.247 | 0.213 |
| [+2°C+HW] x Sp | -2.445 | 5.038 | -0.485 | 0.628 |
| [+2°C] x Sc | 0.813 | 5.195 | 0.156 | 0.876 |
| HW x Sc | 1.383 | 5.038 | 0.275 | 0.784 |
| [+2°C+HW] x Sc | -3.203 | 5.038 | -0.636 | 0.525 |
| D x 1M | 4.739 | 4.363 | 1.086 | 0.277 |
| D x 4M | -1.267 | 4.404 | -0.288 | 0.774 |
| [+2°C] x 1M | -3.912 | 4.544 | -0.861 | 0.389 |
| HW x 1M | -5.021 | 4.363 | -1.151 | 0.250 |
| [+2°C+HW] x 1M | -5.343 | 4.363 | -1.224 | 0.221 |
| [+2°C] x 4M | -4.886 | 4.544 | -1.075 | 0.282 |
| HW x 4M | -7.304 | 4.404 | -1.658 | 0.097 |
| **[+2°C+HW] x 4M** | **-8.681** | **4.363** | **-1.989** | **0.047** |
| Be x 1M | -2.625 | 4.404 | -0.596 | 0.551 |
| Tp x 1M | 3.002 | 4.363 | 0.688 | 0.491 |
| Lc x 1M | 3.347 | 4.404 | 0.760 | 0.447 |
| Cj x 1M | -4.163 | 4.404 | -0.945 | 0.345 |
| To x 1M | 6.526 | 4.363 | 1.496 | 0.135 |
| Sp x 1M | -0.545 | 4.404 | -0.124 | 0.901 |
| Sc x 1M | -0.907 | 4.363 | -0.208 | 0.835 |
| Be x 4M | -3.360 | 4.363 | -0.770 | 0.441 |
| Tp x 4M | -0.708 | 4.454 | -0.159 | 0.874 |
| Lc x 4M | 1.135 | 4.363 | 0.260 | 0.795 |
| Cj x 4M | -5.877 | 4.363 | -1.347 | 0.178 |
| **To x 4M** | **8.667** | **4.363** | **1.986** | **0.047** |
| Sp x 4M | -3.351 | 4.404 | -0.761 | 0.447 |
| Sc x 4M | -6.975 | 4.363 | -1.599 | 0.110 |
| D x [+2°C] x Be | 9.350 | 7.347 | 1.273 | 0.203 |
| **D x HW x Be** | **14.388** | **7.125** | **2.019** | **0.043** |
| D x [+2°C+HW] x Be | 9.579 | 7.125 | 1.344 | 0.179 |
| D x [+2°C] x Tp | -2.582 | 7.237 | -0.357 | 0.721 |
| D x HW x Tp | 4.017 | 7.237 | 0.555 | 0.579 |
| D x [+2°C+HW] x Tp | -1.058 | 7.237 | -0.146 | 0.884 |
| D x [+2°C] x Lc | -7.327 | 7.237 | -1.012 | 0.311 |
| D x HW x Lc | 2.383 | 7.125 | 0.334 | 0.738 |
| D x [+2°C+HW] x Lc | -9.143 | 7.237 | -1.263 | 0.206 |
| D x [+2°C] x Cj | 7.844 | 7.237 | 1.084 | 0.278 |
| **D x HW x Cj** | **15.791** | **7.125** | **2.216** | **0.027** |
| D x [+2°C+HW] x Cj | 4.290 | 7.125 | 0.602 | 0.547 |
| D x [+2°C] x To | 4.727 | 7.237 | 0.653 | 0.514 |
| D x HW x To | 11.217 | 7.125 | 1.574 | 0.115 |
| D x [+2°C+HW] x To | 10.593 | 7.125 | 1.487 | 0.137 |
| D x [+2°C] x Sp | 4.551 | 7.237 | 0.629 | 0.529 |
| D x HW x Sp | 11.809 | 7.125 | 1.657 | 0.097 |
| D x [+2°C+HW] x Sp | -0.222 | 7.125 | -0.031 | 0.975 |
| D x [+2°C] x Sc | -2.058 | 7.237 | -0.284 | 0.776 |
| D x HW x Sc | 2.719 | 7.125 | 0.382 | 0.703 |
| D x [+2°C+HW] x Sc | 0.158 | 7.125 | 0.022 | 0.982 |
| D x [+2°C] x 1M | 9.517 | 6.328 | 1.504 | 0.133 |
| D x HW x 1M | 7.237 | 6.171 | 1.173 | 0.241 |
| D x [+2°C+HW] x 1M | 4.319 | 6.171 | 0.700 | 0.484 |
| D x [+2°C] x 4M | 7.619 | 6.328 | 1.204 | 0.229 |
| **D x HW x 4M** | **16.042** | **6.228** | **2.576** | **0.010** |
| D x [+2°C+HW] x 4M | 8.539 | 6.200 | 1.377 | 0.168 |
| D x Be x 1M | 0.424 | 6.200 | 0.068 | 0.945 |
| D x Tp x 1M | -4.168 | 6.171 | -0.676 | 0.499 |
| D x Lc x 1M | -0.727 | 6.200 | -0.117 | 0.907 |
| D x Cj x 1M | 5.653 | 6.228 | 0.908 | 0.364 |
| D x To x 1M | 1.466 | 6.171 | 0.238 | 0.812 |
| D x Sp x 1M | 5.694 | 6.200 | 0.919 | 0.358 |
| D x Sc x 1M | -6.678 | 6.200 | -1.077 | 0.281 |
| D x Be x 4M | 5.618 | 6.200 | 0.906 | 0.365 |
| D x Tp x 4M | -1.394 | 6.264 | -0.223 | 0.824 |
| D x Lc x 4M | -4.172 | 6.200 | -0.673 | 0.501 |
| D x Cj x 4M | 8.008 | 6.200 | 1.292 | 0.196 |
| D x To x 4M | 5.610 | 6.200 | 0.905 | 0.366 |
| D x Sp x 4M | 0.329 | 6.228 | 0.053 | 0.958 |
| D x Sc x 4M | 4.831 | 6.200 | 0.779 | 0.436 |
| **[+2°C] x Be x 1M** | **13.473** | **6.328** | **2.129** | **0.033** |
| HW x Be x 1M | 7.414 | 6.200 | 1.196 | 0.232 |
| **[+2°C+HW] x Be x 1M** | **14.517** | **6.200** | **2.342** | **0.019** |
| [+2°C] x Tp x 1M | 0.886 | 6.329 | 0.140 | 0.889 |
| HW x Tp x 1M | -0.928 | 6.200 | -0.150 | 0.881 |
| [+2°C+HW] x Tp x 1M | 0.634 | 6.326 | 0.100 | 0.920 |
| [+2°C] x Lc x 1M | -5.130 | 6.328 | -0.811 | 0.418 |
| HW x Lc x 1M | -0.970 | 6.200 | -0.156 | 0.876 |
| [+2°C+HW] x Lc x 1M | -3.479 | 6.328 | -0.550 | 0.582 |
| [+2°C] x Cj x 1M | 9.578 | 6.328 | 1.514 | 0.130 |
| **HW x Cj x 1M** | **12.343** | **6.264** | **1.970** | **0.049** |
| [+2°C+HW] x Cj x 1M | 7.205 | 6.200 | 1.162 | 0.245 |
| [+2°C] x To x 1M | 2.346 | 6.300 | 0.372 | 0.710 |
| HW x To x 1M | 5.372 | 6.171 | 0.871 | 0.384 |
| **[+2°C+HW] x To x 1M** | **12.351** | **6.171** | **2.002** | **0.045** |
| [+2°C] x Sp x 1M | 7.137 | 6.328 | 1.128 | 0.259 |
| HW x Sp x 1M | 6.519 | 6.200 | 1.052 | 0.293 |
| [+2°C+HW] x Sp x 1M | 4.534 | 6.200 | 0.731 | 0.465 |
| [+2°C] x Sc x 1M | 2.073 | 6.300 | 0.329 | 0.742 |
| HW x Sc x 1M | -3.297 | 6.235 | -0.529 | 0.597 |
| [+2°C+HW] x Sc x 1M | 8.563 | 6.171 | 1.388 | 0.165 |
| [+2°C] x Be x 4M | 7.517 | 6.300 | 1.193 | 0.233 |
| HW x Be x 4M | 9.799 | 6.200 | 1.581 | 0.114 |
| **[+2°C+HW] x Be x 4M** | **12.866** | **6.171** | **2.085** | **0.037** |
| [+2°C] x Tp x 4M | 3.174 | 6.363 | 0.499 | 0.618 |
| HW x Tp x 4M | 5.466 | 6.293 | 0.869 | 0.385 |
| [+2°C+HW] x Tp x 4M | 4.463 | 6.389 | 0.698 | 0.485 |
| [+2°C] x Lc x 4M | 0.955 | 6.300 | 0.152 | 0.879 |
| HW x Lc x 4M | 1.415 | 6.200 | 0.228 | 0.819 |
| [+2°C+HW] x Lc x 4M | 1.423 | 6.300 | 0.226 | 0.821 |
| [+2°C] x Cj x 4M | 5.775 | 6.300 | 0.917 | 0.359 |
| HW x Cj x 4M | 10.319 | 6.228 | 1.657 | 0.098 |
| [+2°C+HW] x Cj x 4M | 10.296 | 6.171 | 1.669 | 0.095 |
| [+2°C] x To x 4M | 0.714 | 6.300 | 0.113 | 0.910 |
| HW x To x 4M | 5.656 | 6.200 | 0.912 | 0.362 |
| **[+2°C+HW] x To x 4M** | **14.448** | **6.171** | **2.341** | **0.019** |
| [+2°C] x Sp x 4M | 8.031 | 6.328 | 1.269 | 0.204 |
| HW x Sp x 4M | 10.529 | 6.228 | 1.690 | 0.091 |
| [+2°C+HW] x Sp x 4M | 4.567 | 6.200 | 0.737 | 0.461 |
| [+2°C] x Sc x 4M | 4.511 | 6.300 | 0.716 | 0.474 |
| HW x Sc x 4M | 1.003 | 6.200 | 0.162 | 0.872 |
| [+2°C+HW] x Sc x 4M | 9.253 | 6.171 | 1.500 | 0.134 |
| **D x [+2°C] x Be x 1M** | **-18.440** | **8.969** | **-2.056** | **0.040** |
| D x HW x Be x 1M | -10.135 | 8.747 | -1.159 | 0.247 |
| D x [+2°C+HW] x Be x 1M | -13.546 | 8.747 | -1.549 | 0.121 |
| D x [+2°C] x Tp x 1M | -4.542 | 8.860 | -0.513 | 0.608 |
| D x HW x Tp x 1M | -0.334 | 8.838 | -0.038 | 0.970 |
| D x [+2°C+HW] x Tp x 1M | -1.509 | 8.837 | -0.171 | 0.864 |
| D x [+2°C] x Lc x 1M | -1.578 | 8.859 | -0.178 | 0.859 |
| D x HW x Lc x 1M | -12.609 | 8.747 | -1.441 | 0.149 |
| D x [+2°C+HW] x Lc x 1M | 1.684 | 8.838 | 0.191 | 0.849 |
| D x [+2°C] x Cj x 1M | -11.661 | 8.924 | -1.307 | 0.191 |
| **D x HW x Cj x 1M** | **-18.692** | **8.858** | **-2.110** | **0.035** |
| D x [+2°C+HW] x Cj x 1M | -2.835 | 8.767 | -0.323 | 0.746 |
| D x [+2°C] x To x 1M | -12.523 | 8.838 | -1.417 | 0.157 |
| D x HW x To x 1M | -15.466 | 8.727 | -1.772 | 0.076 |
| D x [+2°C+HW] x To x 1M | -12.788 | 8.727 | -1.465 | 0.143 |
| D x [+2°C] x Sp x 1M | -16.993 | 8.859 | -1.918 | 0.055 |
| D x HW x Sp x 1M | -13.509 | 8.747 | -1.544 | 0.122 |
| D x [+2°C+HW] x Sp x 1M | -6.452 | 8.747 | -0.738 | 0.461 |
| D x [+2°C] x Sc x 1M | -3.453 | 8.859 | -0.390 | 0.697 |
| D x HW x Sc x 1M | 2.821 | 8.793 | 0.321 | 0.748 |
| D x [+2°C+HW] x Sc x 1M | -6.368 | 8.768 | -0.726 | 0.468 |
| D x [+2°C] x Be x 4M | -15.619 | 8.929 | -1.749 | 0.080 |
| **D x HW x Be x 4M** | **-23.612** | **8.768** | **-2.693** | **0.007** |
| **D x [+2°C+HW] x Be x 4M** | **-18.311** | **8.747** | **-2.093** | **0.036** |
| D x [+2°C] x Tp x 4M | -2.080 | 8.884 | -0.234 | 0.815 |
| D x HW x Tp x 4M | -13.588 | 8.945 | -1.519 | 0.129 |
| D x [+2°C+HW] x Tp x 4M | -6.335 | 8.923 | -0.710 | 0.478 |
| D x [+2°C] x Lc x 4M | 0.501 | 8.838 | 0.057 | 0.955 |
| D x HW x Lc x 4M | -9.739 | 8.768 | -1.111 | 0.267 |
| D x [+2°C+HW] x Lc x 4M | 0.474 | 8.838 | 0.054 | 0.957 |
| D x [+2°C] x Cj x 4M | -13.067 | 8.838 | -1.478 | 0.139 |
| **D x HW x Cj x 4M** | **-22.426** | **8.808** | **-2.546** | **0.011** |
| D x [+2°C+HW] x Cj x 4M | -12.900 | 8.747 | -1.475 | 0.140 |
| D x [+2°C] x To x 4M | -5.954 | 8.838 | -0.674 | 0.501 |
| **D x HW x To x 4M** | **-18.391** | **8.768** | **-2.098** | **0.036** |
| **D x [+2°C+HW] x To x 4M** | **-19.095** | **8.747** | **-2.183** | **0.029** |
| D x [+2°C] x Sp x 4M | -8.639 | 8.859 | -0.975 | 0.329 |
| **D x HW x Sp x 4M** | **-20.249** | **8.788** | **-2.304** | **0.021** |
| D x [+2°C+HW] x Sp x 4M | -4.396 | 8.768 | -0.501 | 0.616 |
| D x [+2°C] x Sc x 4M | -10.926 | 8.838 | -1.236 | 0.216 |
| D x HW x Sc x 4M | -11.194 | 8.768 | -1.277 | 0.202 |
| D x [+2°C+HW] x Sc x 4M | -8.966 | 8.747 | -1.025 | 0.305 |

**Table S8b:** Output of the general model of the SLA responses to the treatment and season according to the plants' species *Prunella vulgaris*. The model uses the above-ground biomass as the response variable, and as explanatory variables, the interactive effect of Drought, Temperature, harvest time points. Significant estimates are highlighted in bold. **Abbreviations:** D: Water treatment drought, [+2°C]: Ambient temperature plus constant 2°C warming, HW: Ambient temperature plus heatwave warming events (+10°C), [+2°C+HW]: Ambient temperature plus constant 2°C and heatwave warming. 1M: collection point at 1-month recovery after the end of drought (peak of the growing season), 4M: Data collection point at 4-months recovery after the end of drought (end of the growing season).

| Predictor | Estimate | SE | Z | p-value |
| --- | --- | --- | --- | --- |
| (Intercept) | 10.827 | 0.077 | 140.505 | <0.001 |
| D | -0.078 | 0.115 | -0.676 | 0.499 |
| [+2°C] | 0.042 | 0.109 | 0.387 | 0.698 |
| HW | -0.147 | 0.109 | -1.346 | 0.178 |
| [+2°C+HW] | -0.170 | 0.109 | -1.557 | 0.119 |
| 1M | -0.415 | 0.101 | -4.120 | <0.001 |
| 4M | -0.614 | 0.087 | -7.036 | <0.001 |
| D x [+2°C] | -0.014 | 0.171 | -0.085 | 0.933 |
| **D x HW** | **0.405** | **0.185** | **2.192** | **0.028** |
| D x [+2°C+HW] | 0.257 | 0.185 | 1.391 | 0.164 |
| D x 1M | 0.195 | 0.147 | 1.320 | 0.187 |
| D x 4M | 0.213 | 0.130 | 1.634 | 0.102 |
| [+2°C] x 1M | -0.119 | 0.142 | -0.837 | 0.402 |
| HW x 1M | -0.049 | 0.147 | -0.332 | 0.740 |
| [+2°C+HW] x 1M | -0.150 | 0.147 | -1.015 | 0.310 |
| [+2°C] x 4M | 0.064 | 0.123 | 0.522 | 0.601 |
| HW x 4M | 0.216 | 0.123 | 1.749 | 0.080 |
| **[+2°C+HW] x 4M** | **0.395** | **0.123** | **3.202** | **0.001** |
| D x [+2°C] x 1M | 0.322 | 0.231 | 1.394 | 0.163 |
| D x HW x 1M | 0.079 | 0.242 | 0.325 | 0.745 |
| D x [+2°C+HW] x 1M | 0.115 | 0.270 | 0.427 | 0.669 |
| D x [+2°C] x 4M | -0.147 | 0.198 | -0.739 | 0.460 |
| **D x HW x 4M** | **-0.490** | **0.209** | **-2.352** | **0.019** |
| **D x [+2°C+HW] x 4M** | **-0.696** | **0.227** | **-3.060** | **0.002** |

**Table S9a:** Output of the mixed-effects models testing specific LDMC responses to treatments and season (species-specific recovery model). The model included drought, temperature, harvest time point, and plant species as explanatory variables. Due to model-fitting issues, *Prunella vulgaris* was analysed separately in an independent model (see table S9b). Significant estimates are highlighted in bold. Abbreviations: D: Water treatment drought, [+2°C]: Ambient temperature plus constant 2°C warming, HW: Ambient temperature plus heatwave warming events (+10°C), [+2°C+HW]: Ambient temperature plus constant 2°C and heatwave warming. 1M: collection point at 1-month recovery after the end of drought (peak of the growing season), 4M: Data collection point at 4-months recovery after the end of drought (end of the growing season). *Bromus erectus*: Be, *Trifolium pratense*: Tp, *Lotus corniculatus*: Lc, *Taraxacum officinale*: To, *Centaurea jacea*: Cj, *Salvia pratense*: Sp, *Solidago canadensis*: Sc. The intercept is the fast-growing species *Holcus lanatus*.

| Predictor | Estimate | SE | Z | p-value |
| --- | --- | --- | --- | --- |
| (Intercept) | 229.031 | 15.492 | 14.784 | <0.001 |
| D | -11.506 | 21.809 | -0.528 | 0.598 |
| [+2°C] | 11.901 | 23.125 | 0.515 | 0.607 |
| HW | -4.768 | 21.809 | -0.219 | 0.827 |
| [+2°C+HW] | -10.775 | 21.809 | -0.494 | 0.621 |
| **Be** | **50.524** | **21.642** | **2.334** | **0.020** |
| Tp | -14.940 | 21.642 | -0.690 | 0.490 |
| **Lc** | **-57.015** | **21.642** | **-2.634** | **0.008** |
| Cj | **-60.053** | **21.642** | **-2.775** | **0.006** |
| To | -82.989 | 21.642 | -3.835 | <0.001 |
| Sp | -13.057 | 21.642 | -0.603 | 0.546 |
| Sc | **-65.388** | **21.642** | **-3.021** | **0.003** |
| 1M | -18.374 | 18.783 | -0.978 | 0.328 |
| 4M | -18.337 | 18.783 | -0.976 | 0.329 |
| D x [+2°C] | 35.286 | 31.787 | 1.110 | 0.267 |
| D x HW | 26.331 | 30.843 | 0.854 | 0.393 |
| D x [+2°C+HW] | -3.736 | 30.843 | -0.121 | 0.904 |
| D x Be | 2.316 | 30.607 | 0.076 | 0.940 |
| D x Tp | 22.718 | 30.607 | 0.742 | 0.458 |
| D x Lc | 22.894 | 30.607 | 0.748 | 0.454 |
| D x Cj | 28.885 | 30.607 | 0.944 | 0.345 |
| D x To | 10.863 | 30.607 | 0.355 | 0.723 |
| D x Sp | 12.830 | 30.607 | 0.419 | 0.675 |
| D x Sc | 13.922 | 30.607 | 0.455 | 0.649 |
| [+2°C] x Be | 18.950 | 31.558 | 0.600 | 0.548 |
| HW x Be | 3.278 | 30.607 | 0.107 | 0.915 |
| [+2°C+HW] x Be | 7.535 | 30.607 | 0.246 | 0.806 |
| [+2°C] x Tp | 2.011 | 31.558 | 0.064 | 0.949 |
| HW x Tp | 16.266 | 30.607 | 0.531 | 0.595 |
| [+2°C+HW] x Tp | 27.486 | 31.558 | 0.871 | 0.384 |
| [+2°C] x Lc | 5.745 | 31.558 | 0.182 | 0.856 |
| HW x Lc | 14.870 | 30.607 | 0.486 | 0.627 |
| [+2°C+HW] x Lc | 17.249 | 31.559 | 0.547 | 0.585 |
| [+2°C] x Cj | 4.406 | 31.558 | 0.140 | 0.889 |
| HW x Cj | 39.840 | 30.607 | 1.302 | 0.193 |
| [+2°C+HW] x Cj | 30.624 | 30.607 | 1.001 | 0.317 |
| [+2°C] x To | -20.111 | 31.558 | -0.637 | 0.524 |
| HW x To | 4.824 | 30.607 | 0.158 | 0.875 |
| [+2°C+HW] x To | 24.483 | 30.607 | 0.800 | 0.424 |
| [+2°C] x Sp | -2.004 | 31.558 | -0.063 | 0.949 |
| HW x Sp | 21.167 | 30.607 | 0.692 | 0.489 |
| [+2°C+HW] x Sp | 10.605 | 30.607 | 0.347 | 0.729 |
| [+2°C] x Sc | -11.194 | 31.558 | -0.355 | 0.723 |
| HW x Sc | -0.441 | 30.607 | -0.014 | 0.989 |
| [+2°C+HW] x Sc | 9.011 | 30.607 | 0.294 | 0.768 |
| D x 1M | -15.308 | 26.506 | -0.578 | 0.564 |
| D x 4M | 3.920 | 26.506 | 0.148 | 0.882 |
| [+2°C] x 1M | -1.359 | 27.599 | -0.049 | 0.961 |
| HW x 1M | 19.836 | 26.506 | 0.748 | 0.454 |
| [+2°C+HW] x 1M | 32.099 | 26.506 | 1.211 | 0.226 |
| [+2°C] x 4M | -5.169 | 27.599 | -0.187 | 0.851 |
| HW x 4M | 9.665 | 26.753 | 0.361 | 0.718 |
| [+2°C+HW] x 4M | 21.781 | 26.506 | 0.822 | 0.411 |
| Be x 1M | 31.501 | 26.753 | 1.177 | 0.239 |
| Tp x 1M | -0.476 | 26.506 | -0.018 | 0.986 |
| **Lc x 1M** | **75.149** | **26.753** | **2.809** | **0.005** |
| **Cj x 1M** | **80.057** | **26.753** | **2.992** | **0.003** |
| To x 1M | 36.382 | 26.506 | 1.373 | 0.170 |
| Sp x 1M | -7.151 | 26.754 | -0.267 | 0.789 |
| Sc x 1M | 144.578 | 26.506 | 5.454 | <0.001 |
| Be x 4M | 0.002 | 26.506 | 8.19e-05 | 1.000 |
| Tp x 4M | -3.609 | 27.059 | -0.133 | 0.894 |
| Lc x 4M | 50.094 | 26.506 | 1.890 | 0.059 |
| Cj x 4M | 0.221 | 26.506 | 0.008 | 0.993 |
| To x 4M | 0.382 | 26.506 | 0.014 | 0.988 |
| Sp x 4M | -38.390 | 26.753 | -1.435 | 0.151 |
| Sc x 4M | 152.560 | 26.506 | 5.756 | <0.001 |
| D x [+2°C] x Be | -60.450 | 43.962 | -1.375 | 0.169 |
| D x HW x Be | -44.915 | 43.284 | -1.038 | 0.299 |
| D x [+2°C+HW] x Be | 23.875 | 43.284 | 0.552 | 0.581 |
| D x [+2°C] x Tp | -37.184 | 43.962 | -0.846 | 0.398 |
| D x HW x Tp | -54.055 | 43.963 | -1.230 | 0.219 |
| D x [+2°C+HW] x Tp | -6.068 | 43.962 | -0.138 | 0.890 |
| D x [+2°C] x Lc | -49.540 | 43.962 | -1.127 | 0.260 |
| D x HW x Lc | -36.947 | 43.284 | -0.854 | 0.393 |
| D x [+2°C+HW] x Lc | 5.473 | 43.963 | 0.124 | 0.901 |
| D x [+2°C] x Cj | -62.441 | 43.962 | -1.420 | 0.156 |
| D x HW x Cj | -67.692 | 43.284 | -1.564 | 0.118 |
| D x [+2°C+HW] x Cj | -17.035 | 43.284 | -0.394 | 0.694 |
| D x [+2°C] x To | -27.866 | 43.962 | -0.634 | 0.526 |
| D x HW x To | -28.766 | 43.284 | -0.665 | 0.506 |
| D x [+2°C+HW] x To | 2.921 | 43.284 | 0.067 | 0.946 |
| D x [+2°C] x Sp | -40.208 | 43.962 | -0.915 | 0.360 |
| D x HW x Sp | -36.546 | 43.284 | -0.844 | 0.398 |
| D x [+2°C+HW] x Sp | 7.362 | 43.284 | 0.170 | 0.865 |
| D x [+2°C] x Sc | -30.738 | 43.962 | -0.699 | 0.484 |
| D x HW x Sc | -22.465 | 43.284 | -0.519 | 0.604 |
| D x [+2°C+HW] x Sc | 60.083 | 43.284 | 1.388 | 0.165 |
| D x [+2°C] x 1M | -53.103 | 38.438 | -1.382 | 0.167 |
| D x HW x 1M | -35.867 | 37.485 | -0.957 | 0.339 |
| D x [+2°C+HW] x 1M | -28.746 | 37.485 | -0.767 | 0.443 |
| D x [+2°C] x 4M | -43.392 | 38.266 | -1.134 | 0.257 |
| D x HW x 4M | -26.993 | 37.660 | -0.717 | 0.474 |
| D x [+2°C+HW] x 4M | -2.831 | 37.485 | -0.076 | 0.940 |
| D x Be x 1M | 17.095 | 37.661 | 0.454 | 0.650 |
| D x Tp x 1M | -13.119 | 37.485 | -0.350 | 0.726 |
| D x Lc x 1M | -2.406 | 37.660 | -0.064 | 0.949 |
| D x Cj x 1M | -66.361 | 37.834 | -1.754 | 0.079 |
| D x To x 1M | -14.007 | 37.485 | -0.374 | 0.709 |
| D x Sp x 1M | -40.519 | 37.661 | -1.076 | 0.282 |
| D x Sc x 1M | 1.590 | 37.660 | 0.042 | 0.966 |
| D x Be x 4M | 7.732 | 37.485 | 0.206 | 0.837 |
| D x Tp x 4M | -13.219 | 37.878 | -0.349 | 0.727 |
| D x Lc x 4M | -20.073 | 37.485 | -0.535 | 0.592 |
| D x Cj x 4M | -11.297 | 37.485 | -0.301 | 0.763 |
| D x To x 4M | -2.792 | 37.485 | -0.074 | 0.941 |
| D x Sp x 4M | -0.766 | 37.660 | -0.020 | 0.984 |
| D x Sc x 4M | -62.595 | 37.485 | -1.670 | 0.095 |
| [+2°C] x Be x 1M | -26.214 | 38.438 | -0.682 | 0.495 |
| HW x Be x 1M | -0.665 | 37.661 | -0.018 | 0.986 |
| [+2°C+HW] x Be x 1M | -43.523 | 37.661 | -1.156 | 0.248 |
| [+2°C] x Tp x 1M | 12.299 | 38.266 | 0.321 | 0.748 |
| HW x Tp x 1M | -0.835 | 37.660 | -0.022 | 0.982 |
| [+2°C+HW] x Tp x 1M | -33.139 | 38.431 | -0.862 | 0.389 |
| [+2°C] x Lc x 1M | 22.728 | 38.438 | 0.591 | 0.554 |
| HW x Lc x 1M | -23.986 | 37.660 | -0.637 | 0.524 |
| [+2°C+HW] x Lc x 1M | -33.320 | 38.439 | -0.867 | 0.386 |
| [+2°C] x Cj x 1M | -58.855 | 38.438 | -1.531 | 0.126 |
| **HW x Cj x 1M** | **-85.910** | **38.051** | **-2.258** | **0.024** |
| [+2°C+HW] x Cj x 1M | -49.125 | 37.661 | -1.304 | 0.192 |
| [+2°C] x To x 1M | 22.997 | 38.266 | 0.601 | 0.548 |
| HW x To x 1M | -16.973 | 37.485 | -0.453 | 0.651 |
| [+2°C+HW] x To x 1M | -43.372 | 37.485 | -1.157 | 0.247 |
| [+2°C] x Sp x 1M | 4.289 | 38.437 | 0.112 | 0.911 |
| HW x Sp x 1M | -23.088 | 37.661 | -0.613 | 0.540 |
| [+2°C+HW] x Sp x 1M | -38.959 | 37.661 | -1.034 | 0.301 |
| [+2°C] x Sc x 1M | -11.693 | 38.266 | -0.306 | 0.760 |
| HW x Sc x 1M | -8.003 | 37.878 | -0.211 | 0.833 |
| [+2°C+HW] x Sc x 1M | -8.335 | 37.485 | -0.222 | 0.824 |
| [+2°C] x Be x 4M | -5.641 | 38.266 | -0.147 | 0.883 |
| HW x Be x 4M | -4.105 | 37.660 | -0.109 | 0.913 |
| [+2°C+HW] x Be x 4M | -19.309 | 37.485 | -0.515 | 0.606 |
| [+2°C] x Tp x 4M | -4.923 | 38.652 | -0.127 | 0.899 |
| HW x Tp x 4M | -26.746 | 38.224 | -0.700 | 0.484 |
| [+2°C+HW] x Tp x 4M | -29.937 | 38.813 | -0.771 | 0.441 |
| [+2°C] x Lc x 4M | -18.875 | 38.266 | -0.493 | 0.622 |
| HW x Lc x 4M | -20.649 | 37.660 | -0.548 | 0.583 |
| [+2°C+HW] x Lc x 4M | -23.971 | 38.267 | -0.626 | 0.531 |
| [+2°C] x Cj x 4M | 3.504 | 38.266 | 0.092 | 0.927 |
| HW x Cj x 4M | -43.105 | 37.831 | -1.139 | 0.255 |
| [+2°C+HW] x Cj x 4M | -28.613 | 37.485 | -0.763 | 0.445 |
| [+2°C] x To x 4M | 21.935 | 38.266 | 0.573 | 0.566 |
| HW x To x 4M | 3.863 | 37.660 | 0.103 | 0.918 |
| [+2°C+HW] x To x 4M | -27.508 | 37.485 | -0.734 | 0.463 |
| [+2°C] x Sp x 4M | -6.448 | 38.438 | -0.168 | 0.867 |
| HW x Sp x 4M | -30.619 | 37.835 | -0.809 | 0.418 |
| [+2°C+HW] x Sp x 4M | -19.482 | 37.660 | -0.517 | 0.605 |
| [+2°C] x Sc x 4M | -20.167 | 38.266 | -0.527 | 0.598 |
| HW x Sc x 4M | -7.412 | 37.660 | -0.197 | 0.844 |
| [+2°C+HW] x Sc x 4M | -30.020 | 37.485 | -0.801 | 0.423 |
| D x [+2°C] x Be x 1M | 40.878 | 53.935 | 0.758 | 0.449 |
| D x HW x Be x 1M | 16.385 | 53.136 | 0.308 | 0.758 |
| D x [+2°C+HW] x Be x 1M | -8.156 | 53.136 | -0.153 | 0.878 |
| D x [+2°C] x Tp x 1M | 29.652 | 53.690 | 0.552 | 0.581 |
| D x HW x Tp x 1M | 48.558 | 53.691 | 0.904 | 0.366 |
| D x [+2°C+HW] x Tp x 1M | 26.723 | 53.685 | 0.498 | 0.619 |
| D x [+2°C] x Lc x 1M | -5.178 | 53.813 | -0.096 | 0.923 |
| D x HW x Lc x 1M | 27.055 | 53.136 | 0.509 | 0.611 |
| D x [+2°C+HW] x Lc x 1M | 16.206 | 53.691 | 0.302 | 0.763 |
| D x [+2°C] x Cj x 1M | 83.766 | 54.208 | 1.545 | 0.122 |
| D x HW x Cj x 1M | 103.849 | 53.811 | 1.930 | 0.054 |
| D x [+2°C+HW] x Cj x 1M | 37.751 | 53.259 | 0.709 | 0.478 |
| D x [+2°C] x To x 1M | 43.748 | 53.690 | 0.815 | 0.415 |
| D x HW x To x 1M | 45.121 | 53.012 | 0.851 | 0.395 |
| D x [+2°C+HW] x To x 1M | 30.886 | 53.012 | 0.583 | 0.560 |
| D x [+2°C] x Sp x 1M | 37.479 | 53.811 | 0.696 | 0.486 |
| D x HW x Sp x 1M | 32.363 | 53.137 | 0.609 | 0.542 |
| D x [+2°C+HW] x Sp x 1M | 31.632 | 53.137 | 0.595 | 0.552 |
| D x [+2°C] x Sc x 1M | 52.924 | 53.812 | 0.983 | 0.325 |
| D x HW x Sc x 1M | 11.485 | 53.415 | 0.215 | 0.830 |
| D x [+2°C+HW] x Sc x 1M | -43.548 | 53.260 | -0.818 | 0.414 |
| D x [+2°C] x Be x 4M | 78.939 | 53.567 | 1.474 | 0.141 |
| D x HW x Be x 4M | 41.529 | 53.136 | 0.782 | 0.434 |
| D x [+2°C+HW] x Be x 4M | -15.912 | 53.012 | -0.300 | 0.764 |
| D x [+2°C] x Tp x 4M | 32.433 | 53.843 | 0.602 | 0.547 |
| D x HW x Tp x 4M | 56.495 | 54.214 | 1.042 | 0.297 |
| D x [+2°C+HW] x Tp x 4M | 12.926 | 54.082 | 0.239 | 0.811 |
| D x [+2°C] x Lc x 4M | 66.442 | 53.567 | 1.240 | 0.215 |
| D x HW x Lc x 4M | 47.574 | 53.136 | 0.895 | 0.371 |
| D x [+2°C+HW] x Lc x 4M | 30.150 | 53.568 | 0.563 | 0.574 |
| D x [+2°C] x Cj x 4M | 47.633 | 53.567 | 0.889 | 0.374 |
| D x HW x Cj x 4M | 59.430 | 53.381 | 1.113 | 0.266 |
| D x [+2°C+HW] x Cj x 4M | 25.331 | 53.012 | 0.478 | 0.633 |
| D x [+2°C] x To x 4M | 28.420 | 53.567 | 0.531 | 0.596 |
| D x HW x To x 4M | 18.380 | 53.136 | 0.346 | 0.729 |
| D x [+2°C+HW] x To x 4M | 8.492 | 53.012 | 0.160 | 0.873 |
| D x [+2°C] x Sp x 4M | 57.506 | 53.690 | 1.071 | 0.284 |
| D x HW x Sp x 4M | 31.693 | 53.260 | 0.595 | 0.552 |
| D x [+2°C+HW] x Sp x 4M | 0.002 | 53.136 | 3.44e-05 | 1.000 |
| **D x [+2°C] x Sc x 4M** | **109.675** | **53.567** | **2.047** | **0.041** |
| D x HW x Sc x 4M | 58.607 | 53.136 | 1.103 | 0.270 |
| D x [+2°C+HW] x Sc x 4M | -10.278 | 53.012 | -0.194 | 0.846 |

**Table S9b:** Output of the general model of the LDMC responses to the treatment and season according to the plants' species *Prunella vulgaris*. The model uses the above-ground biomass as the response variable, and as explanatory variables, the interactive effect of Drought, Temperature, harvest time points. Significant estimates are highlighted in bold. Abbreviations: D: Water treatment drought, [+2°C]: Ambient temperature plus constant 2°C warming, HW: Ambient temperature plus heatwave warming events (+10°C), [+2°C+HW]: Ambient temperature plus constant 2°C and heatwave warming. 1M: collection point at 1-month recovery after the end of drought (peak of the growing season), 4M: Data collection point at 4-months recovery after the end of drought (end of the growing season).

| Predictor | Estimate | SE | Z | p-value |
| --- | --- | --- | --- | --- |
| (Intercept) | 11.935 | 0.122 | 97.694 | <0.001 |
| D | -0.007 | 0.183 | -0.036 | 0.972 |
| [+2°C] | 0.081 | 0.173 | 0.471 | 0.637 |
| HW | 0.089 | 0.173 | 0.517 | 0.605 |
| [+2°C+HW] | 0.118 | 0.173 | 0.682 | 0.495 |
| 1M | 0.265 | 0.173 | 1.533 | 0.125 |
| 4M | 0.285 | 0.150 | 1.903 | 0.057 |
| D x [+2°C] | -0.151 | 0.271 | -0.556 | 0.578 |
| D x HW | -0.308 | 0.293 | -1.050 | 0.294 |
| D x [+2°C+HW] | -0.042 | 0.293 | -0.142 | 0.887 |
| D x 1M | -0.085 | 0.252 | -0.337 | 0.736 |
| D x 4M | -0.087 | 0.222 | -0.391 | 0.696 |
| [+2°C] x 1M | -0.120 | 0.244 | -0.493 | 0.622 |
| HW x 1M | -0.037 | 0.252 | -0.146 | 0.884 |
| [+2°C+HW] x 1M | -0.008 | 0.252 | -0.032 | 0.975 |
| [+2°C] x 4M | -0.174 | 0.212 | -0.823 | 0.410 |
| HW x 4M | -0.216 | 0.212 | -1.021 | 0.307 |
| [+2°C+HW] x 4M | -0.189 | 0.214 | -0.887 | 0.375 |
| D x [+2°C] x 1M | -0.261 | 0.394 | -0.661 | 0.508 |
| D x HW x 1M | -0.185 | 0.414 | -0.445 | 0.656 |
| D x [+2°C+HW] x 1M | -0.302 | 0.457 | -0.660 | 0.509 |
| D x [+2°C] x 4M | 0.608 | 0.339 | 1.790 | 0.073 |
| D x HW x 4M | 0.393 | 0.357 | 1.100 | 0.272 |
| D x [+2°C+HW] x 4M | 0.167 | 0.384 | 0.435 | 0.664 |

**Table S10a:** Output of the mixed-effects models testing specific chlorophyll content responses to treatments and season (species-specific recovery model). The model included drought, temperature, harvest time point, and plant species as explanatory variables. Due to model-fitting issues, *Prunella vulgaris* was analysed separately in an independent model (see table S10b). Significant estimates are highlighted in bold. Abbreviations: D: Water treatment drought, [+2°C]: Ambient temperature plus constant 2°C warming, HW: Ambient temperature plus heatwave warming events (+10°C), [+2°C+HW]: Ambient temperature plus constant 2°C and heatwave warming. 1M: collection point at 1-month recovery after the end of drought (peak of the growing season), 4M: Data collection point at 4-months recovery after the end of drought (end of the growing season). *Bromus erectus*: Be, *Trifolium pratense*: Tp, *Lotus corniculatus*: Lc, *Taraxacum officinale*: To, *Centaurea jacea*: Cj, *Salvia pratense*: Sp, *Solidago canadensis*: Sc. The intercept is the fast-growing species *Holcus lanatus*.

| Predictor | Estimate | SE | Z | p-value |
| --- | --- | --- | --- | --- |
| (Intercept) | 10.561 | 0.182 | 58.137 | <0.001 |
| D | -0.048 | 0.108 | -0.446 | 0.656 |
| [+2°C] | -0.191 | 0.108 | -1.775 | 0.076 |
| HW | -0.110 | 0.108 | -1.019 | 0.308 |
| [+2°C+HW] | -0.179 | 0.108 | -1.657 | 0.098 |
| Be | -0.050 | 0.107 | -0.461 | 0.645 |
| Tp | 0.385 | 0.107 | 3.585 | <0.001 |
| **Lc** | **0.220** | **0.107** | **2.050** | **0.040** |
| Cj | 0.087 | 0.107 | 0.814 | 0.416 |
| To | -0.072 | 0.107 | -0.667 | 0.505 |
| Sp | -0.020 | 0.107 | -0.185 | 0.853 |
| Sc | -0.112 | 0.107 | -1.040 | 0.298 |
| **1M** | **-0.247** | **0.093** | **-2.645** | **0.008** |
| **4M** | **-0.206** | **0.094** | **-2.188** | **0.029** |
| D x [+2°C] | 0.053 | 0.152 | 0.346 | 0.729 |
| D x HW | -0.037 | 0.152 | -0.240 | 0.810 |
| D x [+2°C+HW] | 0.219 | 0.152 | 1.434 | 0.152 |
| D x Be | -0.177 | 0.152 | -1.165 | 0.244 |
| D x Tp | 0.014 | 0.152 | 0.089 | 0.929 |
| D x Lc | 0.057 | 0.152 | 0.376 | 0.707 |
| D x Cj | 0.025 | 0.152 | 0.167 | 0.868 |
| D x To | 0.029 | 0.152 | 0.192 | 0.847 |
| D x Sp | -0.026 | 0.152 | -0.170 | 0.865 |
| D x Sc | 0.139 | 0.152 | 0.912 | 0.362 |
| [+2°C] x Be | 0.141 | 0.152 | 0.929 | 0.353 |
| HW x Be | -0.049 | 0.152 | -0.323 | 0.747 |
| [+2°C+HW] x Be | 0.108 | 0.152 | 0.712 | 0.477 |
| [+2°C] x Tp | 0.039 | 0.152 | 0.257 | 0.797 |
| HW x Tp | 0.149 | 0.152 | 0.979 | 0.327 |
| [+2°C+HW] x Tp | 0.107 | 0.157 | 0.683 | 0.495 |
| [+2°C] x Lc | 0.147 | 0.152 | 0.970 | 0.332 |
| HW x Lc | 0.193 | 0.152 | 1.270 | 0.204 |
| [+2°C+HW] x Lc | 0.211 | 0.152 | 1.386 | 0.166 |
| [+2°C] x Cj | 0.185 | 0.152 | 1.218 | 0.223 |
| HW x Cj | 0.133 | 0.152 | 0.873 | 0.383 |
| [+2°C+HW] x Cj | 0.171 | 0.152 | 1.127 | 0.260 |
| [+2°C] x To | 0.239 | 0.152 | 1.575 | 0.115 |
| HW x To | 0.064 | 0.152 | 0.421 | 0.674 |
| [+2°C+HW] x To | 0.188 | 0.152 | 1.240 | 0.215 |
| [+2°C] x Sp | 0.211 | 0.152 | 1.388 | 0.165 |
| HW x Sp | 0.117 | 0.152 | 0.767 | 0.443 |
| [+2°C+HW] x Sp | 0.217 | 0.152 | 1.428 | 0.153 |
| [+2°C] x Sc | 0.282 | 0.152 | 1.858 | 0.063 |
| HW x Sc | 0.185 | 0.152 | 1.217 | 0.224 |
| [+2°C+HW] x Sc | 0.176 | 0.152 | 1.157 | 0.247 |
| D x 1M | 0.015 | 0.132 | 0.115 | 0.908 |
| D x 4M | 0.040 | 0.132 | 0.304 | 0.761 |
| [+2°C] x 1M | 0.177 | 0.132 | 1.344 | 0.179 |
| HW x 1M | 0.064 | 0.132 | 0.490 | 0.624 |
| [+2°C+HW] x 1M | 0.052 | 0.132 | 0.393 | 0.694 |
| **[+2°C] x 4M** | **0.320** | **0.132** | **2.433** | **0.015** |
| HW x 4M | 0.102 | 0.132 | 0.771 | 0.440 |
| [+2°C+HW] x 4M | 0.116 | 0.132 | 0.878 | 0.380 |
| Be x 1M | -0.106 | 0.132 | -0.803 | 0.422 |
| Tp x 1M | -0.060 | 0.132 | -0.454 | 0.650 |
| Lc x 1M | 0.016 | 0.132 | 0.124 | 0.901 |
| Cj x 1M | -0.064 | 0.132 | -0.490 | 0.624 |
| To x 1M | 0.085 | 0.132 | 0.649 | 0.517 |
| Sp x 1M | 0.021 | 0.132 | 0.156 | 0.876 |
| Sc x 1M | 0.079 | 0.132 | 0.601 | 0.548 |
| Be x 4M | 0.021 | 0.132 | 0.161 | 0.872 |
| Tp x 4M | -0.200 | 0.134 | -1.489 | 0.136 |
| Lc x 4M | 0.094 | 0.132 | 0.717 | 0.473 |
| Cj x 4M | -0.079 | 0.132 | -0.603 | 0.546 |
| To x 4M | 0.063 | 0.132 | 0.478 | 0.633 |
| Sp x 4M | -0.061 | 0.133 | -0.457 | 0.647 |
| Sc x 4M | 0.105 | 0.132 | 0.796 | 0.426 |
| D x [+2°C] x Be | 0.173 | 0.215 | 0.804 | 0.421 |
| D x HW x Be | 0.355 | 0.215 | 1.651 | 0.099 |
| D x [+2°C+HW] x Be | 0.097 | 0.215 | 0.453 | 0.650 |
| D x [+2°C] x Tp | 0.108 | 0.215 | 0.501 | 0.617 |
| D x HW x Tp | 0.028 | 0.215 | 0.132 | 0.895 |
| D x [+2°C+HW] x Tp | -0.152 | 0.218 | -0.695 | 0.487 |
| D x [+2°C] x Lc | -0.118 | 0.215 | -0.549 | 0.583 |
| D x HW x Lc | -0.133 | 0.215 | -0.619 | 0.536 |
| D x [+2°C+HW] x Lc | -0.234 | 0.215 | -1.090 | 0.276 |
| D x [+2°C] x Cj | -0.056 | 0.215 | -0.263 | 0.793 |
| D x HW x Cj | 0.014 | 0.215 | 0.064 | 0.949 |
| D x [+2°C+HW] x Cj | -0.193 | 0.215 | -0.897 | 0.370 |
| D x [+2°C] x To | -0.133 | 0.215 | -0.620 | 0.535 |
| D x HW x To | 0.063 | 0.215 | 0.293 | 0.770 |
| D x [+2°C+HW] x To | -0.153 | 0.215 | -0.711 | 0.477 |
| D x [+2°C] x Sp | 0.006 | 0.215 | 0.027 | 0.979 |
| D x HW x Sp | 0.011 | 0.215 | 0.054 | 0.957 |
| D x [+2°C+HW] x Sp | -0.234 | 0.215 | -1.088 | 0.277 |
| D x [+2°C] x Sc | -0.074 | 0.215 | -0.346 | 0.729 |
| D x HW x Sc | 0.033 | 0.215 | 0.154 | 0.878 |
| D x [+2°C+HW] x Sc | -0.144 | 0.215 | -0.670 | 0.503 |
| D x [+2°C] x 1M | -0.083 | 0.186 | -0.446 | 0.656 |
| D x HW x 1M | 0.198 | 0.186 | 1.062 | 0.288 |
| D x [+2°C+HW] x 1M | 0.025 | 0.186 | 0.136 | 0.892 |
| D x [+2°C] x 4M | -0.187 | 0.186 | -1.004 | 0.316 |
| D x HW x 4M | 0.102 | 0.186 | 0.549 | 0.583 |
| D x [+2°C+HW] x 4M | -0.086 | 0.186 | -0.462 | 0.644 |
| D x Be x 1M | 0.238 | 0.186 | 1.281 | 0.200 |
| D x Tp x 1M | -0.094 | 0.186 | -0.505 | 0.614 |
| D x Lc x 1M | 0.003 | 0.186 | 0.017 | 0.987 |
| D x Cj x 1M | 0.116 | 0.187 | 0.619 | 0.536 |
| D x To x 1M | 0.114 | 0.186 | 0.612 | 0.541 |
| D x Sp x 1M | 0.056 | 0.186 | 0.301 | 0.763 |
| D x Sc x 1M | 0.087 | 0.186 | 0.466 | 0.641 |
| D x Be x 4M | 0.285 | 0.186 | 1.529 | 0.126 |
| D x Tp x 4M | 0.097 | 0.188 | 0.515 | 0.607 |
| D x Lc x 4M | -0.059 | 0.186 | -0.318 | 0.751 |
| D x Cj x 4M | -0.123 | 0.187 | -0.658 | 0.511 |
| D x To x 4M | -0.085 | 0.186 | -0.457 | 0.648 |
| D x Sp x 4M | 0.068 | 0.187 | 0.363 | 0.717 |
| D x Sc x 4M | -0.060 | 0.186 | -0.324 | 0.746 |
| [+2°C] x Be x 1M | -0.167 | 0.186 | -0.898 | 0.369 |
| HW x Be x 1M | 0.093 | 0.186 | 0.500 | 0.617 |
| [+2°C+HW] x Be x 1M | -0.212 | 0.186 | -1.139 | 0.255 |
| [+2°C] x Tp x 1M | -0.038 | 0.186 | -0.206 | 0.837 |
| HW x Tp x 1M | -0.090 | 0.186 | -0.486 | 0.627 |
| [+2°C+HW] x Tp x 1M | 0.044 | 0.190 | 0.233 | 0.816 |
| [+2°C] x Lc x 1M | -0.067 | 0.186 | -0.362 | 0.718 |
| HW x Lc x 1M | -0.209 | 0.186 | -1.124 | 0.261 |
| [+2°C+HW] x Lc x 1M | -0.141 | 0.186 | -0.759 | 0.448 |
| [+2°C] x Cj x 1M | -0.059 | 0.186 | -0.319 | 0.750 |
| HW x Cj x 1M | -0.071 | 0.187 | -0.381 | 0.703 |
| [+2°C+HW] x Cj x 1M | 0.014 | 0.186 | 0.073 | 0.942 |
| [+2°C] x To x 1M | -0.136 | 0.186 | -0.729 | 0.466 |
| HW x To x 1M | 0.071 | 0.186 | 0.380 | 0.704 |
| [+2°C+HW] x To x 1M | 0.035 | 0.186 | 0.189 | 0.850 |
| [+2°C] x Sp x 1M | -0.267 | 0.186 | -1.432 | 0.152 |
| HW x Sp x 1M | -0.039 | 0.186 | -0.212 | 0.832 |
| [+2°C+HW] x Sp x 1M | -0.142 | 0.186 | -0.763 | 0.446 |
| [+2°C] x Sc x 1M | -0.256 | 0.186 | -1.375 | 0.169 |
| HW x Sc x 1M | -0.074 | 0.186 | -0.397 | 0.691 |
| [+2°C+HW] x Sc x 1M | 0.020 | 0.186 | 0.107 | 0.915 |
| [+2°C] x Be x 4M | -0.204 | 0.186 | -1.096 | 0.273 |
| HW x Be x 4M | 0.170 | 0.186 | 0.914 | 0.361 |
| [+2°C+HW] x Be x 4M | -0.029 | 0.186 | -0.158 | 0.874 |
| [+2°C] x Tp x 4M | -0.036 | 0.188 | -0.189 | 0.850 |
| HW x Tp x 4M | -0.045 | 0.190 | -0.236 | 0.813 |
| [+2°C+HW] x Tp x 4M | 0.002 | 0.193 | 0.011 | 0.992 |
| [+2°C] x Lc x 4M | -0.250 | 0.186 | -1.343 | 0.179 |
| HW x Lc x 4M | -0.125 | 0.186 | -0.670 | 0.503 |
| [+2°C+HW] x Lc x 4M | -0.127 | 0.186 | -0.681 | 0.496 |
| [+2°C] x Cj x 4M | -0.240 | 0.186 | -1.290 | 0.197 |
| HW x Cj x 4M | -0.110 | 0.188 | -0.586 | 0.558 |
| [+2°C+HW] x Cj x 4M | -0.083 | 0.187 | -0.444 | 0.657 |
| [+2°C] x To x 4M | -0.270 | 0.186 | -1.448 | 0.148 |
| HW x To x 4M | 0.009 | 0.186 | 0.046 | 0.963 |
| [+2°C+HW] x To x 4M | -0.075 | 0.186 | -0.402 | 0.688 |
| [+2°C] x Sp x 4M | -0.284 | 0.187 | -1.516 | 0.129 |
| HW x Sp x 4M | -0.105 | 0.187 | -0.562 | 0.574 |
| [+2°C+HW] x Sp x 4M | -0.082 | 0.187 | -0.438 | 0.662 |
| **[+2°C] x Sc x 4M** | **-0.383** | **0.186** | **-2.058** | **0.040** |
| HW x Sc x 4M | -0.195 | 0.186 | -1.047 | 0.295 |
| [+2°C+HW] x Sc x 4M | -0.122 | 0.186 | -0.656 | 0.512 |
| D x [+2°C] x Be x 1M | 0.008 | 0.263 | 0.031 | 0.976 |
| D x HW x Be x 1M | -0.482 | 0.263 | -1.830 | 0.067 |
| D x [+2°C+HW] x Be x 1M | -0.008 | 0.263 | -0.029 | 0.977 |
| D x [+2°C] x Tp x 1M | 0.129 | 0.263 | 0.491 | 0.623 |
| D x HW x Tp x 1M | -0.088 | 0.263 | -0.336 | 0.737 |
| D x [+2°C+HW] x Tp x 1M | 0.024 | 0.267 | 0.088 | 0.930 |
| D x [+2°C] x Lc x 1M | 0.037 | 0.263 | 0.139 | 0.889 |
| D x HW x Lc x 1M | 0.038 | 0.263 | 0.146 | 0.884 |
| D x [+2°C+HW] x Lc x 1M | -0.057 | 0.263 | -0.218 | 0.827 |
| D x [+2°C] x Cj x 1M | -0.048 | 0.264 | -0.182 | 0.856 |
| D x HW x Cj x 1M | -0.156 | 0.265 | -0.588 | 0.556 |
| D x [+2°C+HW] x Cj x 1M | 0.002 | 0.264 | 0.009 | 0.993 |
| D x [+2°C] x To x 1M | 0.053 | 0.263 | 0.203 | 0.840 |
| D x HW x To x 1M | -0.317 | 0.263 | -1.204 | 0.228 |
| D x [+2°C+HW] x To x 1M | -0.151 | 0.263 | -0.575 | 0.566 |
| D x [+2°C] x Sp x 1M | 0.163 | 0.263 | 0.621 | 0.535 |
| D x HW x Sp x 1M | -0.198 | 0.263 | -0.754 | 0.451 |
| D x [+2°C+HW] x Sp x 1M | 0.101 | 0.263 | 0.385 | 0.700 |
| D x [+2°C] x Sc x 1M | 0.147 | 0.263 | 0.560 | 0.575 |
| D x HW x Sc x 1M | -0.155 | 0.263 | -0.590 | 0.555 |
| D x [+2°C+HW] x Sc x 1M | -0.100 | 0.263 | -0.380 | 0.704 |
| D x [+2°C] x Be x 4M | -0.080 | 0.263 | -0.304 | 0.761 |
| **D x HW x Be x 4M** | **-0.550** | **0.263** | **-2.088** | **0.037** |
| D x [+2°C+HW] x Be x 4M | -0.149 | 0.263 | -0.565 | 0.572 |
| D x [+2°C] x Tp x 4M | -0.116 | 0.265 | -0.439 | 0.660 |
| D x HW x Tp x 4M | -0.162 | 0.267 | -0.608 | 0.543 |
| D x [+2°C+HW] x Tp x 4M | -0.034 | 0.269 | -0.126 | 0.900 |
| D x [+2°C] x Lc x 4M | 0.234 | 0.263 | 0.888 | 0.375 |
| D x HW x Lc x 4M | -0.021 | 0.263 | -0.078 | 0.938 |
| D x [+2°C+HW] x Lc x 4M | -0.026 | 0.263 | -0.100 | 0.920 |
| D x [+2°C] x Cj x 4M | 0.234 | 0.264 | 0.884 | 0.377 |
| D x HW x Cj x 4M | 0.089 | 0.267 | 0.334 | 0.738 |
| D x [+2°C+HW] x Cj x 4M | 0.164 | 0.264 | 0.620 | 0.535 |
| D x [+2°C] x To x 4M | 0.320 | 0.263 | 1.216 | 0.224 |
| D x HW x To x 4M | -0.085 | 0.263 | -0.321 | 0.748 |
| D x [+2°C+HW] x To x 4M | 0.106 | 0.263 | 0.404 | 0.686 |
| D x [+2°C] x Sp x 4M | 0.204 | 0.264 | 0.774 | 0.439 |
| D x HW x Sp x 4M | -0.086 | 0.264 | -0.325 | 0.745 |
| D x [+2°C+HW] x Sp x 4M | 0.080 | 0.264 | 0.305 | 0.760 |
| D x [+2°C] x Sc x 4M | 0.181 | 0.263 | 0.686 | 0.493 |
| D x HW x Sc x 4M | -0.075 | 0.263 | -0.287 | 0.774 |
| D x [+2°C+HW] x Sc x 4M | 0.031 | 0.263 | 0.120 | 0.905 |

**Table S10b:** Output of the general model of the chlorophyll content responses to the treatment and season according to the plants' species *Prunella vulgaris*. The model uses the above-ground biomass as the response variable, and as explanatory variables, the interactive effect of Drought, Temperature, harvest time points. Significant estimates are highlighted in bold. Abbreviations: D: Water treatment drought, [+2°C]: Ambient temperature plus constant 2°C warming, HW: Ambient temperature plus heatwave warming events (+10°C), [+2°C+HW]: Ambient temperature plus constant 2°C and heatwave warming. 1M: collection point at 1-month recovery after the end of drought (peak of the growing season), 4M: Data collection point at 4-months recovery after the end of drought (end of the growing season).

| Predictor | Estimate | SE | Z | p-value |
| --- | --- | --- | --- | --- |
| (Intercept) | 10.035 | 0.082 | 122.862 | <0.001 |
| D | 0.150 | 0.109 | 1.371 | 0.170 |
| [+2°C] | 0.046 | 0.109 | 0.419 | 0.675 |
| HW | 0.084 | 0.116 | 0.729 | 0.466 |
| [+2°C+HW] | 0.043 | 0.109 | 0.391 | 0.696 |
| 1M | 0.098 | 0.100 | 0.985 | 0.325 |
| 4M | 0.540 | 0.098 | 5.500 | <0.001 |
| D x [+2°C] | -0.037 | 0.167 | -0.219 | 0.827 |
| D x HW | -0.215 | 0.171 | -1.257 | 0.209 |
| D x [+2°C+HW] | -0.238 | 0.219 | -1.090 | 0.276 |
| D x 1M | -0.123 | 0.135 | -0.908 | 0.364 |
| D x 4M | -0.169 | 0.136 | -1.236 | 0.216 |
| [+2°C] x 1M | -0.000 | 0.135 | -0.003 | 0.997 |
| HW x 1M | -0.021 | 0.142 | -0.150 | 0.881 |
| [+2°C+HW] x 1M | 0.057 | 0.136 | 0.416 | 0.677 |
| [+2°C] x 4M | -0.083 | 0.135 | -0.613 | 0.540 |
| HW x 4M | -0.107 | 0.141 | -0.761 | 0.446 |
| [+2°C+HW] x 4M | -0.115 | 0.136 | -0.840 | 0.401 |
| D x [+2°C] x 1M | -0.046 | 0.212 | -0.218 | 0.827 |
| D x HW x 1M | 0.189 | 0.210 | 0.902 | 0.367 |
| D x [+2°C+HW] x 1M | 0.006 | 0.296 | 0.022 | 0.983 |
| D x [+2°C] x 4M | 0.091 | 0.209 | 0.437 | 0.662 |
| D x HW x 4M | 0.290 | 0.221 | 1.309 | 0.191 |
| D x [+2°C+HW] x 4M | 0.353 | 0.270 | 1.311 | 0.190 |

**Table S11a:** Output of the mixed-effects models testing specific stomatal conductance responses to treatments and season (species-specific recovery model). The model included drought, temperature, harvest time point, and plant species as explanatory variables. Due to model-fitting issues, *Prunella vulgaris* was analysed separately in an independent model (see table S11b). Significant estimates are highlighted in bold. Abbreviations: D: Water treatment drought, [+2°C]: Ambient temperature plus constant 2°C warming, HW: Ambient temperature plus heatwave warming events (+10°C), [+2°C+HW]: Ambient temperature plus constant 2°C and heatwave warming. 1M: collection point at 1-month recovery after the end of drought (peak of the growing season), 4M: Data collection point at 4-months recovery after the end of drought (end of the growing season). *Bromus erectus*: Be, *Trifolium pratense*: Tp, *Lotus corniculatus*: Lc, *Taraxacum officinale*: To, *Centaurea jacea*: Cj, *Salvia pratense*: Sp, *Solidago canadensis*: Sc. The intercept is the fast-growing species *Holcus lanatus*.

| Predictor | Estimate | SE | Z | p-value |
| --- | --- | --- | --- | --- |
| (Intercept) | 5.517 | 0.236 | 23.371 | <0.001 |
| D | 0.070 | 0.327 | 0.214 | 0.830 |
| [+2°C] | -0.053 | 0.327 | -0.163 | 0.870 |
| HW | -0.194 | 0.327 | -0.593 | 0.553 |
| [+2°C+HW] | -0.497 | 0.327 | -1.518 | 0.129 |
| Be | 0.401 | 0.327 | 1.226 | 0.220 |
| Tp | 0.184 | 0.327 | 0.563 | 0.574 |
| Lc | 0.389 | 0.327 | 1.189 | 0.235 |
| Cj | 0.319 | 0.327 | 0.975 | 0.329 |
| To | -0.095 | 0.327 | -0.291 | 0.771 |
| Sp | -1.325 | 0.327 | -4.048 | <0.001 |
| **Sc** | **-0.750** | **0.327** | **-2.291** | **0.022** |
| **1M** | **-0.618** | **0.284** | **-2.178** | **0.029** |
| 4M | 0.078 | 0.284 | 0.274 | 0.784 |
| D x [+2°C] | -0.263 | 0.463 | -0.568 | 0.570 |
| D x HW | 0.105 | 0.463 | 0.227 | 0.820 |
| D x [+2°C+HW] | 0.249 | 0.463 | 0.538 | 0.591 |
| D x Be | -0.488 | 0.463 | -1.054 | 0.292 |
| D x Tp | -0.493 | 0.463 | -1.065 | 0.287 |
| D x Lc | -0.344 | 0.463 | -0.743 | 0.458 |
| D x Cj | -0.336 | 0.463 | -0.726 | 0.468 |
| D x To | -0.275 | 0.463 | -0.595 | 0.552 |
| D x Sp | -0.172 | 0.463 | -0.373 | 0.710 |
| D x Sc | -0.553 | 0.463 | -1.195 | 0.232 |
| [+2°C] x Be | -0.226 | 0.463 | -0.489 | 0.625 |
| HW x Be | -0.540 | 0.463 | -1.167 | 0.243 |
| [+2°C+HW] x Be | 0.286 | 0.463 | 0.617 | 0.537 |
| [+2°C] x Tp | 0.025 | 0.463 | 0.053 | 0.958 |
| HW x Tp | 0.192 | 0.463 | 0.415 | 0.678 |
| [+2°C+HW] x Tp | 0.565 | 0.477 | 1.185 | 0.236 |
| [+2°C] x Lc | -0.077 | 0.463 | -0.167 | 0.868 |
| HW x Lc | 0.193 | 0.463 | 0.418 | 0.676 |
| [+2°C+HW] x Lc | 0.310 | 0.463 | 0.670 | 0.503 |
| [+2°C] x Cj | -0.074 | 0.463 | -0.161 | 0.872 |
| HW x Cj | -0.164 | 0.463 | -0.354 | 0.723 |
| [+2°C+HW] x Cj | 0.631 | 0.463 | 1.363 | 0.173 |
| [+2°C] x To | -0.067 | 0.463 | -0.144 | 0.885 |
| HW x To | 0.269 | 0.463 | 0.581 | 0.561 |
| [+2°C+HW] x To | 0.567 | 0.463 | 1.225 | 0.220 |
| [+2°C] x Sp | -0.280 | 0.463 | -0.605 | 0.545 |
| HW x Sp | 0.047 | 0.463 | 0.101 | 0.920 |
| [+2°C+HW] x Sp | 0.470 | 0.463 | 1.015 | 0.310 |
| [+2°C] x Sc | 0.161 | 0.463 | 0.349 | 0.727 |
| HW x Sc | 0.017 | 0.463 | 0.038 | 0.970 |
| [+2°C+HW] x Sc | 0.615 | 0.463 | 1.330 | 0.184 |
| D x 1M | -0.061 | 0.401 | -0.151 | 0.880 |
| D x 4M | -0.058 | 0.401 | -0.145 | 0.885 |
| [+2°C] x 1M | -0.313 | 0.401 | -0.780 | 0.435 |
| HW x 1M | 0.029 | 0.401 | 0.073 | 0.942 |
| [+2°C+HW] x 1M | 0.262 | 0.401 | 0.654 | 0.513 |
| [+2°C] x 4M | 0.094 | 0.401 | 0.234 | 0.815 |
| HW x 4M | 0.223 | 0.401 | 0.556 | 0.578 |
| [+2°C+HW] x 4M | 0.243 | 0.405 | 0.600 | 0.549 |
| Be x 1M | -0.405 | 0.401 | -1.011 | 0.312 |
| Tp x 1M | 0.381 | 0.405 | 0.942 | 0.346 |
| Lc x 1M | -0.171 | 0.401 | -0.426 | 0.670 |
| Cj x 1M | 0.431 | 0.405 | 1.065 | 0.287 |
| To x 1M | 0.321 | 0.401 | 0.802 | 0.423 |
| **Sp x 1M** | **0.923** | **0.423** | **2.183** | **0.029** |
| Sc x 1M | 0.725 | 0.401 | 1.809 | 0.070 |
| Be x 4M | -0.477 | 0.401 | -1.190 | 0.234 |
| Tp x 4M | -0.183 | 0.423 | -0.433 | 0.665 |
| Lc x 4M | -0.505 | 0.401 | -1.260 | 0.208 |
| Cj x 4M | -0.483 | 0.401 | -1.204 | 0.228 |
| To x 4M | -0.164 | 0.401 | -0.408 | 0.683 |
| **Sp x 4M** | **0.840** | **0.405** | **2.077** | **0.038** |
| Sc x 4M | 0.156 | 0.401 | 0.388 | 0.698 |
| D x [+2°C] x Be | 0.621 | 0.655 | 0.948 | 0.343 |
| D x HW x Be | 0.658 | 0.655 | 1.005 | 0.315 |
| D x [+2°C+HW] x Be | 0.040 | 0.655 | 0.062 | 0.951 |
| D x [+2°C] x Tp | 0.419 | 0.655 | 0.641 | 0.522 |
| D x HW x Tp | -0.114 | 0.655 | -0.175 | 0.861 |
| D x [+2°C+HW] x Tp | -0.337 | 0.665 | -0.507 | 0.612 |
| D x [+2°C] x Lc | 0.130 | 0.655 | 0.199 | 0.842 |
| D x HW x Lc | 0.047 | 0.655 | 0.072 | 0.942 |
| D x [+2°C+HW] x Lc | -0.563 | 0.655 | -0.860 | 0.390 |
| D x [+2°C] x Cj | 0.210 | 0.655 | 0.321 | 0.748 |
| D x HW x Cj | -0.045 | 0.655 | -0.068 | 0.946 |
| D x [+2°C+HW] x Cj | -0.679 | 0.655 | -1.038 | 0.299 |
| D x [+2°C] x To | 0.276 | 0.655 | 0.421 | 0.674 |
| D x HW x To | -0.071 | 0.655 | -0.108 | 0.914 |
| D x [+2°C+HW] x To | -0.316 | 0.655 | -0.483 | 0.629 |
| D x [+2°C] x Sp | 0.570 | 0.655 | 0.871 | 0.384 |
| D x HW x Sp | -0.188 | 0.655 | -0.288 | 0.773 |
| D x [+2°C+HW] x Sp | -0.579 | 0.655 | -0.885 | 0.376 |
| D x [+2°C] x Sc | 0.004 | 0.655 | 0.006 | 0.996 |
| D x HW x Sc | -0.126 | 0.655 | -0.193 | 0.847 |
| D x [+2°C+HW] x Sc | -0.728 | 0.655 | -1.112 | 0.266 |
| D x [+2°C] x 1M | 0.524 | 0.567 | 0.924 | 0.355 |
| D x HW x 1M | 0.100 | 0.567 | 0.177 | 0.859 |
| D x [+2°C+HW] x 1M | 0.118 | 0.567 | 0.208 | 0.835 |
| D x [+2°C] x 4M | 0.325 | 0.567 | 0.573 | 0.566 |
| D x HW x 4M | -0.064 | 0.567 | -0.112 | 0.911 |
| D x [+2°C+HW] x 4M | 0.153 | 0.569 | 0.269 | 0.788 |
| D x Be x 1M | 0.197 | 0.567 | 0.348 | 0.728 |
| D x Tp x 1M | 0.622 | 0.572 | 1.088 | 0.277 |
| D x Lc x 1M | 0.116 | 0.567 | 0.205 | 0.838 |
| D x Cj x 1M | 0.067 | 0.572 | 0.117 | 0.907 |
| D x To x 1M | 0.406 | 0.567 | 0.716 | 0.474 |
| D x Sp x 1M | 1.050 | 0.598 | 1.757 | 0.079 |
| D x Sc x 1M | 0.700 | 0.567 | 1.235 | 0.217 |
| D x Be x 4M | 0.120 | 0.567 | 0.212 | 0.832 |
| D x Tp x 4M | 0.447 | 0.582 | 0.768 | 0.442 |
| D x Lc x 4M | 0.428 | 0.567 | 0.756 | 0.450 |
| D x Cj x 4M | 0.257 | 0.569 | 0.452 | 0.651 |
| D x To x 4M | 0.460 | 0.567 | 0.812 | 0.417 |
| D x Sp x 4M | 0.360 | 0.569 | 0.632 | 0.527 |
| D x Sc x 4M | 0.393 | 0.567 | 0.693 | 0.489 |
| [+2°C] x Be x 1M | 0.585 | 0.567 | 1.031 | 0.302 |
| HW x Be x 1M | 0.135 | 0.567 | 0.237 | 0.812 |
| [+2°C+HW] x Be x 1M | -0.443 | 0.567 | -0.781 | 0.435 |
| [+2°C] x Tp x 1M | 0.215 | 0.572 | 0.375 | 0.708 |
| HW x Tp x 1M | -0.375 | 0.569 | -0.659 | 0.510 |
| [+2°C+HW] x Tp x 1M | -0.522 | 0.584 | -0.895 | 0.371 |
| [+2°C] x Lc x 1M | 0.512 | 0.567 | 0.903 | 0.366 |
| HW x Lc x 1M | -0.344 | 0.567 | -0.607 | 0.544 |
| [+2°C+HW] x Lc x 1M | -0.477 | 0.567 | -0.842 | 0.400 |
| [+2°C] x Cj x 1M | 0.308 | 0.575 | 0.535 | 0.593 |
| HW x Cj x 1M | -0.091 | 0.572 | -0.159 | 0.874 |
| [+2°C+HW] x Cj x 1M | -0.701 | 0.569 | -1.231 | 0.218 |
| [+2°C] x To x 1M | 0.785 | 0.567 | 1.385 | 0.166 |
| HW x To x 1M | 0.001 | 0.567 | 0.001 | 0.999 |
| [+2°C+HW] x To x 1M | -0.103 | 0.567 | -0.181 | 0.856 |
| **[+2°C] x Sp x 1M** | **1.350** | **0.588** | **2.295** | **0.022** |
| HW x Sp x 1M | 0.442 | 0.592 | 0.746 | 0.455 |
| [+2°C+HW] x Sp x 1M | 0.380 | 0.598 | 0.637 | 0.524 |
| [+2°C] x Sc x 1M | 0.177 | 0.569 | 0.311 | 0.756 |
| HW x Sc x 1M | 0.217 | 0.567 | 0.383 | 0.702 |
| [+2°C+HW] x Sc x 1M | -0.113 | 0.567 | -0.199 | 0.842 |
| [+2°C] x Be x 4M | -0.379 | 0.567 | -0.668 | 0.504 |
| HW x Be x 4M | 0.299 | 0.567 | 0.528 | 0.597 |
| [+2°C+HW] x Be x 4M | -0.347 | 0.569 | -0.609 | 0.543 |
| [+2°C] x Tp x 4M | -0.120 | 0.582 | -0.206 | 0.837 |
| HW x Tp x 4M | -0.148 | 0.585 | -0.253 | 0.801 |
| [+2°C+HW] x Tp x 4M | -0.132 | 0.599 | -0.221 | 0.825 |
| [+2°C] x Lc x 4M | 0.001 | 0.567 | 0.001 | 0.999 |
| HW x Lc x 4M | -0.197 | 0.567 | -0.347 | 0.729 |
| [+2°C+HW] x Lc x 4M | -0.056 | 0.569 | -0.099 | 0.921 |
| [+2°C] x Cj x 4M | -0.073 | 0.567 | -0.129 | 0.898 |
| HW x Cj x 4M | 0.122 | 0.573 | 0.212 | 0.832 |
| [+2°C+HW] x Cj x 4M | -0.414 | 0.569 | -0.727 | 0.467 |
| [+2°C] x To x 4M | 0.118 | 0.567 | 0.209 | 0.835 |
| HW x To x 4M | -0.070 | 0.567 | -0.124 | 0.901 |
| [+2°C+HW] x To x 4M | -0.220 | 0.569 | -0.387 | 0.699 |
| [+2°C] x Sp x 4M | 0.598 | 0.569 | 1.051 | 0.293 |
| HW x Sp x 4M | 0.041 | 0.569 | 0.072 | 0.943 |
| [+2°C+HW] x Sp x 4M | -0.209 | 0.572 | -0.365 | 0.715 |
| [+2°C] x Sc x 4M | -0.265 | 0.567 | -0.467 | 0.640 |
| HW x Sc x 4M | 0.184 | 0.567 | 0.324 | 0.746 |
| [+2°C+HW] x Sc x 4M | -0.336 | 0.569 | -0.590 | 0.555 |
| D x [+2°C] x Be x 1M | -0.555 | 0.802 | -0.692 | 0.489 |
| D x HW x Be x 1M | -0.330 | 0.802 | -0.411 | 0.681 |
| D x [+2°C+HW] x Be x 1M | 0.343 | 0.802 | 0.428 | 0.668 |
| D x [+2°C] x Tp x 1M | -0.467 | 0.814 | -0.573 | 0.567 |
| D x HW x Tp x 1M | 0.256 | 0.816 | 0.313 | 0.754 |
| D x [+2°C+HW] x Tp x 1M | -0.141 | 0.820 | -0.172 | 0.864 |
| D x [+2°C] x Lc x 1M | -0.169 | 0.802 | -0.211 | 0.833 |
| D x HW x Lc x 1M | 0.245 | 0.802 | 0.305 | 0.760 |
| D x [+2°C+HW] x Lc x 1M | 0.974 | 0.802 | 1.215 | 0.224 |
| D x [+2°C] x Cj x 1M | -0.245 | 0.811 | -0.302 | 0.762 |
| D x HW x Cj x 1M | 0.069 | 0.809 | 0.085 | 0.933 |
| D x [+2°C+HW] x Cj x 1M | 0.862 | 0.816 | 1.056 | 0.291 |
| D x [+2°C] x To x 1M | -0.976 | 0.802 | -1.218 | 0.223 |
| D x HW x To x 1M | -0.031 | 0.802 | -0.039 | 0.969 |
| D x [+2°C+HW] x To x 1M | -0.266 | 0.804 | -0.331 | 0.741 |
| **D x [+2°C] x Sp x 1M** | **-1.900** | **0.838** | **-2.266** | **0.023** |
| D x HW x Sp x 1M | -0.165 | 0.847 | -0.195 | 0.845 |
| D x [+2°C+HW] x Sp x 1M | -0.201 | 0.841 | -0.239 | 0.811 |
| D x [+2°C] x Sc x 1M | -0.342 | 0.804 | -0.425 | 0.671 |
| D x HW x Sc x 1M | -0.052 | 0.802 | -0.065 | 0.949 |
| D x [+2°C+HW] x Sc x 1M | -0.269 | 0.804 | -0.335 | 0.738 |
| D x [+2°C] x Be x 4M | -0.094 | 0.802 | -0.117 | 0.907 |
| D x HW x Be x 4M | -0.342 | 0.802 | -0.427 | 0.669 |
| D x [+2°C+HW] x Be x 4M | 0.167 | 0.804 | 0.208 | 0.835 |
| D x [+2°C] x Tp x 4M | -0.188 | 0.813 | -0.232 | 0.817 |
| D x HW x Tp x 4M | 0.229 | 0.816 | 0.280 | 0.779 |
| D x [+2°C+HW] x Tp x 4M | -0.149 | 0.829 | -0.179 | 0.858 |
| D x [+2°C] x Lc x 4M | -0.276 | 0.802 | -0.344 | 0.731 |
| D x HW x Lc x 4M | -0.177 | 0.802 | -0.221 | 0.825 |
| D x [+2°C+HW] x Lc x 4M | 0.016 | 0.804 | 0.019 | 0.984 |
| D x [+2°C] x Cj x 4M | 0.026 | 0.805 | 0.032 | 0.974 |
| D x HW x Cj x 4M | 0.337 | 0.812 | 0.415 | 0.678 |
| D x [+2°C+HW] x Cj x 4M | 0.595 | 0.805 | 0.739 | 0.460 |
| D x [+2°C] x To x 4M | -0.261 | 0.802 | -0.326 | 0.744 |
| D x HW x To x 4M | 0.036 | 0.802 | 0.045 | 0.964 |
| D x [+2°C+HW] x To x 4M | -0.091 | 0.804 | -0.114 | 0.910 |
| D x [+2°C] x Sp x 4M | -1.155 | 0.804 | -1.438 | 0.150 |
| D x HW x Sp x 4M | 0.121 | 0.804 | 0.151 | 0.880 |
| D x [+2°C+HW] x Sp x 4M | 0.170 | 0.805 | 0.212 | 0.832 |
| D x [+2°C] x Sc x 4M | 0.225 | 0.802 | 0.281 | 0.779 |
| D x HW x Sc x 4M | 0.081 | 0.802 | 0.102 | 0.919 |
| D x [+2°C+HW] x Sc x 4M | 0.216 | 0.804 | 0.269 | 0.788 |

**Table S11b:** Output of the general model of the stomatal conductance responses to the treatment and season according to the plants' species *Prunella vulgaris*. The model uses the above-ground biomass as the response variable, and as explanatory variables, the interactive effect of Drought, Temperature, harvest time points. Significant estimates are highlighted in bold. Abbreviations: D: Water treatment drought, [+2°C]: Ambient temperature plus constant 2°C warming, HW: Ambient temperature plus heatwave warming events (+10°C), [+2°C+HW]: Ambient temperature plus constant 2°C and heatwave warming. 1M: collection point at 1-month recovery after the end of drought (peak of the growing season), 4M: Data collection point at 4-months recovery after the end of drought (end of the growing season).

| Predictor | Estimate | SE | Z | p-value |
| --- | --- | --- | --- | --- |
| (Intercept) | 8.918 | 0.223 | 39.927 | <0.001 |
| D | 0.006 | 0.314 | 0.021 | 0.984 |
| [+2°C] | 0.235 | 0.300 | 0.784 | 0.433 |
| HW | 0.012 | 0.362 | 0.032 | 0.974 |
| [+2°C+HW] | -0.072 | 0.320 | -0.224 | 0.823 |
| 1M | -0.029 | 0.317 | -0.092 | 0.926 |
| **4M** | **0.758** | **0.252** | **3.005** | **0.003** |
| D x [+2°C] | -0.525 | 0.494 | -1.063 | 0.288 |
| D x HW | -0.303 | 0.581 | -0.521 | 0.602 |
| D x [+2°C+HW] | -0.125 | 0.665 | -0.188 | 0.851 |
| **D x 1M** | **0.896** | **0.415** | **2.158** | **0.031** |
| D x 4M | -0.494 | 0.364 | -1.359 | 0.174 |
| [+2°C] x 1M | -0.393 | 0.446 | -0.882 | 0.378 |
| HW x 1M | -0.262 | 0.510 | -0.514 | 0.607 |
| [+2°C+HW] x 1M | 0.382 | 0.449 | 0.850 | 0.395 |
| [+2°C] x 4M | -0.506 | 0.346 | -1.459 | 0.144 |
| HW x 4M | -0.412 | 0.404 | -1.019 | 0.308 |
| [+2°C+HW] x 4M | -0.274 | 0.368 | -0.744 | 0.457 |
| D x [+2°C] x 1M | 0.037 | 0.666 | 0.056 | 0.955 |
| D x HW x 1M | 0.397 | 0.730 | 0.544 | 0.586 |
| D x [+2°C+HW] x 1M | NA | NA | NA | NA |
| **D x [+2°C] x 4M** | **1.259** | **0.565** | **2.227** | **0.026** |
| D x HW x 4M | 0.725 | 0.673 | 1.076 | 0.282 |
| D x [+2°C+HW] x 4M | 0.791 | 0.754 | 1.049 | 0.294 |
